# Decoding scalp health and microbiome dysbiosis in dandruff

**DOI:** 10.1101/2024.05.02.592279

**Authors:** Viduthalai Rasheedkhan Regina, Tarun Chopra, Kwok Weihao, Sreelakshmi Cheruvalli, Ang Sabrina, Hashmath Fatimah Binte Jamal Mohamed, Kesava Priyan Ramasamy, Kay Sarah, Chong Yik Yan, Eganathan Kaliyamoorthi, Rohan Williams, Liu Xianghui, Vedula Krishna, Nasrine Bourokba, Anjali Jhingan, Steven Thng Tien Guan, Olivier Da Cruz, Sylvie Riu, Romain De Dormael, Kahina Abed, Olivia Touriguine, Roland Jourdain, Sylvie Cupferman, Luc Aguilar, Scott A. Rice

## Abstract

A balanced scalp microbiome is crucial for scalp health, yet the mechanisms governing this balance and the etiology of dysbiosis in scalp disorders remain elusive. We conducted a detailed investigation of the scalp and hair follicles, in healthy individuals and those with dandruff/seborrheic dermatitis (D/SD). It was demonstrated that the microbiome inhabiting hair follicles serves as a reservoir for the scalp microbiome, thereby integrating the scalp, follicle, and the hair into one functional unit. Using in vitro models, we further elucidated mechanisms governing the assembly and interactions of the follicular microbiome under healthy and D/SD conditions. We show that propionic acid, produced by *C. acnes*, plays a pivotal role in maintaining microbiome balance, with implications for scalp health, which was validated through a clinical study.

## Main Text

Dandruff and seborrheic dermatitis (D/SD) are disorders considered as the same basic condition differing only in magnitude (*1–3*). D/SD are characterized by abnormalities of the scalp, which includes flaking, inflammation, and barrier dysfunction that are largely driven by dysbiosis of the scalp microbiome (*1*, *4*, *5*). The microbiome dysbiosis in D/SD is mainly defined by increased absolute and relative abundance of the fungus, *Malassezia restricta* and the bacterium *Staphylococcus spp*., relative to the bacterium *Cutibacterium acnes* (*6–11*). Of these three organisms, *Malassezia* has been extensively studied due to early identification of its association with the D/SD conditions (*1*, *2*, *12*, *13*), making it the main target of most anti-dandruff treatments. These anti-fungal treatments are effective in relieving D/SD symptoms by keeping *Malassezia* numbers low, but they require regular application. In fact, stopping anti-fungal usage is associated with a strong relapse in the D/SD phenotype, with an increase in *Malassezia* numbers (*14*). This suggests that there is a potential reservoir of the scalp microbiome that replenishes the dysbiotic D/SD microbiome after treatment and may extend below the scalp surface. Interestingly, metagenomic sequencing of hair follicles from a small number of samples has indicated that the hair follicles are colonized by microbes (*15–17*). However, a detailed description of the hair follicle microbiome in a larger sample size, its potential connection with that of the scalp, and its role in the scalp conditions such as D/SD remain underexplored. Furthermore, the ecology and interaction dynamics between the key microbial species and the factors that affect such interactions under both healthy and D/SD conditions are not fully understood.

In this study, using hair follicles and scalp samples from 65 human volunteers (33 Healthy + 32 D/SD), we conducted a detailed investigation of the hair follicle microbiome composition and investigated if this microbiome is connected with that of the scalp, potentially acting as a reservoir. We further developed *in vitro* models to identify key metabolites that dictate the microbiome community structure in healthy and D/SD conditions. The significance of these effectors was revealed through a second clinical study, wherein we unambiguously demonstrate the role of a *C. acnes* postbiotic in reinforcing healthy scalp and follicle conditions. This study significantly advances our understanding of microbial ecology in skin and scalp, provides a mechanistic basis of onset of dandruff, and for the first time defines clear signatures for a healthy scalp and hair ecosystem.

### Microbiome signatures of the scalp and hair follicles unravel an interconnected scalp-follicle unit

To understand the mechanisms associated with the community structure of health and dandruff scalp microbiome, we conducted a clinical knowledge study in Singapore with two groups of subjects that were classified as, Healthy (n=33, total dandruff score = 0) and having Dandruff or Seborrheic Dermatitis (D/SD, n=32, total dandruff score >1.5 on a scale of 0 to 5). All volunteers were given a neutral shampoo to be used for three weeks (washout period), followed by biological sample collection at the end of the washout period. Representative samples for both scalp (one swab and one Squame/corneum disc) and follicles (2 sets of 10 hair follicles with intact outer root sheath) were collected for analyses.

A swab sample and a set of 10 hair follicles from each volunteer of Healthy and D/SD groups were subjected to microbiome analysis by shotgun metagenomic sequencing (Fig. 1A., S1A). The most abundant genus of the scalp microbiome of the Healthy group was *Cutibacterium* with a median relative abundance (MRA) of 0.84, and within this genus, *C. acnes* was the most abundant species (MRA = 0.77) (Fig. 1A). *Malassezia* and *Staphylococcus* genera were found in very low abundance, with a MRA of 0.07 and 0.03, respectively. *M. restricta* (MRA = 0.8) and *Staphylococcus capitis* (MRA = 0.75) constituted the most abundant species within their respective genera (Fig. 1A). In contrast, the most abundant genus in the D/SD group was *Malassezia*, with a MRA of 0.36, closely followed by *Cutibacterium* (MRA = 0.29), and *Staphylococcus* (MRA = 0.16) (Fig. 1A). At the species level, *M. restricta, C. acnes,* and *S. capitis* were the most abundant species within their respective genera, with MRA of 0.94, 0.91, and 0.76, respectively (Fig. 1A). These results strongly suggest that the relative abundances of the genera *Cutibacterium*, *Malassezia* and *Staphylococci* drive the signatures of healthy (higher relative abundance of *Cutibacterium spp.,*) or dandruff scalp (higher relative abundances of *Malassezia spp.,* and *Staphylococcus spp.,*) of the Singaporean population, which aligns with previous dandruff scalp microbiome studies (*6–8*).

**Fig. 1.**
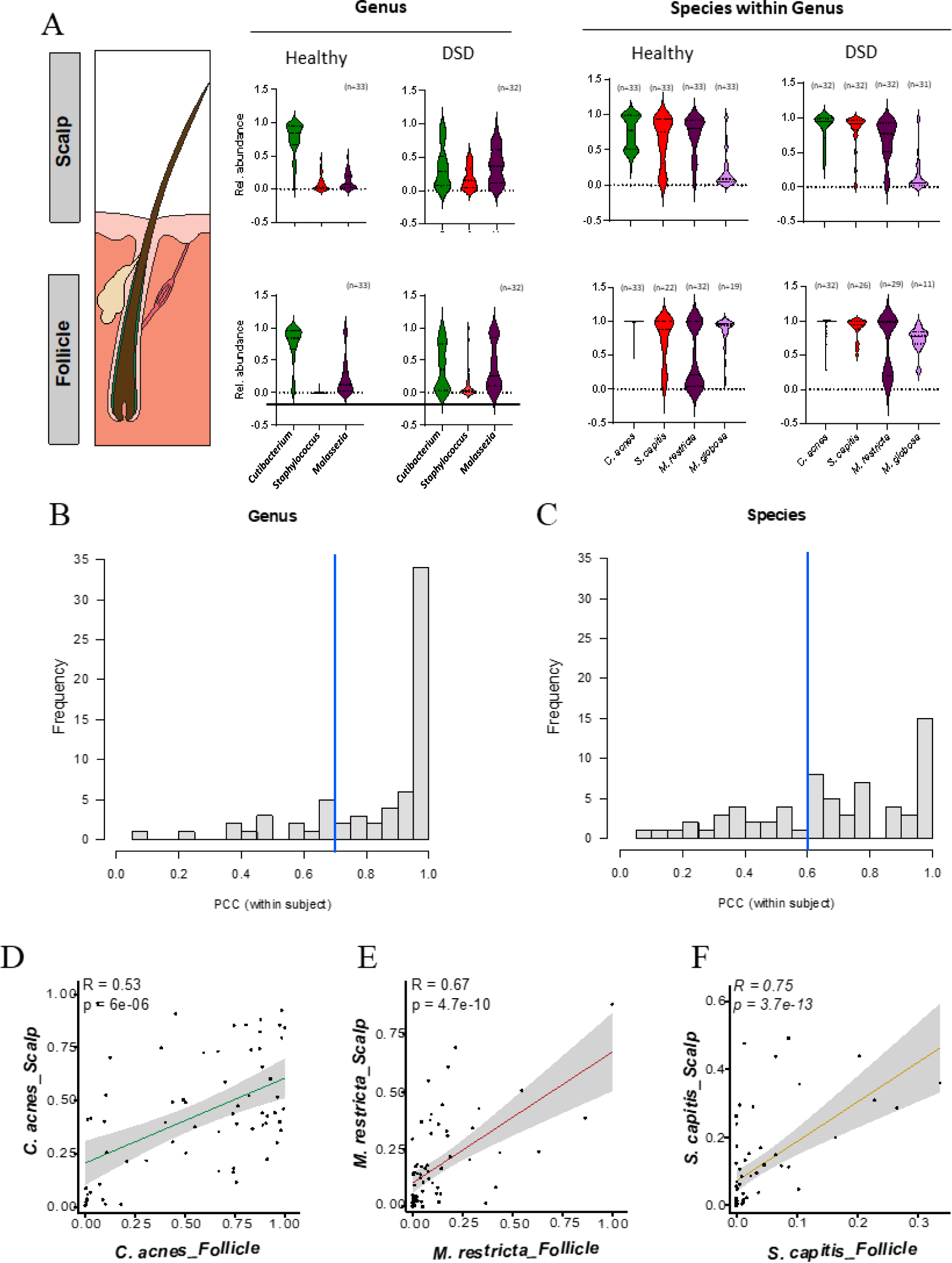
The scalp and follicular microbiome of healthy vs dandruff and seborrheic dermatitis (DSD) volunteers. Relative abundance of the key genera, Cutibacterium (C), Staphylococcus (S), and Malassezia (M), and their member species of the scalp and follicle microbiomes collected from healthy and DSD volunteers (A). Correlation analysis (Pearson’s correlation coefficient, PCC) of the relative abundance of the microbiome members between the scalp and follicle of each volunteer in both Healthy and DSD groups at genus (B) and Species (C) level. The blue line indicates the PCC cut-off for over 75% of the individuals compared. Correlation of relative abundance between scalp and follicle of C. acnes (D), M. restricta (E), and S. capitis (F).

Comparisons with other studies revealed that while *C. acnes* and *M. restricta* were the common species across all populations, differences were seen in the *Staphylococci*: *S. epidermidis* was predominant in the European (*6*) and American (*18*) populations. Together with *S. epidermidis, S. capitis* and *S. caprae* were found in higher abundance in Indian (*8*), Chinese (*7*), and South Korean population (*19*). In this study, *Staphylococcus* species were dominated by *S. capitis,* followed by *S. caprae* and *S. epidermidis*. It is not clear if this species level variation was due to different sequencing methods used, or if there is variation across different populations. Nevertheless, since the relative proportions of the *Staphylococcus* genus were similar across different populations, it is possible that different *Staphylococcus* species contribute to the same function within the skin microbiome. This functional redundancy, an ecological property where taxonomic units at the lower taxonomic ranks contribute to the same function, despite their taxonomic differences (e.g. species), has been suggested to be responsible for the resilience and stability of a community. For example, the gut microbiome of different individuals varies significantly at the species level, but the underlying composition of functional genes is similar (*20*). Therefore, it is quite likely that all of the *Staphylococcus spp*. (*S. capitis*, *S. caprae* and *S. epidermidis)* perform similar ecological functions on the scalp.

We next investigated the microbiome of the hair follicles from the same individuals to determine if the follicle microbiomes were like that of the scalp or if they represent a completely separate community. Microbiome composition for both healthy and D/SD follicles showed similar taxonomic composition as that of their scalp counterparts. Follicles from healthy volunteers had a higher proportion of *Cutibacterium* (MRA = 0.84), followed by *Malassezia* (MRA = 0.12) and *Staphylococcus* (MRA < 0.01). D/SD follicles exhibited a higher relative abundance of *Malassezia* (MRA = 0.36) and *Staphylococcus* (MRA = 0.04) with a lower relative abundance of *Cutibacterium* (MRA = 0.35) (Fig. 1A). At the species level, *C. acnes* and *S. capitis* constituted most of their respective genera both in Healthy (MRA = 0.99 and 0.88, respectively) and D/SD (MRA = 1 and 0.94, respectively) groups (Fig. 1A). For *Malassezia*, *M. globosa* and *M. restricta* had variable relative abundances, with *M. globosa* present only in about half of the samples in both groups. When present, *M. globosa* was found in higher abundance in both groups (MRA = 0.94 in Healthy and MRA = 0.77 in D/SD) (Fig. 1A). The exact role of *M. globosa*, including functional redundancy with *M. restricta*, in the follicles of this subpopulation requires further investigation. *M. restricta,* however, was found in all follicle samples, with very few exceptions in the D/SD group. It dominated the genus in the D/SD group with MRA of 0.94, strongly suggesting its role in the D/SD condition.

This data from follicles and swabs from the same volunteers also allowed us to perform correlation analysis (Pearson’s correlation coefficient, PCC) between the relative abundances of the microbiome members of follicles and the scalp to further test if these two niches possess a common core microbiome, and thus an interconnected microbiome unit. Over 75% of the individuals showed very high correlations (>0.8) between the scalp and follicle microbiome at the genus level, and moderate to high correlations (between 0.6 and 1) at the species level (Fig. 1B, C). Furthermore, to understand if the relative abundance of the key microbiome member species was similar among follicles and the scalp, correlation analysis was conducted for *C. acnes*, *S. capitis* and *M. restricta*. A positive correlation in the relative abundances of these major species was observed between the two regions (Fig. 1D, E, F), suggesting that scalp shares a common core microbiome with the follicles, connecting both the niches, despite the scalp being under constant exposure to environmental factors. Thus, microbiome data from the follicles and the scalp strongly suggest that both niches are interconnected and dysbiosis in D/SD conditions is not limited to the scalp surface alone but is also associated with the follicles. In fact, hair fibers from D/SD individuals compared to healthy control groups show significant differences in multiple parameters, ranging from diameter to shine and hardness (*21*), suggesting that the follicles, scalp, and hair could be functioning as one interconnected unit. These findings present a new paradigm wherein the scalp, follicles, and hair function as one interconnected unit rather than being independent niches and further defines new biomarkers for scalp health.

### Microbial biofilms in the hair follicles: a potential reservoir for the scalp microbiome

Metagenomic sequencing revealed that, despite differential environmental conditions on the scalp (aerobic, exposed to the atmospheric environment) and follicles (hypoxic, protected from direct exposure to atmospheric environment), there is a core microbiome that connects both these niches irrespective of a healthy or D/SD scalp condition. This is particularly relevant for *C. acnes*, an aerotolerant anaerobe, whose colonization would be impaired under oxygen rich conditions, unless present under a protective environment, such as a biofilm. Interestingly, the presence of biofilms on hair follicles has been suggested previously with a small number of samples (*22*, *23*), but to date, lacks unambiguous demonstration. To ascertain if microbial biofilms are present on scalp and follicles, we analyzed tape strips from the scalp surface and hair follicles collected from both healthy and D/SD volunteers by scanning electron microscopy (SEM). Unlike indirect methods such as metagenomic sequencing or cultivation, SEM imaging allows for visualization of microbes and biofilm structures that would provide direct evidence for their presence.

Analysis of tape strips revealed a randomly collected biological material, mostly resembling parts of the stratum corneum, that varied between tape strips and did not conform with metagenome sequencing results (Fig. S2). It is possible that the tape strips only collected a few microbes or loose biofilms from the top layer of the scalp and are not representative of the scalp surface. Nevertheless, microbial structures on these biological materials could be visualized, with biofilm like structures present on a few samples (Fig. S2). Based on the dominant taxa identified through metagenomic analysis (section 2.1), the observed microbes presumably represent *Malassezia spp*. (budding/globular structure, ∼5 µm), *Staphylococcus spp.* (spherical/cocci structure, ∼1 µm), and *Cutibacterium spp.* (rod shaped, ∼1.5 µm). Due to this limitation of the sampling procedure, we concluded that while microbes and biofilms were found on the scalp surface samples, a widespread presence of biofilm on the scalp surface requires further investigation, perhaps using scalp biopsies. We next visualized the hair follicles, which were carefully extracted from scalp to maintain the structure of outer root sheath (ORS). Each hair follicle was classified into three regions to represent the infundibulum, isthmus, and bulb; each of these regions were imaged at 250X, 1500X, and 5000X magnification to ensure that all details were captured (Fig. 2). At least three hair follicles from each volunteer were analyzed to be confident with the observations. Irrespective of their healthy or D/SD origin, all hair follicles were colonized by microbes that were embedded in a matrix like structure associated with the ORS, suggesting the presence of a biofilm (Fig. 2A, B).

**Fig. 2.**
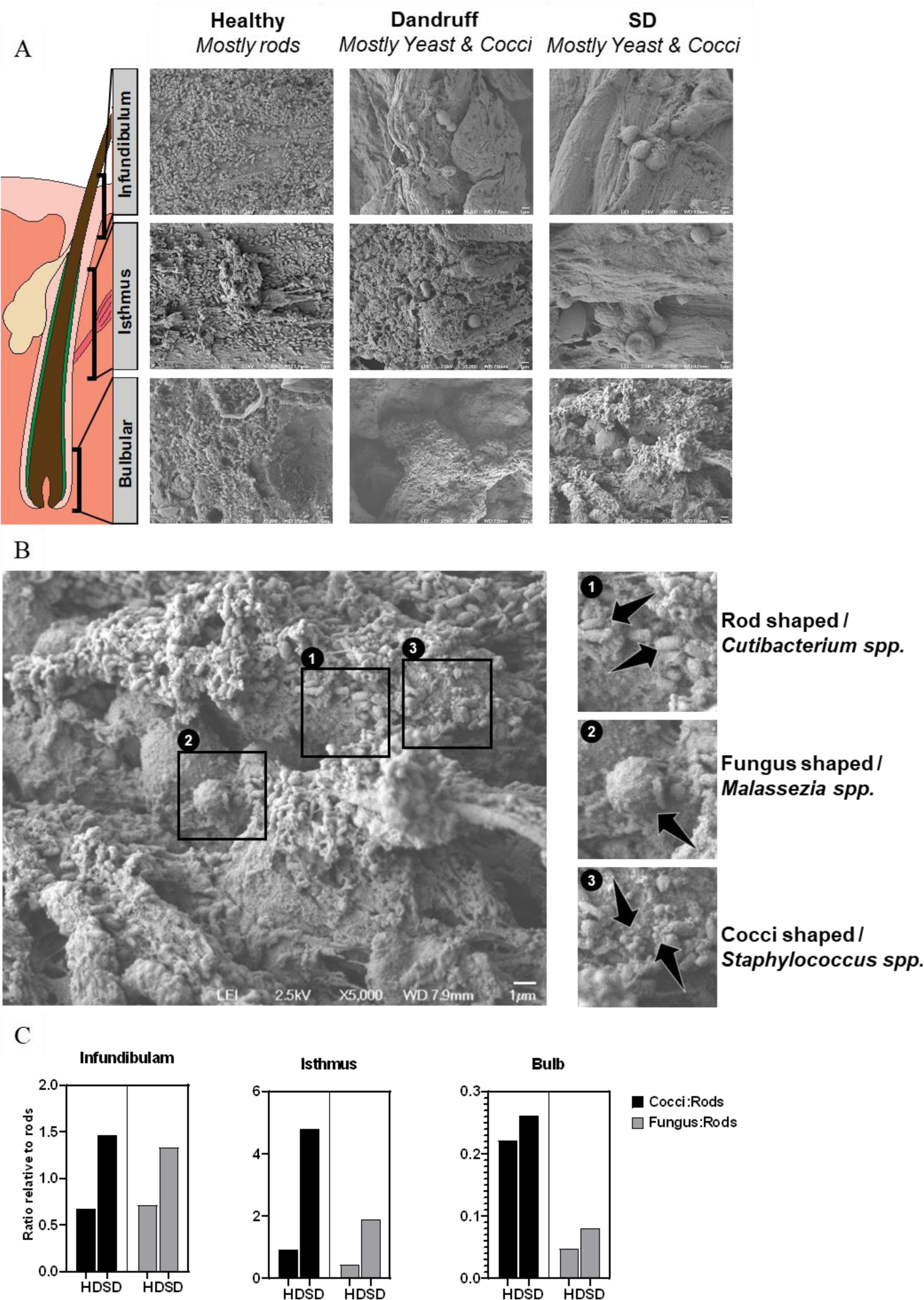
Scanning electron microscopy analysis of hair follicles. Images of representative hair follicles of healthy, dandruff (DFF) and seborrheic dermatitis (SD) volunteers (A). Microbial colonization was visualized at 5000 X magnification at the infundibulum, isthmus and the bulb regions of the hair follicle. Images from all the samples are presented in the supplementary material. Microbes were putatively identified based on their shape and size (B). The ratio of Cocci (Staphylococcus spp.) and Fungus (Malassezia spp.) shaped microbes compared to Rods (Cutibacterium spp.) in the three regions of hair follicle derived from semi-quantitative estimation of different microbial population.

When comparing the follicle biofilms from healthy and D/SD individuals, a relatively higher density of cells was always found on the infundibulum and the bulb regions. Sometimes, an extension of the biofilm structure was seen on the hair shaft, which was heavily colonized with microbes (Fig. S3). Based on metagenomic identities (section 2.1) as well as the shape and size of the microbial structures, we inferred that most of the healthy follicles were heavily colonized with *Cutibacterium spp.*, while both *Staphylococcus spp.* and *Malassezia spp.* were present at a very low abundance. In contrast, images from the D/SD follicles showed a higher abundance of *Malassezia spp.* and *Staphylococcus spp.* in addition to the *Cutibacterium spp.* (Fig. 2A). We further established a five-point (0 to 4) scoring system to assess the relative proportions of these microbial structures found on the follicles (supplementary methods). This semi-quantitative analysis of over 1700 SEM images (65 volunteers x 3 replicate samples x 3 follicle regions x 3 replicate images per region, Supplementary data 1) showed that the ratio of *Staphylococcus spp.* (cocci) and *Malassezia spp.* (fungus) to *Cutibacterium spp.* (rods) was low in healthy, and high in D/SD follicles (Fig. 2C). These results from the image analysis of biofilms were aligned with metagenomic sequencing results from the hair follicle samples (section 2.1). Thus, through a systematic analysis of SEM images of hair follicles from Healthy and D/SD groups, we conclude that hair follicles are colonized by microbes embedded in biofilm structures, and that a healthy follicle microbiome has a distinct microbial signature, dominated by rod like organisms that are likely to be *C. acnes*. Taken together, it is likely that these follicular biofilms act as a reservoir for the scalp microbiome, and that changes in the follicular niche would impact the scalp surface microbiome.

Under healthy conditions, the follicular niche is hypoxic (*24*, *25*), which is required for normal functioning of cellular pathways such as those regulated by Hypoxia-inducible factor-1α (*26*), which also leads to more effective maintenance of an intact epidermal barrier (*27–29*). *Cutibacterium* species grow under such hypoxic conditions and have been shown *in vitro* to activate longevity associated genes like the silent mating type information regulation 2 homolog-1 (SIRT1) and telomerase reverse transcriptase (TERT) genes, which further contribute to repair processes in the scalp and hair growth (*30*). One of the hallmarks of a dandruff is the increase in the trans epidermal water loss (TEWL), which indicates that the scalp integrity is compromised and would lead to a change in oxygen saturation, potentially compromising the hypoxic nature of the follicular regions (*3*, *5*, *13*, *31*, *32*). Moreover, acute skin barrier defects are associated with stimulation of vascular endothelial growth factor-A that promotes capillary growth thereby increasing the local oxygen saturation (*33*). These changes in oxygen saturation levels would lead to an unfavorable environment, including increased oxidative stress, which would limit *C. acnes* growth and favor facultative anaerobes such as *Malassezia* and *Staphylococci*. An increase in the abundance of *M. restricta* would further lead to an increase in lipid oxidation, exacerbating the barrier dysfunction (*13*, *32*), and contributing to a vicious cycle where the D/SD state is maintained. To test this hypothesis and to understand the mechanisms that drive the healthy microbiome composition, we recapitulated the follicle microbiome *in vitro* and studied the model under healthy (hypoxic) and D/SD (aerobic) conditions, wherein the community would experience higher oxidative stress due to exposure to atmospheric oxygen.

### Recreating the follicular environment in vitro replicates healthy skin microbiome community

To create an *in vitro* biofilm model representative of the follicles, a growth medium was developed to co-culture the key members of the follicle microbiome (*C. acnes*, *M. restricta*, and *S. epidermidis* - a common *Staphylococcus* species across all populations). Species-specific fluorescent *in situ* hybridization (FISH) probes, and culture-based methods were also developed to track and enumerate these microbes either in mono- or co-culture conditions. These tools allowed investigation of the interactions between these three species and facilitated a deeper understanding of community dynamics under different growth conditions.

To emulate the biofilm community associated with healthy follicles, experiments were performed under anaerobic conditions with identical cell densities for each species (1 × 10^5^ cells/ml). Under anaerobic monoculture conditions, *C. acnes* and *S. epidermidis* readily formed biofilms, while *M. restricta* did not (Fig. 3A). Moreover, no biofilm formation was seen for *M. restricta* when grown in mDixon medium that was optimized for robust growth of *Malassezia*, suggesting that *M. restricta* does not form a biofilm. Amongst all two species co-culture experiments, only a combination of *C. acnes* and *S. epidermidis* (C+S) led to a significantly higher biofilm biomass as compared to their monocultures (Fig. 3A). Addition of *M. restricta* to *C. acnes* (C+M) or *S. epidermidis* (S+M) or to C+S cocultures (C+S+M), did not lead to any significant increase in biofilm biomass (Fig. 3A). FISH analysis of the mature biofilm (Day 4) further revealed that *C. acnes* grew in higher abundance in the three-species biofilm, with a lower abundance of *S. epidermidis* (Fig. 3B). The majority of *M. restricta* cells emitted autofluorescence in the FITC channel with no labelling from the FISH probes, suggesting that *M. restricta* viability could be compromised (Fig. 3B). Visualization using SEM imaging confirmed that most of the cells had collapsed cell wall structures (Fig. 3C), indicating that viability of *M. restricta* under healthy conditions is indeed compromised. Therefore, FISH and SEM image analysis suggested that under healthy follicle conditions, *C. acnes* dominates the community with a lower abundance of *S. epidermidis* and *M. restricta*, as was found in the clinical samples through metagenomic and SEM analyses (sections 2.1 and 2.2).

**Fig. 3.**
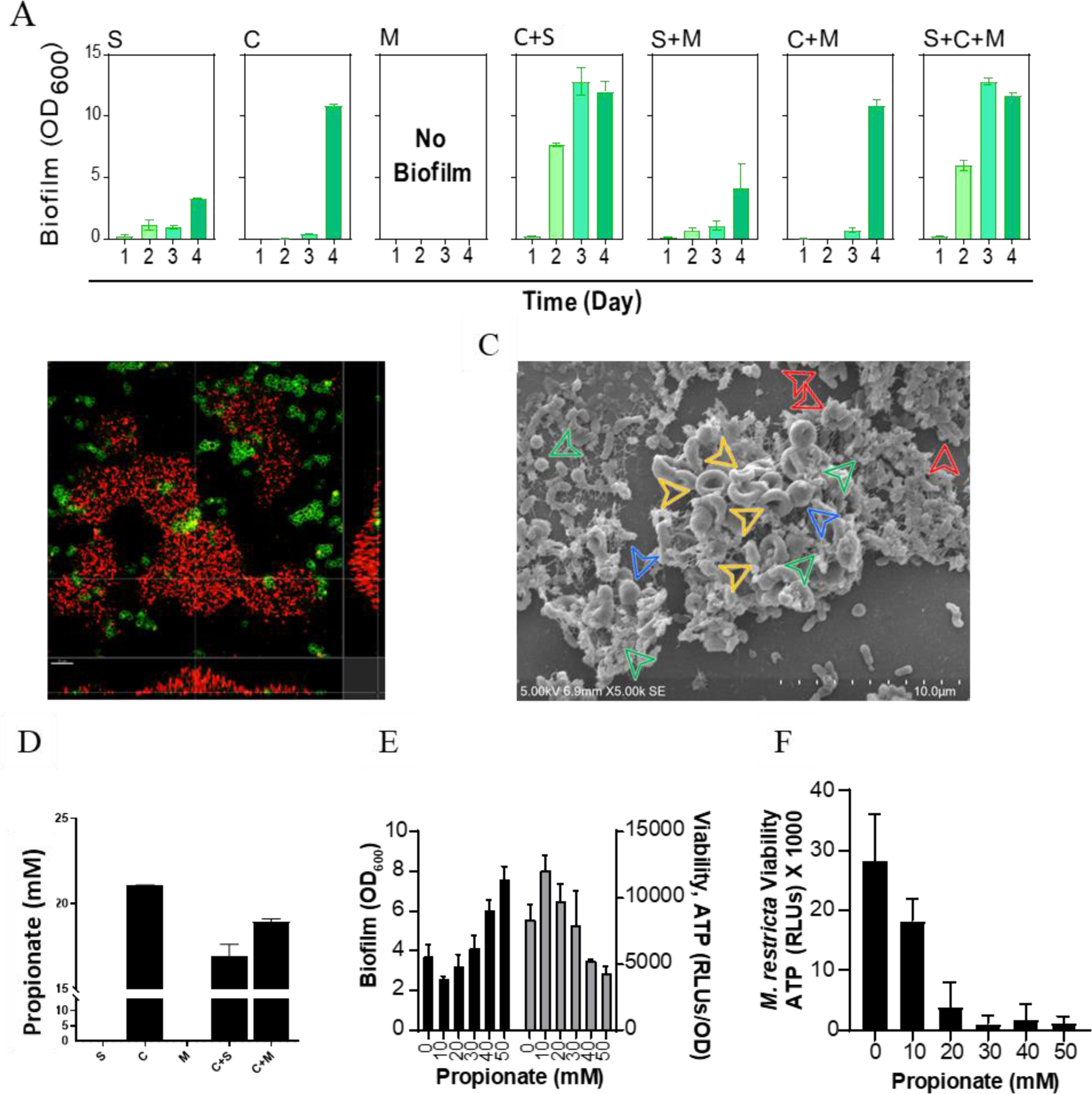
Biofilm formation and interaction of the skin microbes under anaerobic conditions. Quantification of biofilms formed by individual microbes, S. epidermidis (S), C. acnes (C) and M. restricta (M) as well as their combinations in co-culture using the crystal violet biofilm assay (A). Confocal laser scanning microscopy of C. (Red), S (Green) and M (Blue) in single- and multi-species biofilm visualized using Fluorescent in-situ hybridization (B). SEM imaging of the co-culture biofilm containing (C). Arrows indicate C (Red), S (Green), intact M (Blue) and cell wall collapsed M (Yellow). Estimation of propionate produced by the skin bacteria biofilms when grown as monoculture and co-culture, and with M. restricta (D). Biofilm formation and viability of S. epidermidis with propionate (E). Viability of M. restricta after 24 h of Propionate treatment (F).

Despite starting biofilm experiments with the same number of each member species (1 × 10^5^ cells/ml), the relative abundances of each species in mature *in vitro* biofilms mimicked the microbiome signatures of healthy follicles, with *C. acnes* dominating the community and possibly controlling the growth of *S. epidermidis* and *M. restricta*. This *in vitro* follicle microbial community model was further used to dissect the mechanisms by which *C. acnes* controls the growth of both *S. epidermidis* and *M. restricta* and thereby maintains a healthy scalp microbiome community.

### C. acnes postbiotic dictates community dynamics in healthy follicle

Interactions between members of the microbial community are often driven by secreted metabolites that can alter population dynamics through cross feeding and/or growth inhibition mechanisms (*34–37*). More specifically, microbiome derived short chain fatty acids (SCFAs) play critical role in gut health and metabolism (*38–42*), including regulation of pathogenesis in the gut (*43–46*). Interestingly, gut derived SCFAs were also shown to play a role in inflammatory skin conditions, including modulation of skin immune system and skin barrier properties (*47–50*). External addition of a SCFA, butyrate, to organ cultured hair follicles was recently shown to delay the catagen progression accompanied by stimulation of mitochondrial activity and antibiotic peptide expression, suggesting that SCFAs can have a direct impact on the hair follicle function (*17*). Similarly, addition of SCFAs to *S. epidermidis in vitro* was shown to inhibit biofilm formation of the skin commensal bacteria (*51*). However, the role of SCFAs that are locally produced in the follicular environment with respect to the resident microbiome community is largely unknown.

To check if a secreted SCFA could drive the antagonistic relationship of *C. acnes* (C) with both *S. epidermidis* (S) and *M. restricta* (M), we analyzed the secretome from two conditions: 1) C, S and C+S, 2) C, M and C+M. *C. acnes* produced both acetic acid and propionic acid, while *S. epidermidis* produced only acetic acid among the SCFAs tested (acetic acid, propionic acid, butyric acid, and Valeric acid). Analysis of supernatants from co-culture combinations revealed significant changes in the production of propionic acid. Propionic acid concentration, when measured under co-culture conditions (Fig. 3D, C+S and C+M) was lower when compared to the monocultures on Day 3, suggesting that either *C. acnes* produces less propionic acid under these conditions, or the produced propionic acid may be consumed by the co-occurring species. We therefore investigated if propionic acid could be a central effector through which *C. acnes* exerts control on both *Staphylococcus* and *Malassezia* species in the follicle microbiome.

To dissect the potential role of propionic acid in the healthy follicle microbiome, we analyzed the impact of increasing amounts of propionic acid on *S. epidermidis* and *M. restricta,* under anaerobic conditions. Supplementation of propionic acid (sodium salt) to *S. epidermidis* biofilms led to increase in biofilm biomass, however, this was also associated with a reduction in viability (Fig. 3E). This suggests that *C. acnes* uses propionic acid as an effector to modulate the biofilm formation and viability of *S. epidermidis* in healthy follicle microbiomes. Interestingly, while the physiological concentration of propionic acid on the scalp or in the follicle is unknown, complete killing was not observed at all concentrations tested (Fig. 3E). This suggests that *C. acnes* could potentially use propionic acid to limit the growth of *S. epidermidis* but is not likely to be the sole effector. Competitive interactions between *C. acnes* and *S. epidermidis* have been reported previously, and a role of *C. acnes* secreted thiopeptides, e.g. Cutimycin, has been suggested to play an important role in suppressing growth of *S. epidermidis* (*52*). It is possible that *C. acnes* uses a range of effectors including propionic acid and Cutimycin to exert different levels of control on *S. epidermidis* within the follicle. Nevertheless, these experiments validate a role of propionic acid in controlling the growth and biofilm formation of *S. epidermidis* within the follicles. The addition of propionic acid to *M. restricta* cultures under anaerobic conditions revealed a dose dependent killing effect, with complete growth inhibition at 30 mM (Fig. 3F). This suggests that *C. acnes* produces propionic acid to completely control the growth of *M. restricta* in the healthy follicles. These data match clinical findings within the follicles, wherein *C. acnes* was found at a high relative abundance (section 2.1) and would generate a high, localized concentration of propionic acid, which would control the follicle microbiome community through direct action on both *S. epidermidis* and *M. restricta*.

### D/SD conditions of the follicle skew community balance favoring malassezia growth

We next investigated the community dynamics and the role of propionic acid under D/SD conditions where the scalp barrier is defective and is associated with higher oxidative stress levels. Precise oxygen concentrations or the oxidative stress levels within the follicles in D/SD conditions are unknown and are difficult to replicate experimentally. Conventional approaches to introduce oxidative stress such as addition of hydrogen peroxide while being very effective, represent an extreme situation that might not be physiologically relevant. Therefore, experiments were performed under atmospheric oxygen saturation conditions (aerobic) that are known to introduce ambient levels of oxidative stress under *in vitro* conditions (*53*, *54*). Under aerobic mono-culture conditions, *S. epidermidis* formed a stable biofilm that matured around 48 h, and gradually decreased at later timepoints (Fig. 4A). *C. acnes*, being an aerotolerant anaerobe, did not grow or form biofilms. *M. restricta* showed aerobic growth with no biofilm formation observed at any time point. Amongst the 2- and 3 -species co-culture experiments, biofilm formation was similar to that of *S. epidermidis* monocultures up to 48 h, except for C+M, where no biofilm formation was observed. Interestingly, at later timepoints (day 3 and 4), no reduction in the biofilm biomass was seen for all co-culture conditions (Fig.4A), and this effect was particularly prominent for all combinations containing *C. acnes*. To understand the community composition in the biofilm co-cultures, we performed FISH analysis (Fig. 4B). Biofilm co-cultures of the three species revealed a mutualistic growth pattern, wherein all three microbes, *S. epidermidis*, *M. restricta* as well as *C. acnes* showed active grow (Fig. 4B). *M. restricta*, and *C. acnes* grew as microcolonies within the *S. epidermidis* biofilm matrix and were rarely associated with each other (Fig. 4B). Furthermore, SEM analysis revealed that most of the *M. restricta* cells under aerobic conditions were intact (Fig. 4C). The growth of *C. acnes* was unexpected as the aerotolerant anaerobe does not grow under aerobic conditions, suggesting a mutualistic interaction between *C. acnes* and *S. epidermidis*.

**Fig. 4.**
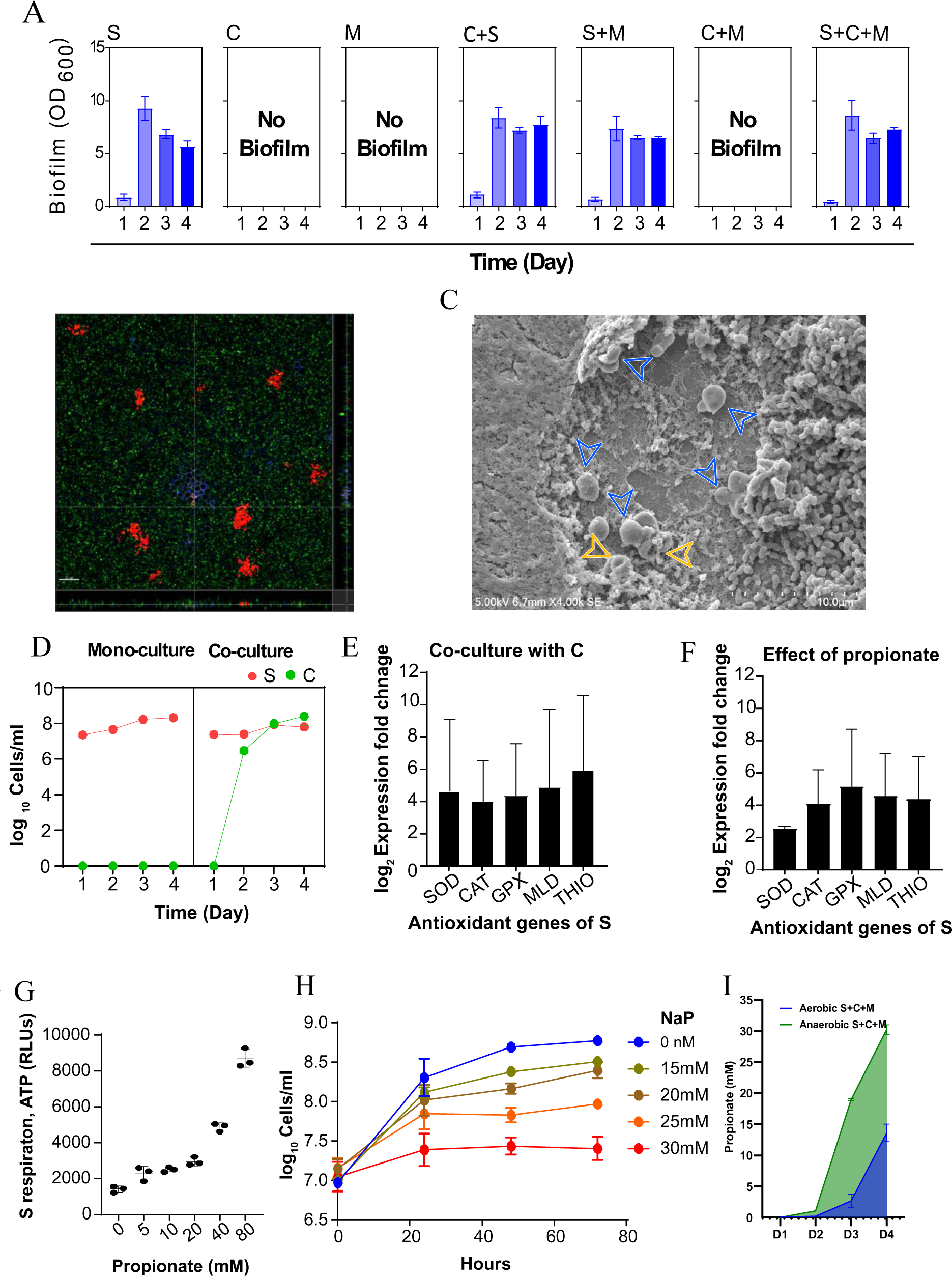
Biofilm formation and interaction of the skin microbes under aerobic conditions. Quantification of biofilms formed by individual microbes, *S. epidermidis* (S), *C. acnes* (C) and *M. restricta* (M) as well as their combinations in co-culture using crystal violet assay (A). Confocal laser scanning microscopy of *C* (Red), *S* (Green) and *M* (Blue) in single- and multi-species biofilm visualized using Fluorescent in-situ hybridization (B). SEM imaging of the co-culture biofilm containing (C). Arrows indicate intact M (Blue) and cell wall collapsed M (Yellow). Growth of C and S as single- and multi-species under aerobic conditions (D). Change in gene expression of antioxidants, super oxide dismutase; SOD, catalase; CAT, glutathione peroxide; GPX, malate dehydrogenase; MLD, and thioredoxin; THIO, of S during biofilm formation together with C under aerobic conditions (E). Change in antioxidant gene expression of S when treated with 5 mM sodium propionate under aerobic conditions (F). Estimation of ATP from S forming biofilms when treated with increasing concentration of propionate (G). Growth of *M. restricta* with Propionate (H).

We further quantified *C. acnes* and *S. epidermidis* CFUs within the C+S co-cultures, which revealed that *C. acnes* indeed grows exponentially in the presence of *S. epidermidis* (Fig. 4D). Robust growth of *C. acnes* microcolonies within *S. epidermidis* biofilms under aerobic conditions could be mediated through the creation of anoxic zones within *S. epidermidis* biofilms. Creation of such zones would only be possible through rapid utilization of oxygen, mediated by increased metabolism and respiration of *S. epidermidis* within the biofilm matrix. One way to test increased respiration of *S. epidermidis* within the co-cultures is to selectively quantify the expression of *S. epidermidis* antioxidant genes that would help in neutralization of reactive oxygen species (ROS) produced because of increased respiration of oxygen. We indeed observed increased expression of Super oxide dismutase (SOD), Catalase (CAT), Glutathione peroxidase (GPX), Malate dehydrogenase (MLD) and Thioredoxin (THIO), in *S. epidermidis* when co-cultured with *C. acnes* compared to monocultures of *S. epidermidis* (Fig. 4E). While a role of other naturally produced antioxidants cannot be ruled out (for example the secretion of bacterial antioxidant protein, RoxP, released by *C. acnes* (*55*)), this data clearly indicates that *S. epidermidis* supports *C. acnes* growth through rapid consumption of oxygen and creation of an anoxic environment, through a process that is facilitated by anti-oxidative mechanisms.

Since propionic acid is a metabolite that directly feeds into the central carbon metabolism pathways, we hypothesized that it could be the effector that *C. acnes* uses to accelerate the respiration rate of *S. epidermidis*, and in turn facilitate its own growth. To test this possibility, we quantified antioxidant gene expression from *S. epidermidis* monocultures supplemented with propionic acid and found similar increases in antioxidant gene expression (Fig. 4F). To further demonstrate the role of propionic acid in increasing *S. epidermidis* respiration, we quantified ATP production in *S. epidermidis* biofilms supplemented with propionic acid and found a dose dependent increase in ATP production with propionic acid supplementation (Fig. 4G). These results show that to grow under oxygen rich conditions, *C. acnes* uses propionic acid to generate anoxic zones created as a result of accelerated respiration of *S. epidermidis*, a process facilitated by anti-oxidative mechanisms.

To determine the effect of propionic acid on *M. restricta* under aerobic conditions, we supplemented propionic acid to *M. restricta* monocultures. A dose dependent growth inhibition was observed with complete inhibition at 30 mM (Fig. 4H). However, under coculture conditions, *M. restricta* growth inhibition was not seen (Fig. 4B), suggesting that propionic acid concentration may be lower under the aerobic conditions. When measured, the propionic acid concentration in the three species biofilm under aerobic conditions was significantly lower than that of anaerobic conditions (Fig. 4I), suggesting that the available propionic acid is sufficient to be utilized by *S. epidermidis* to create anoxic zones for facilitating *C. acnes* growth, but below concentrations necessary to inhibit *M. restricta*.

Taken together, this data suggests that *S. epidermidis* plays a key role in mediating the survival of *C. acnes* as well as *M. restricta* under aerobic conditions. These mutualistic interactions potentially explain the increase in the relative abundance of both *S. epidermidis* and *M. restricta* to *C. acnes* in the D/SD follicles (Fig. 1A, 2).

### Propionic acid induces an iron-starvation response that inhibits the growth of *M. restricta*

We next determined the mechanism of propionic acid induced *Malassezia* growth inhibition through metabolomic and transcriptomic analysis of *Malassezia* cells treated with propionic acid. Metabolomic analysis revealed that treatment of *M. restricta* with propionic acid resulted in a lower abundance of glycolytic intermediates (Fig. 5A, Glycolysis), accumulation of TCA cycle intermediates (Fig. 5B, TCA cycle) and an increased concentration of free amino acids (Fig. 5C, amino acids). The NAD+ relative to NADH, NADP+ relative to NADPH, and ADP relative to ATP ratios all higher in treated cells (Fig. 5D, energy carriers), suggesting significant reduction in respiratory activity that could eventually lead to cell death. Interestingly, similar metabolic changes have been reported for yeast cells undergoing iron starvation (*56*), suggesting that iron starvation could be the mechanism of propionic acid mediated growth inhibition of *M. restricta*. To test this, we quantified the intracellular iron concentration and enzymatic activity of an iron-sulfur protein, aconitase, in *M. restricta* cells treated with propionic acid. As expected, treated cells had significantly lower intracellular concentrations of iron (Fig. 5E), which was associated with a decrease in aconitase activity (Fig. 5F).

**Fig. 5.**
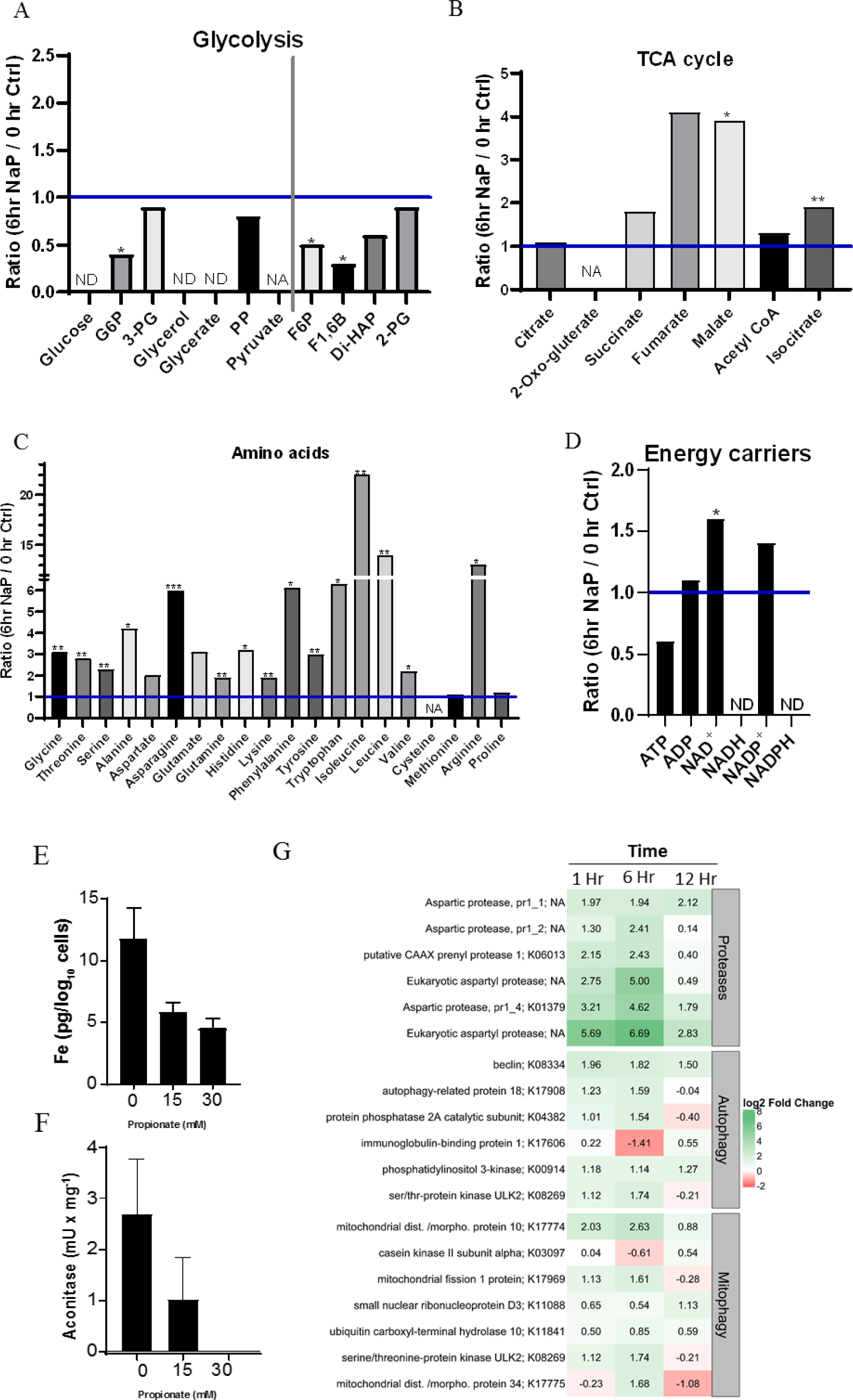
Mechanism of propionate mediated growth inhibition of *M. restricta*. Relative quantification of central carbon metabolic intermediates of glycolysis (A), TCA (B), amino acids (C) and energy carriers (D) of *M. restricta* treated with 30 mM sodium propionate for 6 h compared with 0 h. Blue line represents concentration of metabolites before propionate treatment (0 h). Intracellular metal iron concentration of *M. restricta* (E). Activity of an iron-sulfur enzyme, aconitase (F). Genes coding for proteases, autophagy and mitophagy enriched at 30 min, 3 h and 6 h after treatment of 30 mM propionate (G).

Transcriptomic analysis revealed that propionic acid treatment led to changes in transcripts involved in central carbon metabolism and an increased abundance of transcripts associated with proteases, autophagy and mitophagy (Fig. 5G). Induction of autophagy and mitophagy in response to iron starvation has been shown in several fungal species (*57*, *58*), further confirming iron starvation as the probable mechanism of propionic acid induced *M. restricta* inhibition. Interestingly, inhibition of *Malassezia spp.* by Zinc Pyrithione (ZPT), a common anti-dandruff and anti-malassezia agent, also targets the Iron-Sulphur proteins (*59*).

### Propionic acid improves dandruff and reduces Malassezia colonization on scalp

To demonstrate the role of propionic acid in boosting a healthy scalp microbiome, we conducted a proof-of-concept clinical study on dandruff volunteers to evaluate if propionic acid can have an anti-dandruff efficacy *in vivo*. Since the activity of propionic acid is dependent on antioxidant mechanisms under D/SD conditions (section 2.5), an antioxidant, Ascorbyl glucoside (AG), was supplemented with propionic acid. A hydroalcoholic lotion as a vehicle for the active ingredients containing 1% sodium propionate and 3% AG was formulated and a randomized, comparative, monocentric, double-blind, interindividual clinical trial, versus vehicle alone was conducted at a single investigational site with 57 volunteers having moderate to severe dandruff (total dandruff score ≥ 4.5). The test lotion was applied daily by volunteers, at home for four weeks, with a neutral shampoo 2 to 3 times per week. Efficacy was assessed by clinical scoring and microbiome evaluation (*Malassezia*, *Cutibacterium* and *Staphylococcus* species) at day 0, and at the end of treatment. The propionic acid formulation had a two fold reduction on the dandruff scoring compared to the vehicle (Fig. 6A), alongside a significant reduction in the *Malassezia* to *Cutibacterium* ratio (Fig. 6B). No change in *Staphylococci* to *Cutibacterium* ratios was seen (Fig. 6C). Thus, the clinical data supported the anti-*Malassezia* effect of propionic acid and suggests that propionic acid can be used as a postbiotic to maintain a healthy scalp microbiome and resolve scalp disorders like dandruff. These findings establish that propionic acid is an important marker for scalp health, contributes to a more resilient scalp and can also help resolve scalp disorders. These findings are well aligned with the extensive literature on gut microbiomes, where the beneficial role of SCFAs is well characterized (*40*, *42*, *60*).

**Fig. 6.**
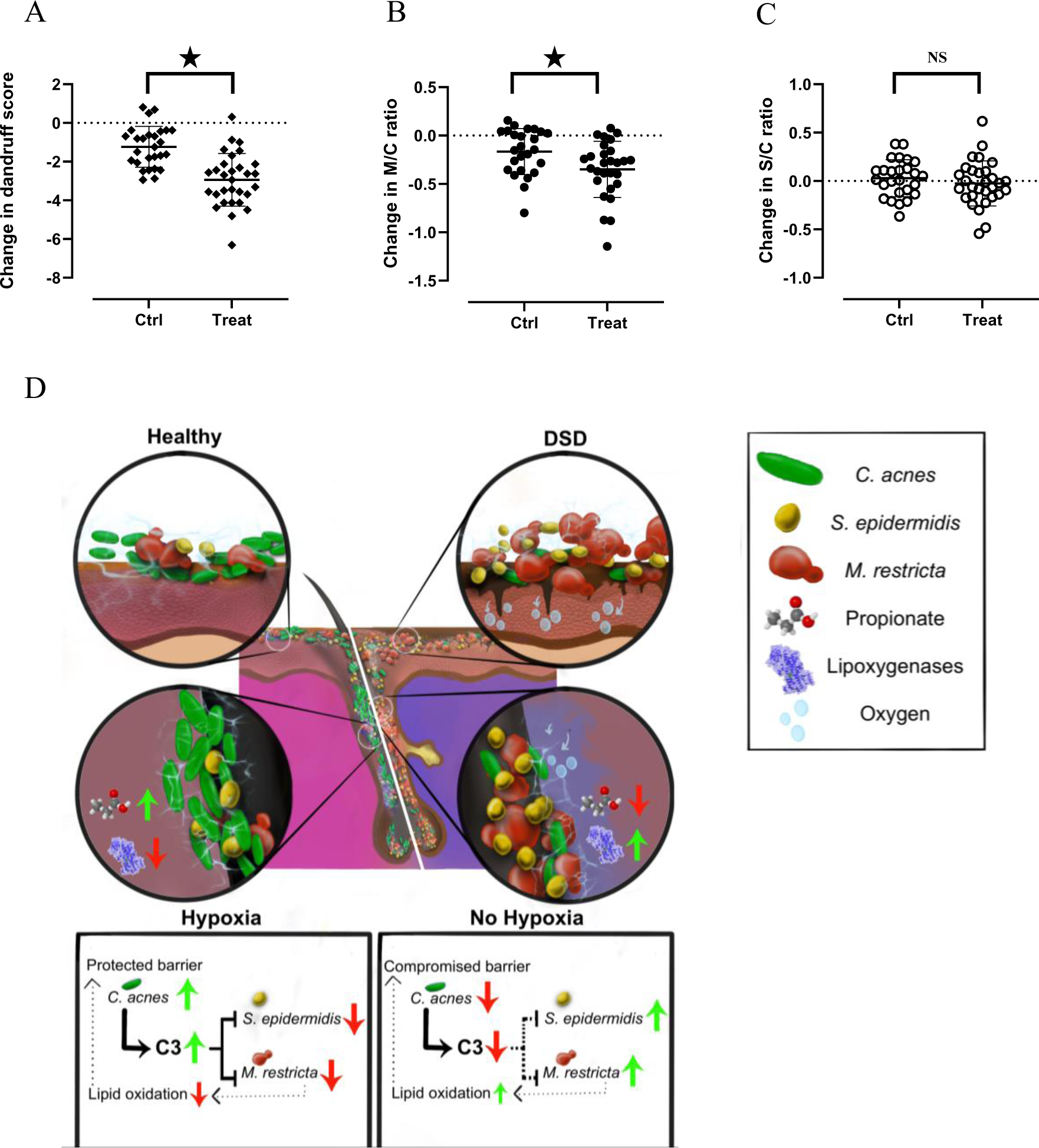
Role of Propionate in the microbial ecology of the healthy and DSD scalp. Change in total dandruff score (A), and M. restricta numbers in relation to C. acnes (B) after 28 days of application of formulation with and without propionate. Schematic representing the mechanisms by which the members of healthy and DSD follicle-scalp microbiome interact and impact the host health (C). * P< 0.05.

### A new paradigm in scalp microbiome research

The microbiome has been the focus of understanding host health for more than a decade and many studies on skin/scalp disorders have described the association of microbiome changes mainly at the taxonomy level. Here, we took an integrated clinical and synthetic ecology approach to functionally dissect the role of microbiome in dandruff and scalp health. This led to the identification of effectors that drive these phenotypes, which were further validated in a clinical efficacy study.

Findings from the clinical knowledge study suggest that the scalp and follicles share a common microbiome and function as one interconnected follicular unit. This strongly advocates for holistic studies that investigate the hair, follicle and the scalp as an interconnected niche. Presence of microbial biofilms within the hair follicles as a reservoir, further explain the inherent resilience of the scalp microbiome to different environmental exposures. This knowledge of the involvement of the hair follicles in scalp biology opens many new avenues to manage scalp disorders in a holistic way.

*In vitro* studies helped in identification of mechanisms that drive the healthy or dandruff community within the follicles and scalp. In healthy follicles under low oxygen saturation, *C. acnes* thrives and produces high amounts of propionic acid that exert a direct control over the other two species and helps maintain the dominance of *C. acnes* within the community. In D/SD conditions, a defective scalp barrier, and the resulting oxidative stress, compromises the growth of *C. acnes*, which in turn switches from antagonistic to a mutualistic interaction with *S. epidermidis*, thereby facilitating its own growth and survival over maintaining dominance in the community. This renders *M. restricta* unchecked, which increases in numbers and produces effectors like lipoxygenases that further worsen the epithelial barrier. Thus, we reveal that a disruption of the host barrier drives dysbiosis within the follicular unit, which leads to secretion of microbial effectors contributing directly to host barrier dysfunction, leading to a vicious cycle that defines D/SD etiology. Identification of propionic acid as a key scalp health biomarker further provides clues to building microbiome-based therapeutics for scalp health.

We conducted a clinical efficacy study to directly demonstrate the therapeutic role of propionic acid in dandruff condition. Application of propionic acid helped reset the community balance and led to alleviation of dandruff symptoms, alongside reduction in *Malassezia* (Fig. 6B). This highlights the importance of short chain fatty acids in skin/scalp biology, analogous to what is observed in the gut (*36–38*). In fact, recent reports suggest a strong role of the skin microbiome in skin aging and longevity (*38–40*). This study provides new insights into scalp biology and paves the way for the next generation of microbiome targeted therapeutics.

## Acknowledgments

The authors would like to acknowledge the support by the National Research Foundation and Ministry of Education Singapore under its Research Centre of Excellence Program to the Singapore Centre for Environmental Life Sciences Engineering, and Ministry of Education AcRF-Tier 1 grant. The authors would also like to thank Audrey Guéiniche, Cécile Clavaud, Quah Yiling Samantha, Chew Yean Ming, Li Yongling Adelicia and Kai Wei Kelvin Lee for their inputs during the initial stages of the project. The authors also thank Prof. Alain Filloux for his valuable comments that helped shape the final version of the manuscript.

## Funding

Singapore Centre for Environmental Life Sciences Engineering EDUN C33-62-036-V4 (VRR, SC, HFBJM, KPR, KS, CYY, EK, RW, LX, SAR)

Ministry of Education AcRF-Tier 1 grant 2017-T1-002-155 (VRR, SAR)

L’oral Researh and Innovation

## Author contributions

Conceptualization: VRR, TC, SAR

Methodology: VRR, TC, RW, VK, NB, AJ, STTG, SAR,

Investigation: VRR, KW, SC, AS, HFBJM, KPR, KS, CYY, EK

Visualization: VRR, TC, KPR, RW, LX

Funding acquisition: SAR

Project administration: VRR, SAR, TC

Supervision: VRR, SAR, TC

Writing – original draft: VRR, TC

Writing – review & editing: VRR, TC, SAR

## Data and materials availability

Sequence data are available at the National Centre for Biotechnology information. Image data are available at the NTU-DR data repository.

## Supplementary Materials

Materials and Methods

Figs. S1 to S4

Table S1

## Materials and methods

### 1. Clinical sample collection

A mono-center matched control study with two parallel groups, one with and one without dandruff or seborrheic dermatitis was conducted. The study was approved by institutional review board at the Nanyang Technological University (IRB-2019-11-021). All volunteers were provided with written informed consent prior to enrolment in the study. Sixty-five volunteers were enrolled in the study and were classified as healthy with dandruff adherent score of 0, and D/SD with dandruff adherent score of 1.5 and above on a scale of 0 to 5. All volunteers were given a neutral shampoo and were asked to use it for 3 times a week for up to 21 days and then invited for sampling. Three types of samples were collected. 1) Scalp swab samples were collected using sterile cotton swabs soaked in sterile collection buffer (0.15 M NaCl, 0.1% Tween 20). The wet swab was rubbed on the scalp surface covering an area of 16 cm^2^. The sampling procedure was repeated 8 times in the same area and the swabs were collected in a sterile Eppendorf tube, transported in dry ice, and stored at −20°C until processing. 2) Corneum discs, made of polyester film covered with transparent adhesive, was placed on the scalp surface after trimming the hair in that area using sterile scissors and removed to collect the adherent scalp material. 4 corneum discs were applied and collected from the same area and the discs were folded and collected in individual Eppendorf tubes and stored at −20°C until further processing. For SEM analysis these samples were fixed using the scanning electron microscope (SEM) fixation buffer (2.5% glutaraldehyde in 0.1 M sodium cacodylate buffer, pH 7.4, Electron Microscopy Sciences, USA) 3) Two sets of 10 hair follicles were plucked out one by one from the scalp using sterile forceps by a trained dermatologist. One set of follicles were immersed in SEM fixation buffer and stored at 4°C until sample processing. The second set was collected in sterile tubes and stored at −20°C until DNA extraction.

### 2. Metagenomic DNA extraction, sequencing, and analysis

Total DNA was extracted from swabs and follicles for sequencing. For swab samples, after thawing on ice, the biomass was extracted from the swabs by adding 700 µl of 0.9% NaCl to the tubes containing the swabs, and vortexed for 30 seconds. The swab was then carefully removed from the liquid, pressed against the wall of the tube to dry out the excess liquid and placed in a fresh sterile Eppendorf tube. Another round of extraction was carried out as above to the same swab in the fresh tube, and the dried swab was placed in a third tube. The two tubes containing the extracts were centrifuged at 10,000 xg for 10 minutes. The supernatants were discarded and the pellets from both tubes were resuspended one by one in to 750 µl of bead solution (PowerLyzer PowerSoil DNA isolation kid, cat. no. 12855). The cotton portion of the swab was also cut and transferred to the same tube containing the biomass suspension n the bead solution. Same procedure was followed for tow swabs that were not used for sampling as controls. For the follicles, the follicle part of the plucked hair samples (10 samples) were cut from each volunteer and added to the bead tube provided by the DNA isolation kit and 750 µl of the bead solution was added. DNA extraction was performed as recommended by the kit manufacturer. The extracted DNA was dissolved in sterile nuclease-free water.

For metagenomic sequencing, library preparation was performed according to Swift Biosciences’ Accel-NGS 2S Plus DNA Library Prep kit (cat. no. 21024) protocol. The samples were sheared on a Covaris E220 to ∼450bp, following the manufacturer’s recommendation, and uniquely tagged with Swift Biosciences’ Accel-NGS 2S Dual Indexing barcodes (cat. no. 29096) to enable sample pooling for sequencing. Finished libraries were quantitated using Promega’s QuantiFluor dsDNA assay (cat. no. E4871) and the average library size was determined on an Agilent Tapestation 4200. Library concentrations were then normalized to 4nM and validated by qPCR on a QuantStudio-3 real-time PCR system (Applied Biosystems), using the Kapa library quantification kit for Illumina platforms (Cat. no. 07960573001). The libraries were then pooled at equimolar concentrations and sequenced on the Illumina HiSeq2500 platform at a read-length of 250bp paired-end. Unassembled sequencing reads were directly analysed by CosmosID-HUB multi-kingdom microbiome analysis Platform (CosmosID Inc., Germantown, MD, USA) for taxonomic profiling of the microbes. Further analyses were done using phyloseq package in R (https://joey711.github.io/phyloseq/index.html). The DNA sequence data is uploaded to NCBI (Accession No. pending).

### 3. Microbes and growth conditions

*C. acnes* ATCC 6919, and *S. epidermidis* ATCC 12228 were routinely grown on brain heart infusion (BHI) broth or with 1.5% agar (Accumedia 7116B). *M. restricta* ATCC MYA-4611 was routinely cultured in modified Dixon both or with 1.5% agar (36 g/L of malt extract, 20 g/L Dessicated oxbile, 6 g/L bacto peptone supplemented with 1% v/v tween 40, 0.2% oleic acid, and 0.2% glycerol). For biofilm and co-culture experiments, a modified BHI media (mBHI) was formulated with 37 g/L of BHI base supplemented with 0.4% v/v tween 40, 0.2% v/v oleic acid, and 0.2% glycerol. Biofilms were formed on either 24 well glass bottom plates of glass bottom 8 well chambers (Cellvis). For SEM imaging, biofilms were grown on 13 mm diameter glass coverslips in 24 Well glass plastic plates (Cellcis). Biofilms were seeded with exponentially growing cultures at an initial inoculum density of 10^5^ cell/ml. For co-culture experiments a 1:1 ratio of all the organisms were ensured as the initial seeding density. All growth experiments were conducted at 33°C. For anaerobic experiments, including growing *C. ances*, the experimental set up was incubated in sealed GENbag anaer with Anaer indicator (Biomerieux).

### 4. Scanning Electron microscopy and analysis

Clinical samples and biofilm samples fixed using the SEM fixation buffer were subsequently subjected to a series of washing steps (3 repetitions for 5 mins each) with 0.1M Cacodylate buffer pH 7.4(Electron Microscopy Sciences, USA) and distilled water (Gibco). Samples were dehydrated by incubating them with mild rocking for 10 minutes each with increasing concentrations of absolute ethanol (Merck) in water from 25% to 100% ethanol (v/v). Ethanol was sequentially replaced by increasing concentrations of hexamethyldisilazane (HMDS, Electron microscopy solutions) in ethanol and finally the samples in HMDS was dried overnight. Samples were coated 4nM platinum (Hitachi FlexSEM 1000 II) prior to SEM imaging.

### 5. Quantification of microbes

Quantification of the bacteria were carried out by resuspending the microbes in biofilm in sterile PBS (Difco). 10 x serial dilutions were carried out using sterile PBS up to 8 dilutions. The serially diluted suspensions were drop plated on BHI agar plates and grown at 33°C until colonies were counted. For *M. restricta*, cells were stained using SYTO 9 (Thermo Fisher Scientific) and the fluorescent intensity was measured using Tecan microplate reader at 485 nm excitation and 530 nm emission wavelength. A caliberation curve for cell counts (direct counting under microscope) against the SYTO 9 measurements was generated and used to calculate the cell numbers.

### 6. ATP estimation

For ATP estimations, cells or biofilms were resuspended in 1x PBS (Difco) measurements carried out using BacTiterGlo™Microbial cell viability Assay (Promega) in accordance with manufacturer’s protocol.

### 7. Aconitase assay

Aconitase activity was measured using the Aconitase assay kit (ab83459, Abcam). Briefly, cell pellets were resuspended in Aconitase preservation solution with 1/10 volume of detergent supplied and incubated on ice for 30 minutes. The suspension was then centrifuged at 20,000 xg for 10 minutes and supernatant was collected. For the measurement of aconitase activity, 50 μl of the sample was added to microplate wells followed by the addition of 200 μl of Assay buffer to each well. Enzyme activity was measured at OD_240_ 20-second intervals for 30 min at room temperature. Aconitase enzyme activity was expressed as mU per log10 cells.

### 8. Intracellular-iron estimation

Cell pellets were stored in −80°C freezer overnight and the lyophilized using a freeze dryer (Labconco FreeZone −84°C). Freeze-dried samples were digested with 400 uL 70% nitric acid and 200 uL 30% hydrogen peroxide for 3 days at room temperature. After 3 days, 600 uL of digested sample was aliquoted and diluted with 3.4mL of LC-MS grade water. The diluted samples were filtered with 0.2um syringe filter. An aliquot of 2mL of filtered sample was mixed with 4mL of LC-MS grade water to achieve a final concentration of 1.98% of nitric acid. Intracellular iron concentrations were measued using ICP-MS analysis conducted in Central Environmental Science and Engineering Laboratory (CEE, NTU, Singapore), and are expressed in pg/log10 cells.

### 9. Fluorescent in situ hybridisation

Biofilms were gently washed with 1X PBS (Gibco) and then fixed with 4% (w/v) paraformaldehyde at room temperature for an hour. Following fixation, 0.25% (w/v) low temperature gelling agarose was added to the wells and allowed to solidify at 4°C. Biofilms were gently washed again with 1X PBS. Dehydration was performed with consecutive additions of 50%, 80% and 96% (v/v) ethanol, incubating for 5 minutes each. Wells were dried at 46°C for 30 minutes and stored at 4°C until further processing.

For cell lysis, lysis buffer [1,000,000 U/mL lysozyme, 0.05M EDTA, 0.1M Tris/HCl (pH 8.0), dH_2_O] in combination with 0.4 U/µL (v/v) RNase inhibitor (Applied Biosystems) was added to the dehydrated biofilms and incubated in a moist chamber at 37°C for an hour. Biofilms were then washed 3 times with distilled water followed by consecutive additions of 50%, 80% and 96% (v/v) ethanol for 3 minutes each. Chambers were placed at 46°C for drying of biofilms.

Fluorescent probes specific for *S. epidermidis*, *C.acnes* and *M. restricta* (M&M_Tabel 1) were employed for hybridization to the target sequences in the organism. Hybridization buffer [0.9M NaCl, 0.02M Tris/HCl, 30% Formamide, 0.1% SDS, ddH2O] was loaded to the wells together with the probes and incubated at 46°C for 24 hours. Following incubation, hybridization buffer was aspirated from the wells, and biofilms were rinsed once with wash buffer [0.215M NaCl, 0.02M Tris/HCl, 0.005M EDTA, ddH_2_O] that was prewarmed to 48°C. Biofilms were incubated with the wash buffer and washed 3 times at 15-minute intervals. A final rinse with distilled water is performed and the biofilms were subsequently dried at 46°C. Chambers were stored in the dark at 4°C until microscope analysis.

### 10. Confocal microscopy and analysis

FISH biofilm samples were imaged using confocal laser scanning microscope (Carl Zeiss LSM 780, Germany) with 20X and 63X (oil immersion) objective lenses. Z-stack images of the biofilms were obtained using Zen Microscopy Software. 488 nm, 518 nm, and 633 nm lasers were used for excitation of Alexa Fluor 488, Cy3 and Cy5, respectively. Image processing was performed on Imaris 9.0.2.

### 11. Gas chromatography

For the estimation of short chain fatty acids (acetic acid, propionic acid, butyric acid, and valeric acid), cell free supernatants were collected from bacterial and fungal cultures and were centrifuged at 10,000 xg for 5 minutes, followed by filtration using 0.2 u pore size filters. Cell free supernatants were acidified with formic acid (0.1% final concentration) and analysed by GC-FID using a DB-FFAP column (Agilent).

### 12. RNA extraction

Samples were concentrated using centrifugation (10,000 xg for 10 minutes) and added with 300ul of TRIzol™(Invitrogen) with rocking for 5 minutes, resuspended in trizol and stored at −80c. For RNA extraction, thawed samples were lysed by bead beating (Lysing Matrix E, 2 mL, MP Biomedicals) with three repeated cycles at 6,.5 m/s for 45 seconds. Subsequently, 200ul of Chloroform (Merck) was added per 1ml of trizol followed by centrifugation at 12,000 xg for 10mins. RNA was extracted using chloroform: isoamyl alcohol (24:1) (Merck), followed by precipitation by isopropanol (Merck). The final RNA pellet was washed with 70% ethanol followed by air drying for 2 hrs followed by dissolving in nuclease free water (Invitrogen). RNA cleanups were performed using TURBO DNase (Invitrogen) and RNA Clean & Concentrator™-5 (Zymo Research), the concentration of purified RNA was determined using Qubit™ RNA HS Assay Kit (Invitrogen) according to manufacturer’s protocols.

### 13. Quantitative gene expression analysis

cDNA was synthesised using SuperScript™ III First-Strand Synthesis System (Invitrogen). Briefly, a cDNA synthesis mix was prepared as follows. One hundred ng of RNA was used a template for the reaction mixture that consisted of 4ul 10x RT buffer, 8ul 25mM Mgcl_2_, 4ul 0.1M DDT,2ul RNase OUT (40U/ul), and 2ul SuperScript™ III RT (200U/ul). A 1:1 mixture of 20ul of cDNA synthesis mix and 20 ul RNA-primer mix that contained 2 ul of 50 ng/ul random hexamers and 10 mM dnTPs mix and nuclease free water, was prepared in 0.2ml strip tubes for synthesis. This mixture was incubated at 65℃ for 5 minutes and then on ice for 1 minute. The tubes were then subjected to incubation at 25℃ for 10 minutes, 50℃ for 50 minutes, and 85℃ for 5 minutes in a thermocycler (Mastercycler® Pro, Eppendorf). After cooling on ice, the reaction mixture was collected by brief centrifugation. Finally, 2ul of RNase H was added to the samples and, were incubated at 37℃ for 20 minutes.

Gene specific qPCR primers were designed (M&M_Table 2) and commercially synthesized by IDT and validated. qRT-PCR were performed in a 20ul reaction volume containing 10ul KAPA SYBR® FAST 2x mix (Merck), 0.4 μL ROX high (KAPA SYBR® FAST, Merck), 0.4 μL forward primer, 0.4 μL reverse primer and 6.8 μL autoclaved filtered water in StepOnePlus Real-Time PCR Systems (Applied Biosystems). Gene expression was calculated as expression fold change using double-delta Ct method.

### 14. Transcriptomic sequencing

RNA library preparation was performed using Illumina’s TruSeq® Stranded Total RNA Library Prep, following the manufacturer’s protocol with 50 ng of the total RNA as input. RNA was subjected to fragmentation (5 min at 94 °C) and reverse transcription to obtain double-stranded DNA with fragment size in the range between 120 and 250 bp. The fragmented dsDNA was subjected to end repair, adapter ligation, and PCR amplification to generate sequencing-ready library. Quality of the library was checked using Agilent D1000ScreenTape. Sequencing was performed using the Illumina® HiSeq 4000 (Paired-End) Cluster kit on the cBot 2 System, generating paired end 150 bp reads. Quality of pair-ended reads was assessed using FastQC v0.11.7 (Andrews, S. (2010). FastQC A Quality Control Tool for High Throughput Sequence Data. Available at: http://www.bioinformatics.babraham.ac.uk/projects/fastqc/). rRNA reads were discarded using SortMeRNA v4.2.0(Kopylova, E., Noe, L., and Touzet, H., 2012) and the remaining rRNA reads were trimmed using Trimmomatic v0.38 (Bolger, A. M., Lohse, M., & Usadel, B. (2014). Trimmomatic: A flexible trimmer for Illumina Sequence Data. Bioinformatics, btu170.) and mapped against the reference genomes (GenBank) using HISAT2 v2.2.1 (Kim, D., Paggi, J.M., Park, C. et al. Graph-based genome alignment and genotyping with HISAT2 and HISAT-genotype. Nat Biotechnol 37, 907–915 (2019)). Aligned data were sorted with SAMTools v1.13(Li, H., Handsaker, B., Wysoker, A., Fennell, T., Ruan, J., Homer, N., et al. (2009). The sequence alignment/map format and SAMtools. Bioinformatics 25, 2078– 2079. doi: 10.1093/bioinformatics/btp352) and then used as input for the feature counts function (Yang Liao, Gordon K Smyth and Wei Shi. featureCounts: an efficient general-purpose program for assigning sequence reads to genomic features. Bioinformatics, 30(7):923-30, 2014) from the Rsubread v2.4.3 package (Liao Y, Smyth GK, Shi W (2019). “The R package Rsubread is easier, faster, cheaper and better for alignment and quantification of RNA sequencing reads.” Nucleic Acids Research, 47, e47. doi: 10.1093/nar/gkz114.) to generate a matrix of annotated genes with their corresponding raw counts. An average of 93.25% reads were successfully mapped to the reference genome. The count data were then analyzed to look for differential gene expression levels and statistical significance using DESeq2 v1.30.1 (Love MI, Huber W, Anders S (2014). “Moderated estimation of fold change and dispersion for RNA-seq data with DESeq2.” Genome Biology, 15, 550. doi: 10.1186/s13059-014-0550-8). Genes with absolute value of Log2 fold change (Log2FC) ≥1.5 and adjusted p-value <=0.05 were considered as differential expressed genes (DEGs) in comparative analysis. RNA sequencing data is uploaded to NCBI (Accession no. pending).

### 15. Metabolomics

Untargeted metabolomics analysis of the samples was carried out using capillary-electrophoresis time of flight mass spectroscopy (CE-TOFMS, Human Metabolome Technologies inc.). Samples were prepared according to the procedure and reagents provided by the service provider. Briefly, cells or biofilms were concentrated using centrifugation (10,000 xg for 10 minutes), followed by washing the pellet using sterile Milli-Q water and resuspended in 1.6 ml of ethanol and ultrasonicated for 30 seconds. An ‘internal standard solution’ provided by the service provider was added to the lysate in methanol and the metabolites were isolated from cell debris by centrifugation coupled with filtration as specified by the service provider and sent for metabolomics analysis in dry ice.

### 16. Anti-dandruff formulation and clinical study

A mono-center, randomized, matched control, double blinded, clinical study with 57 volunteers exhibiting moderate to severe dandruff (total dandruff score > 4.5 in 0 to 5 scale) was conducted. Volunteers were randomly classified into treatment and control groups. Treatment group was provided with a hydroalcoholic lotion as the vehicle containing 1% sodium propionate and 3% ascorbyl glucoside. The control group was provided with the vehicle formulation. The volunteers were asked to use the formulation every day on their scalp with the usage of a neutral shampoo 2 to 3 times a week for up to 28 days. Scalp swab samples were collected (as described above) from the volunteers on Day 0 and Day 28 of the study period. These samples were subjected to DNA extraction followed by quantification of *Malassezia*, *Cutibacterium*, and *Staphylococcus* using quantitative real time PCR using genus specific probes and primers (M&M_Table 3). PCR reactions were performed in 10 ul reaction volume with SsoADV Univer Probes Supermix 2500 Rx (Bio-Rad). 0.4 ul of the DNA sample was used in the reaction together wth 400 nM of each primer. PCR was performed with initial denaturation for 2 minutes followed by 40 cycels of 95°C for 10 seconds and 55°C (60°C for *M. restricta*) for 5 seconds. Copy numbers were calculated using the Biorad CFX Manger program by single threshold method using standard curves established using 7 known amounts of the purified DNA projects (1 to 10^6^ copies/reaction).

### 17. Statistical analysis

Statistical analyses were conducted in R and in MS Excel software. Correlation analyses were performed using Pearson correlation test and evaluated using the associated p-values. Students t-test was used to evaluate the significance of difference between two sets of data. p-value less than 0.05 was considered as statistically significant.

**M&M_Table 1.**
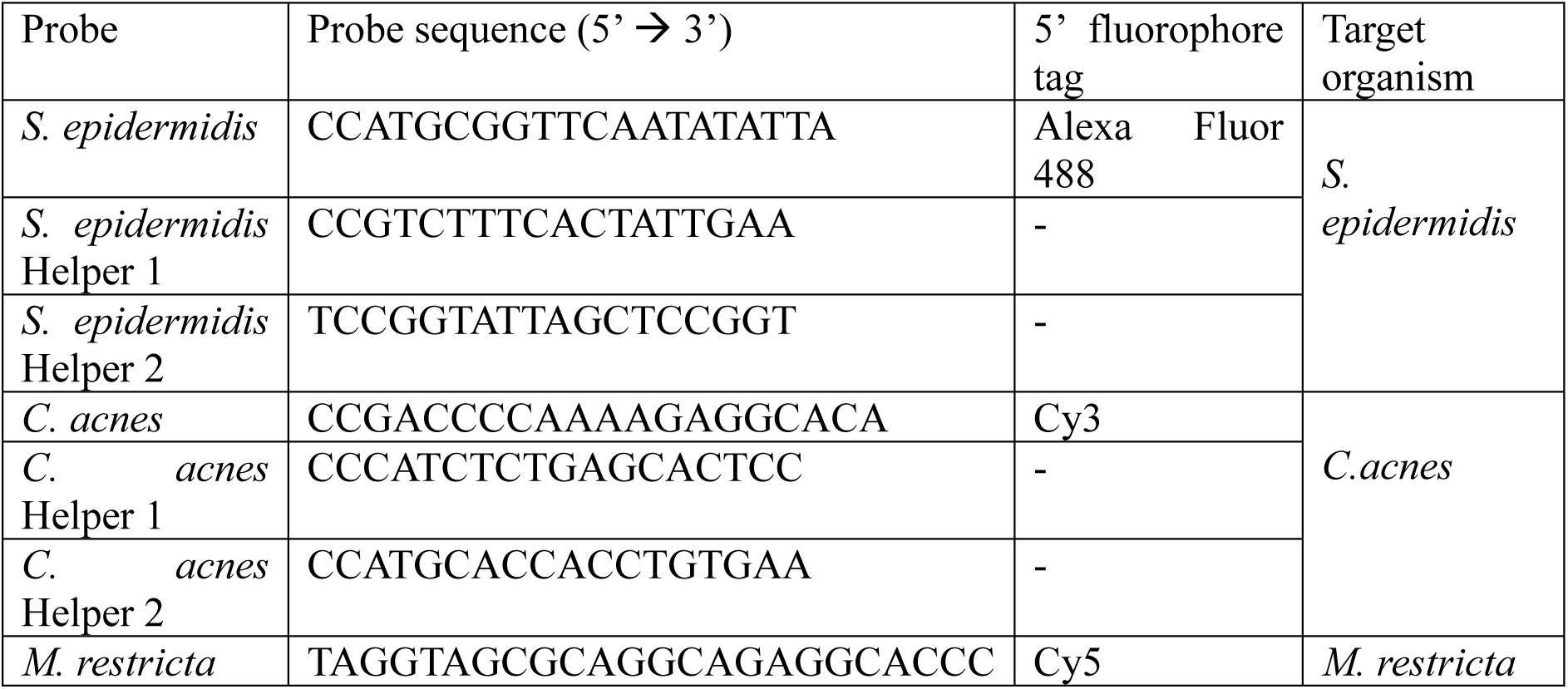
Species specific probe sequences used for FISH.

**M&M_Table 2.**
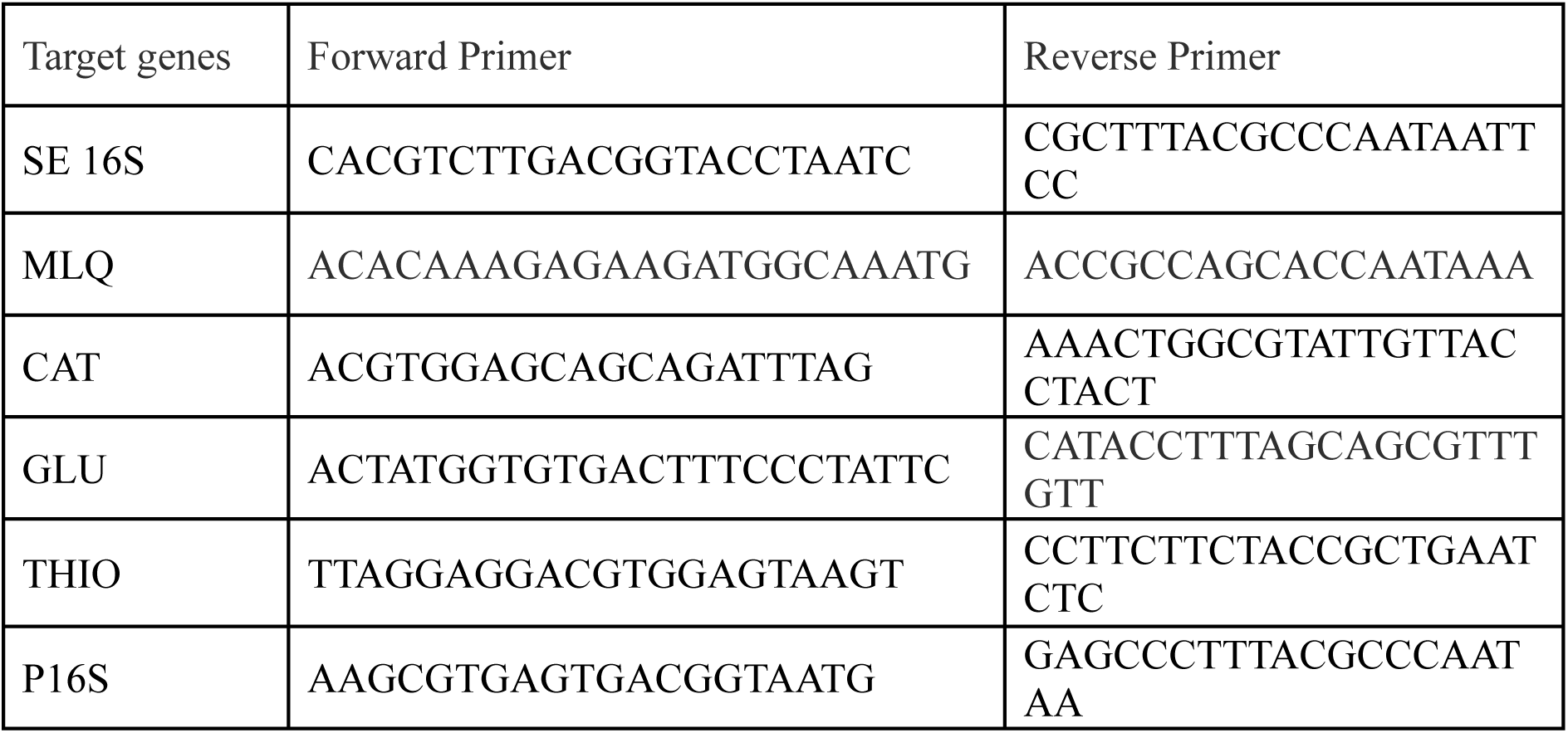
Primers used for gene expression analysis.

**M&M_Table 3.**
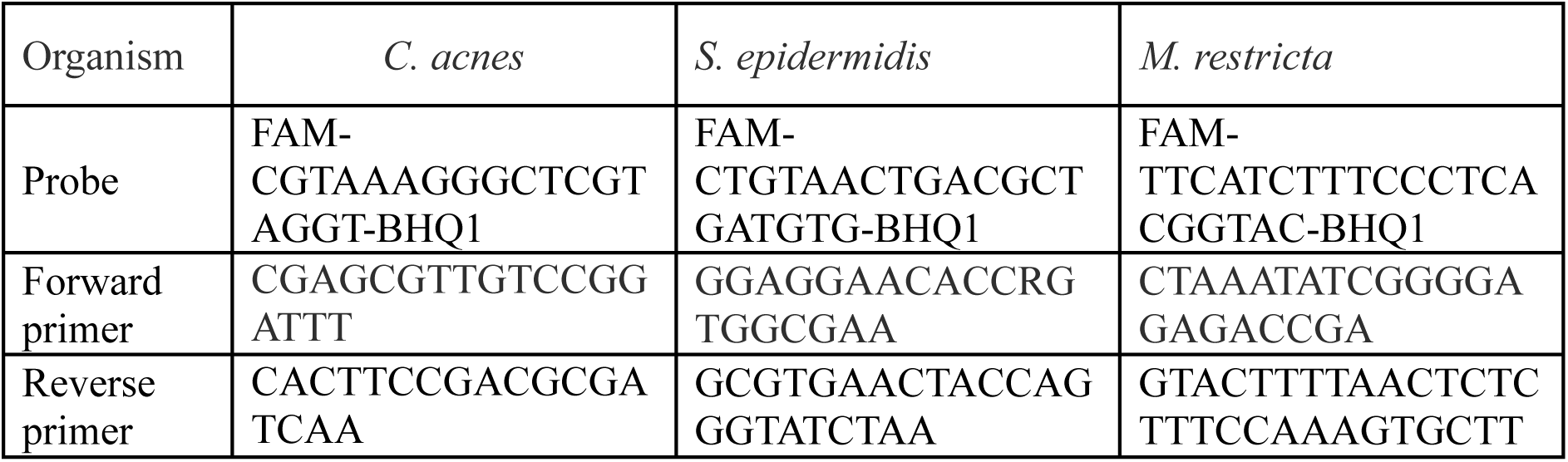
Primers and probes used for quantification of the microbes in clinical samples.

