## Supplementary figures for "Decoding scalp health and microbiome dysbiosis in dandruff"

**Fig S1.**

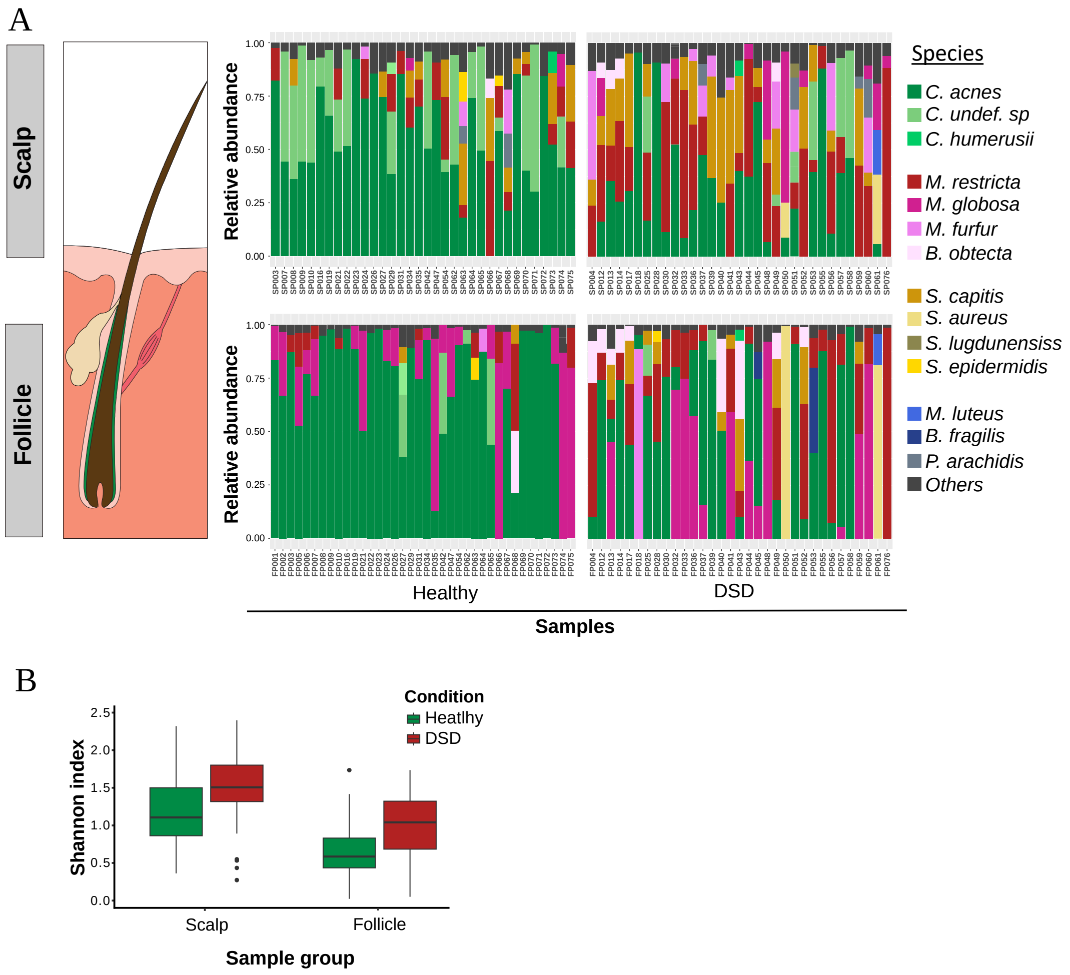
A
Scalp
Follicle
Relative abundance
1.00
0.75
0.50
0.25
0.00
Healthy
DSD
Samples
Species
C. acnes
C. undef. sp
C. humerusii
M. restricta
M. globosa
M. furfur
B. obtecta
S. capitis
S. aureus
S. lugdunensiss
S. epidermidis
M. luteus
B. fragilis
P. arachidis
Others
B
Condition
Heatlhy
DSD
Shannon index
2.5
2.0
1.5
1.0
0.5
0.0
Scalp
Follicle
Sample group

**Fig S4**

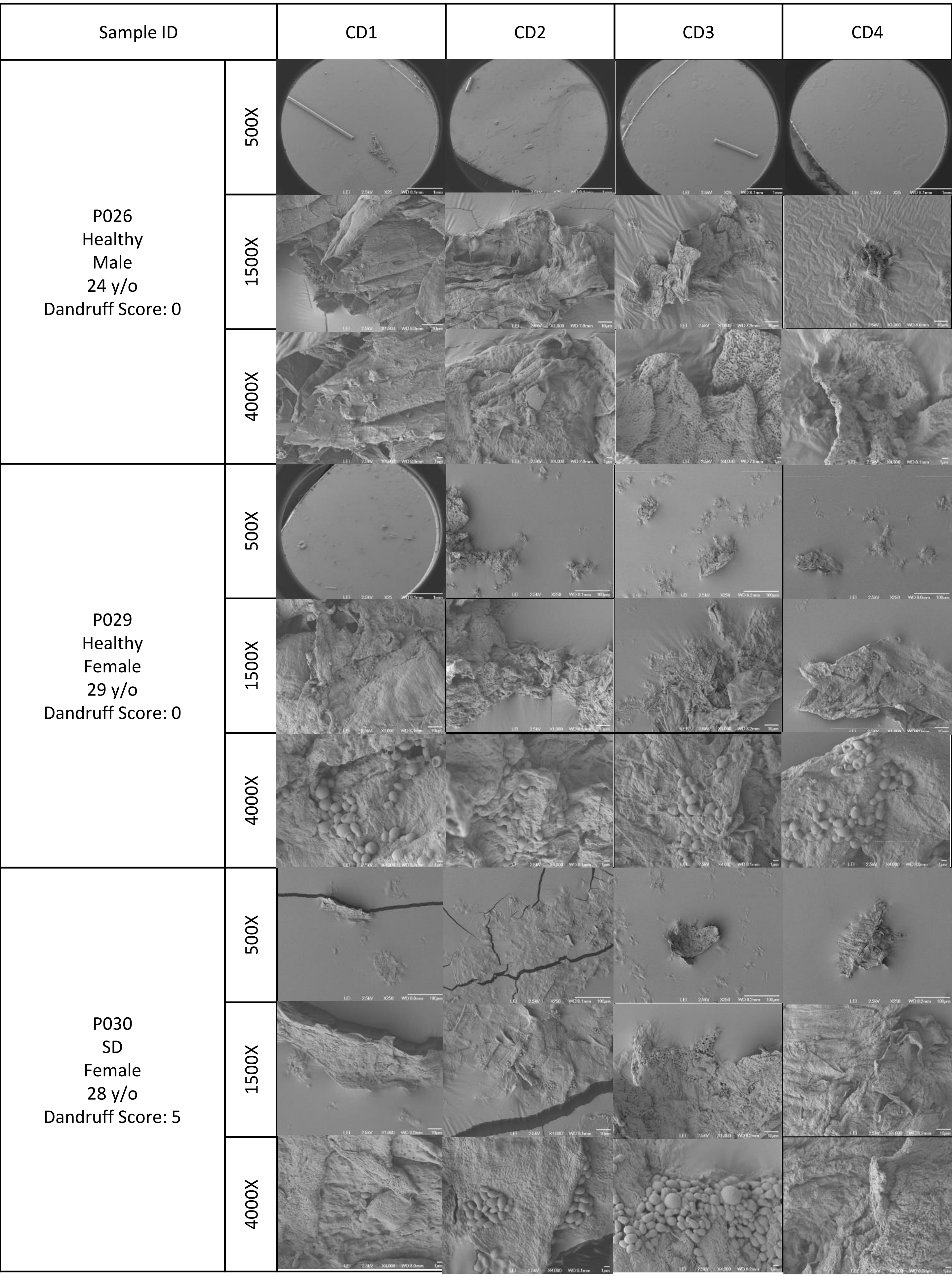
Sample ID
CD1
CD2
CD3
CD4
P026
Healthy
Male
24 y/o
Dandruff Score: 0
P029
Healthy
Female
29 y/o
Dandruff Score: 0
P030
SD
Female
28 y/o
Dandruff Score: 5
500X
1500X
4000X

| Sample ID | | CD1 | CD2 | CD3 | CD4 |
|---|---|---|---|---|---|
| P032<br>Dandruff<br>Female<br>40 y/o<br>Dandruff Score: 2.825 | 500X | 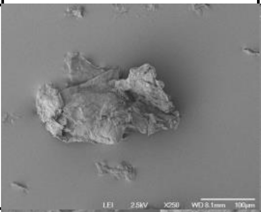 | 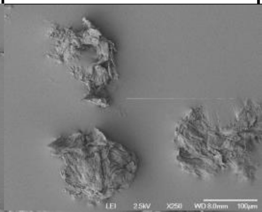 | 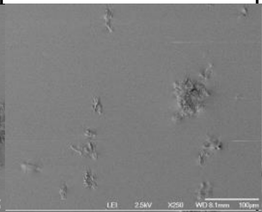 | 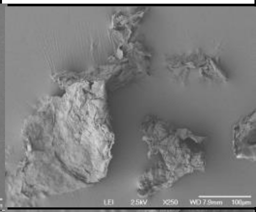 |
| | 1500X | 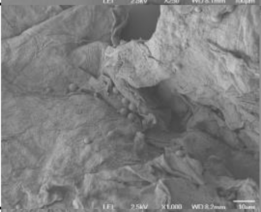 | 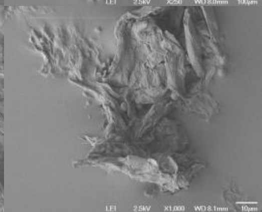 | 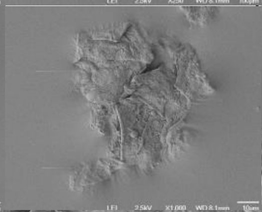 | 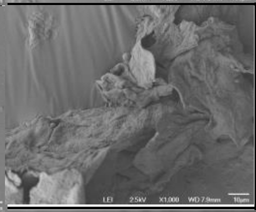 |
| | 4000X | 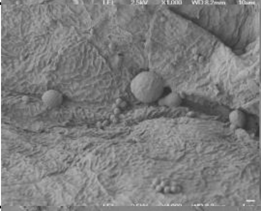 | 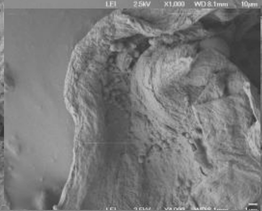 | 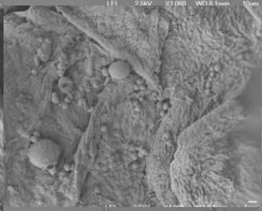 | 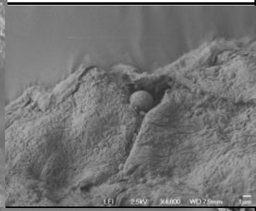 |
| P033<br>Dandruff<br>Male<br>39 y/o<br>Dandruff Score: 2.125 | 500X | 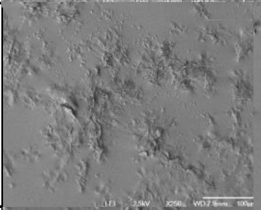 | 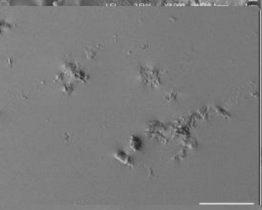 | 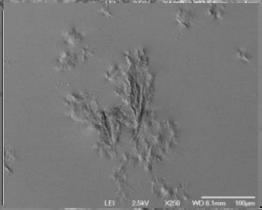 | 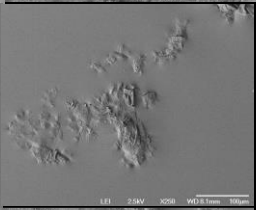 |
| | 1500X | 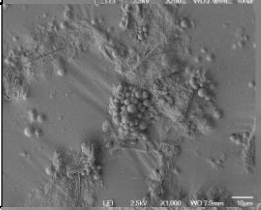 | 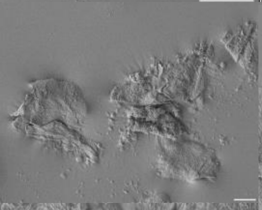 | 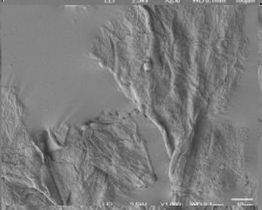 | 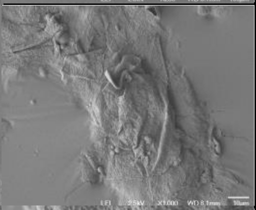 |
| | 4000X | 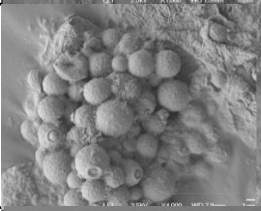 | 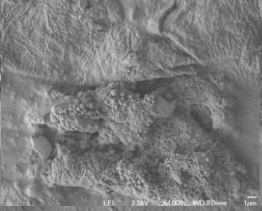 | 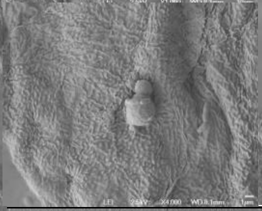 | 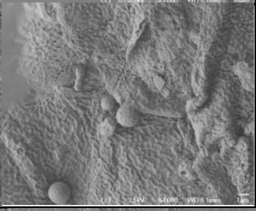 |
| P034<br>Healthy<br>Female<br>25 y/o<br>Dandruff Score: 0 | 500X | 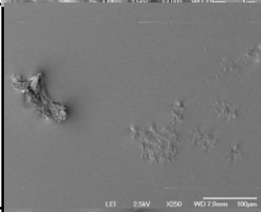 | 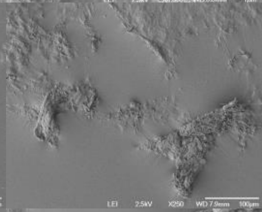 | 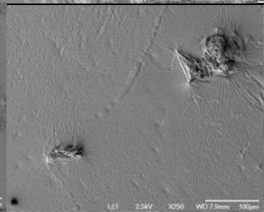 | 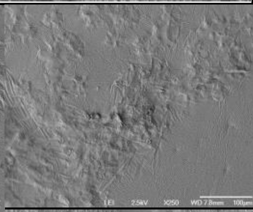 |
| | 1500X | 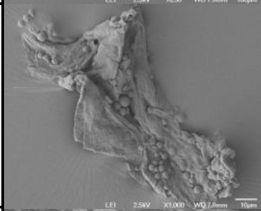 | 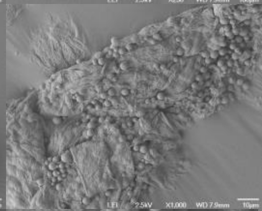 | 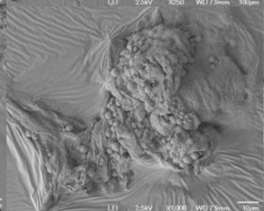 | 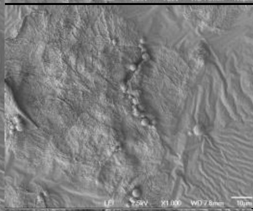 |
| | 4000X | 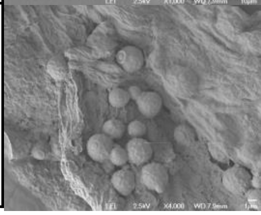 | 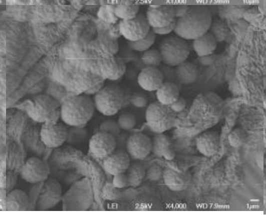 | 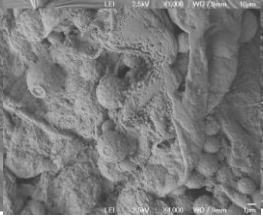 | 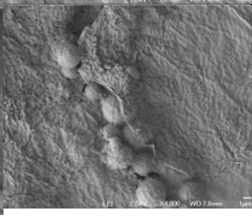 |

**Fig S2.**

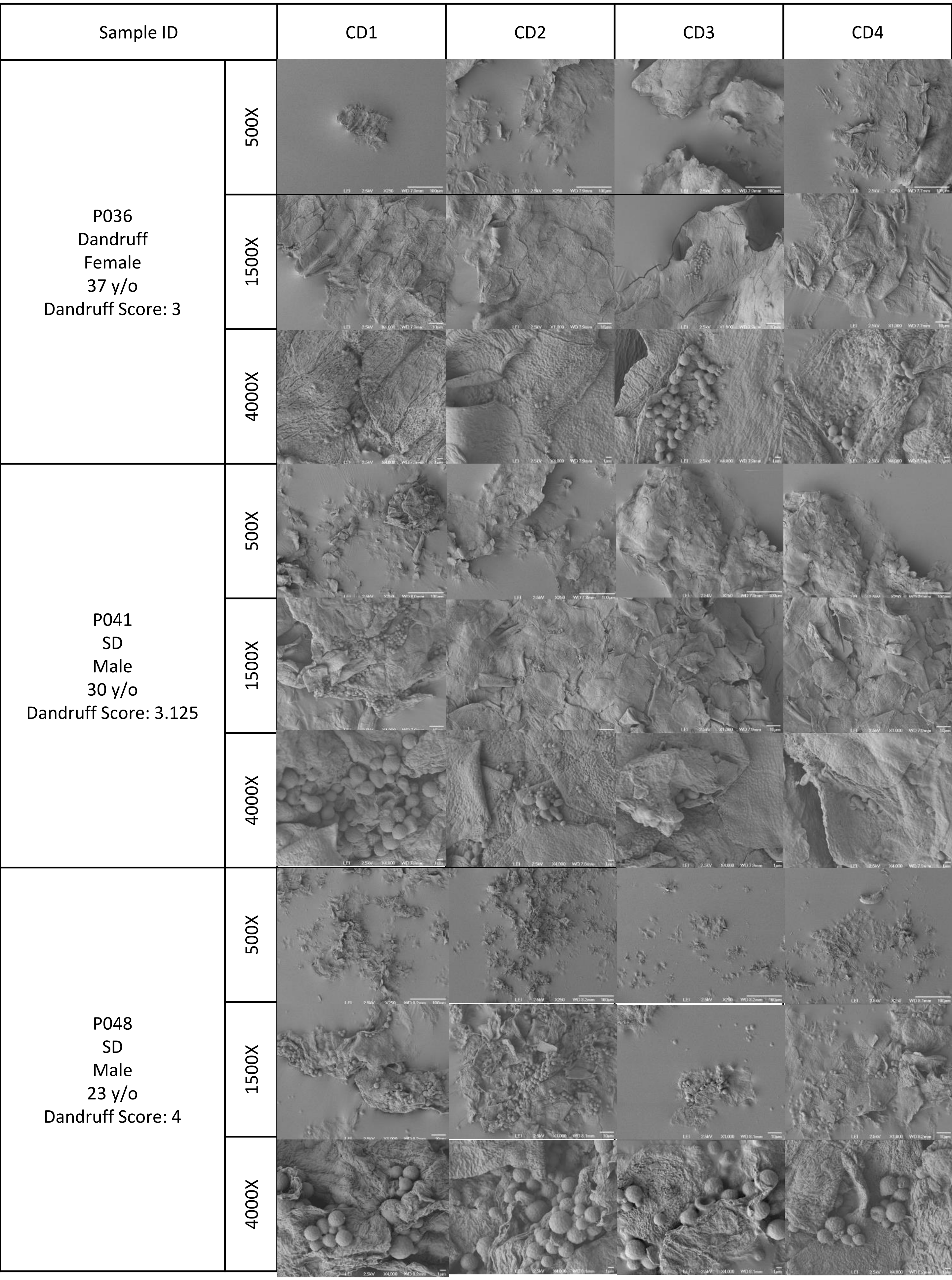
Sample ID
CD1
CD2
CD3
CD4
P036
Dandruff
Female
37 y/o
Dandruff Score: 3
500X
1500X
4000X
P041
SD
Male
30 y/o
Dandruff Score: 3.125
500X
1500X
4000X
P048
SD
Male
23 y/o
Dandruff Score: 4
500X
1500X
4000X

| Sample ID | | CD1 | CD2 | CD3 | CD4 |
|---|---|---|---|---|---|
| P062<br>Healthy<br>Female<br>35 y/o<br>Dandruff Score: 0 | 500X | 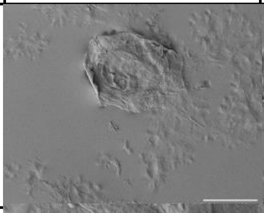 | 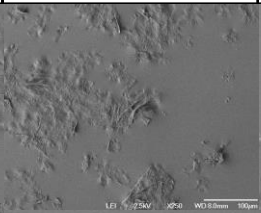 | 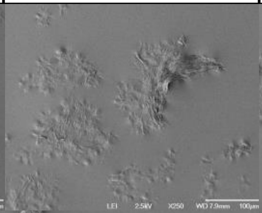 | 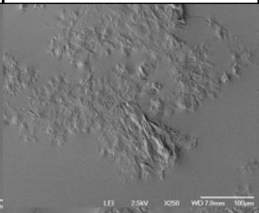 |
| | 1500X | 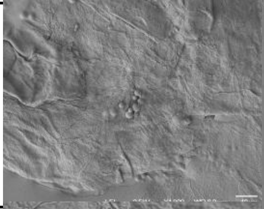 | 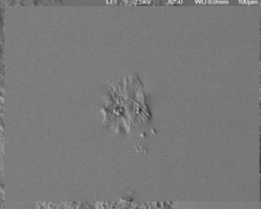 | 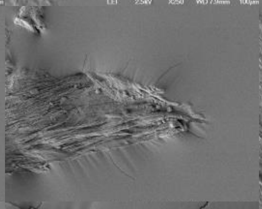 | 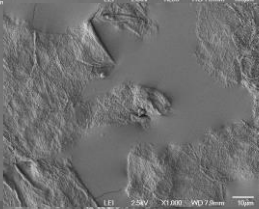 |
| | 4000X | 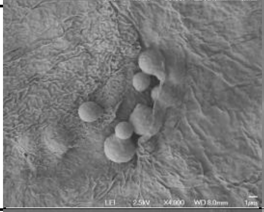 | 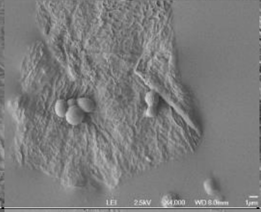 | 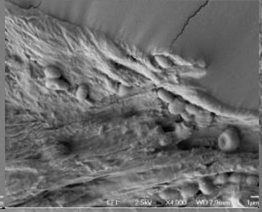 | 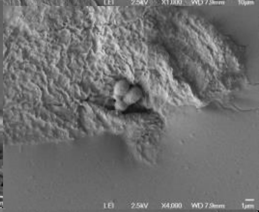 |
| P064<br>Healthy<br>Female<br>33 y/o<br>Dandruff Score: 0 | 500X | 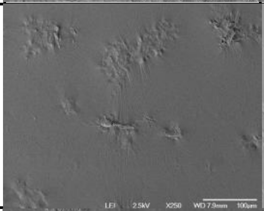 | 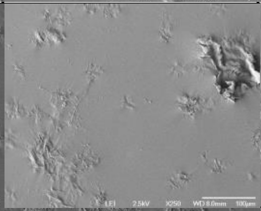 | 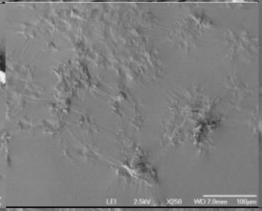 | 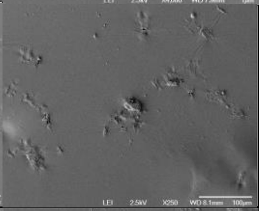 |
| | 1500X | 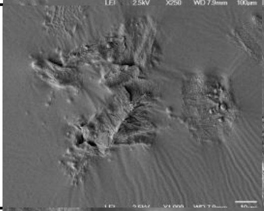 | 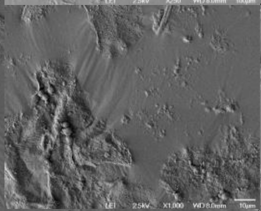 | 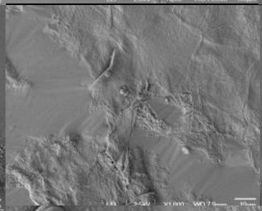 | 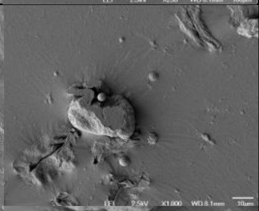 |
| | 4000X | 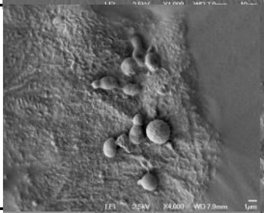 | 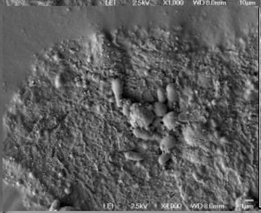 | 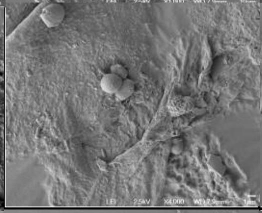 | 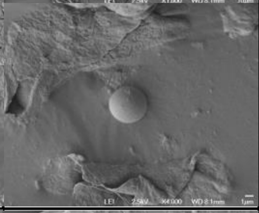 |
| P065<br>Healthy<br>Female<br>48 y/o<br>Dandruff Score: 0 | 500X | 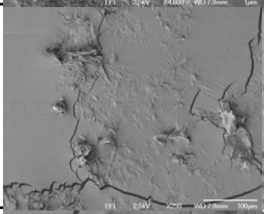 | 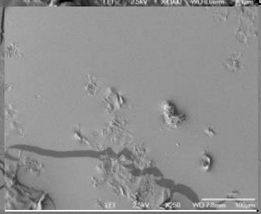 | 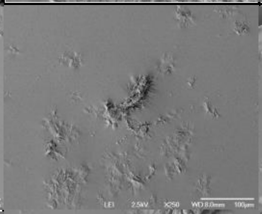 | 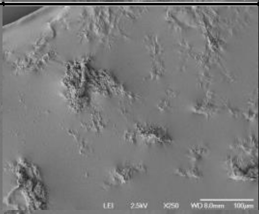 |
| | 1500X | 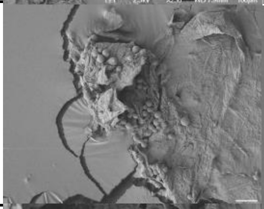 | 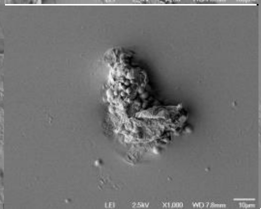 | 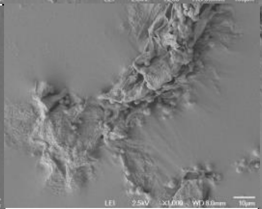 | 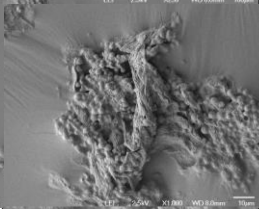 |
| | 4000X | 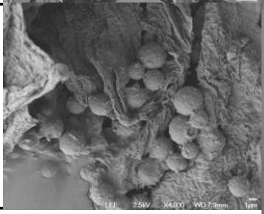 | 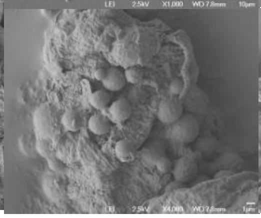 | 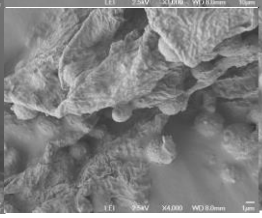 | 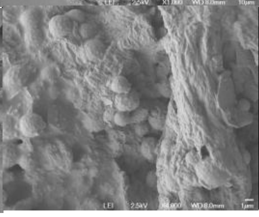 |

**Fig S3.**

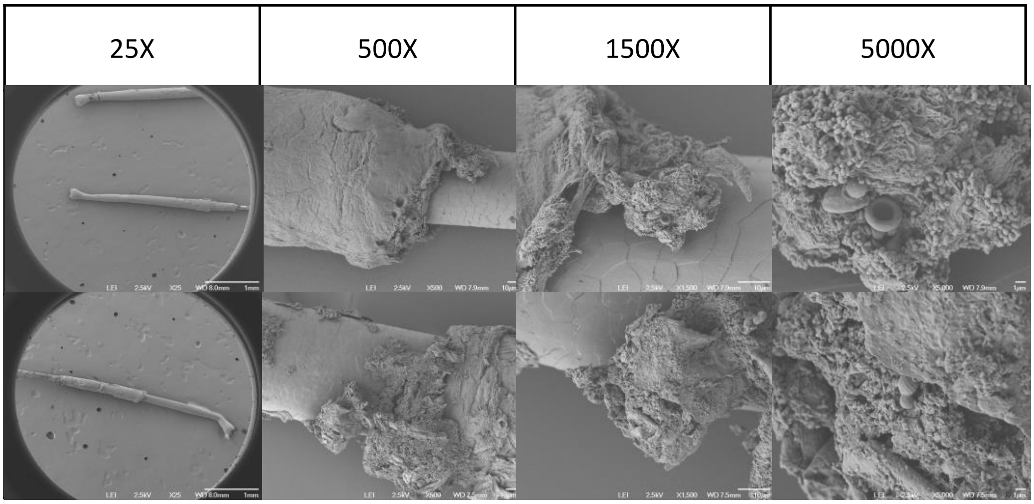

| Sample ID | | Full | Top | Middle | Root |
|---|---|---|---|---|---|
| P008<br>Healthy<br>Dandruff Score: 0 | Sample 1 | | | | |
| | Sample 2 | | | | |
| | Sample 3 | | | | |
| P009<br>Healthy<br>Dandruff Score: 0 | Sample 1 | | | | |
| | Sample 2 | | | | |
| | Sample 3 | | | | |
| P010<br>Healthy<br>Dandruff Score: 0 | Sample 1 | | | | |
| | Sample 2 | | | | |
| | Sample 3 | | | | |

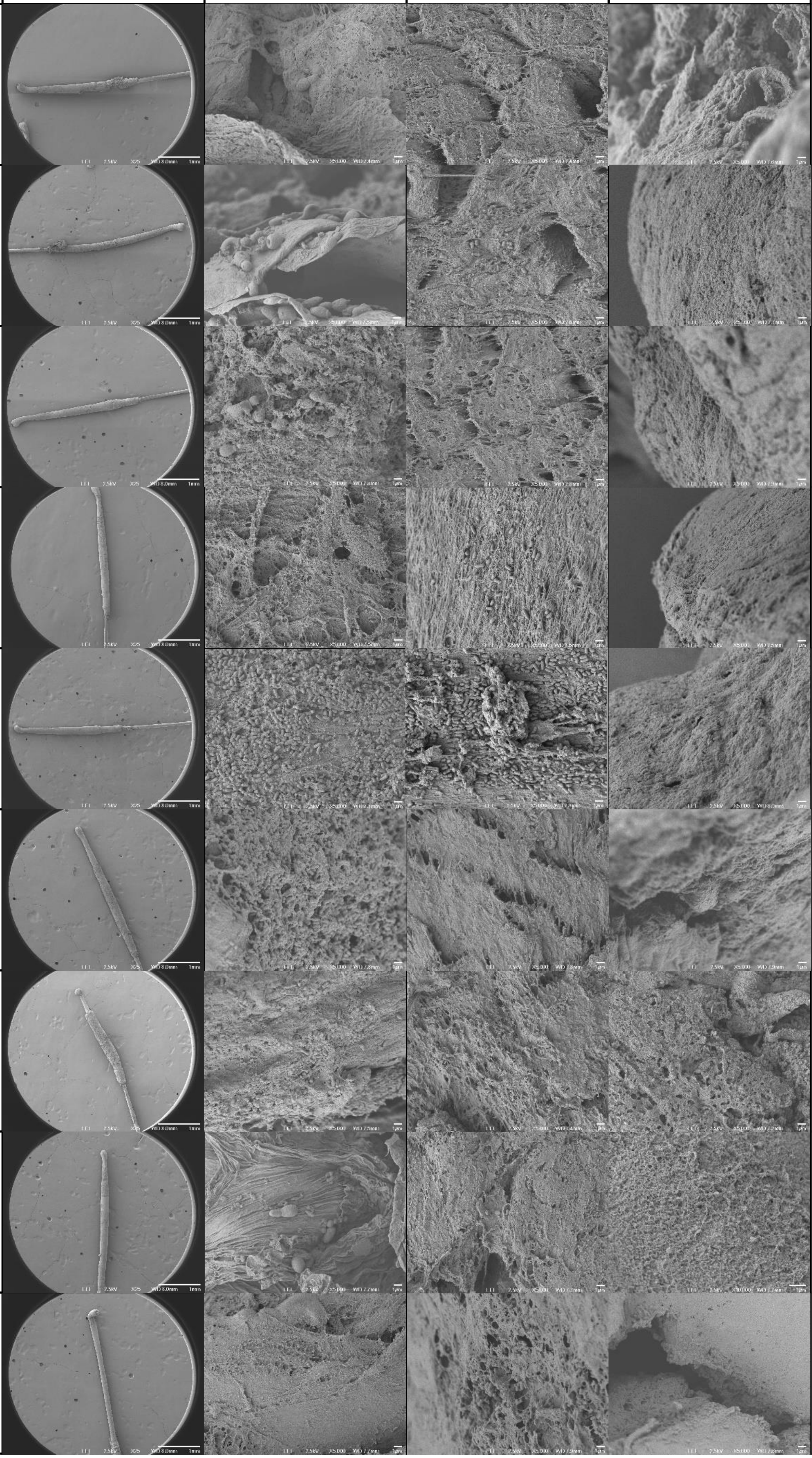

| Sample ID | | Full | Top | Middle | Root |
|---|---|---|---|---|---|
| P012<br>SD<br>Dandruff Score: 4 | Sample 1 | 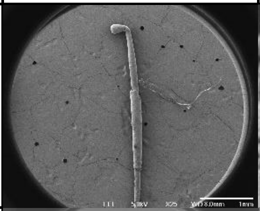 | 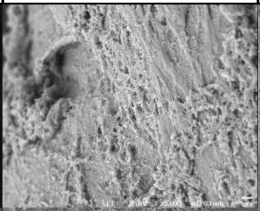 | 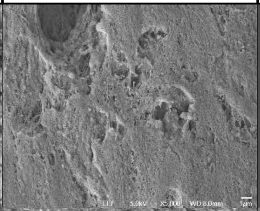 | 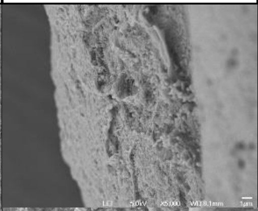 |
| | Sample 2 | 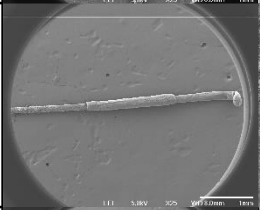 | 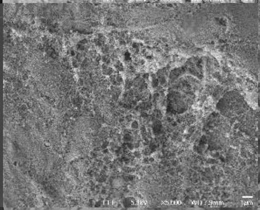 | 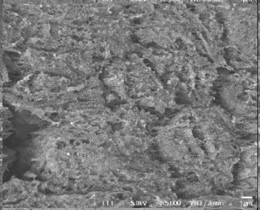 | 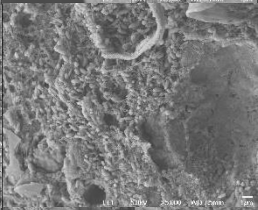 |
| | Sample 3 | 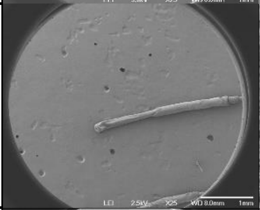 | 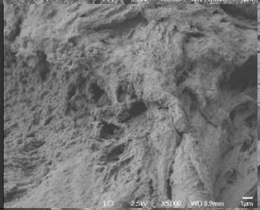 | 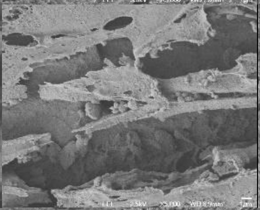 | 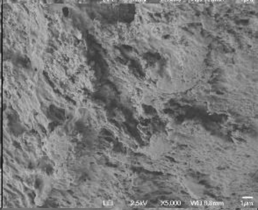 |
| P013<br>SD<br>Dandruff Score: 4.125 | Sample 1 | 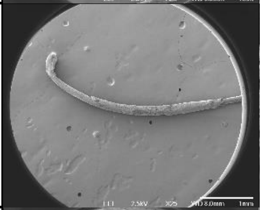 | 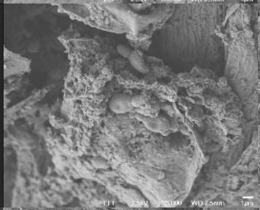 | 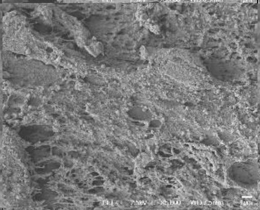 | 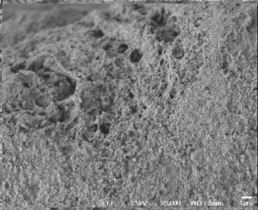 |
| | Sample 2 | 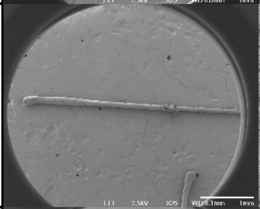 | 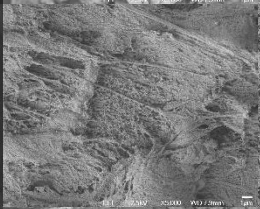 | 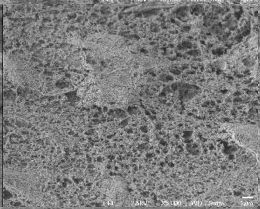 | 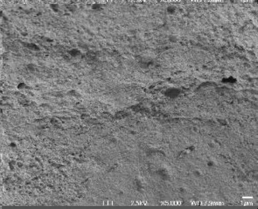 |
| | Sample 3 | 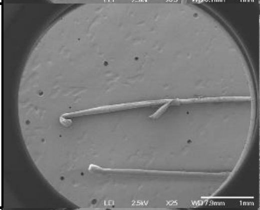 | 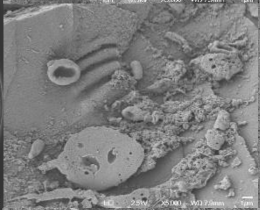 | 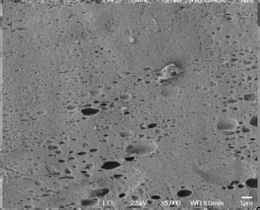 | 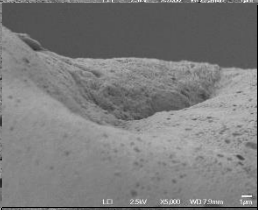 |
| P014<br>SD<br>Dandruff Score: 4 | Sample 1 | 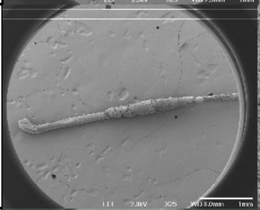 | 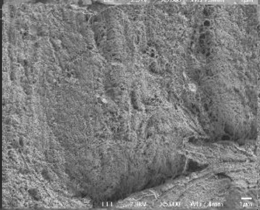 | 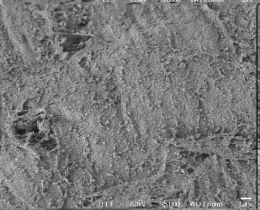 | 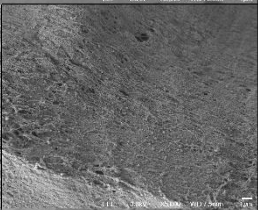 |
| | Sample 2 | 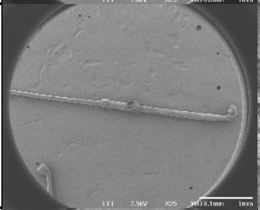 | 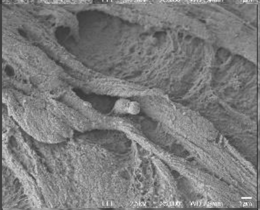 | 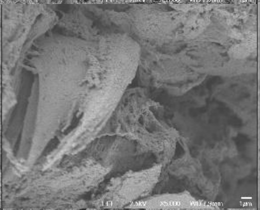 | 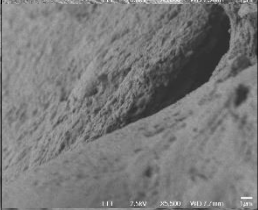 |
| | Sample 3 | 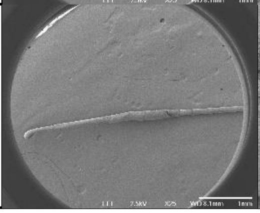 | 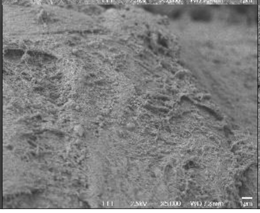 | 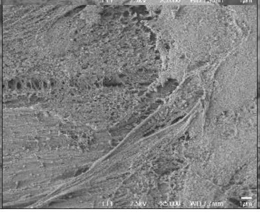 | 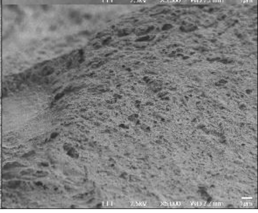 |

| Sample ID | | Full | Top | Middle | Root |
|---|---|---|---|---|---|
| P016<br>Healthy<br>Dandruff Score: 0 | Sample 1 | 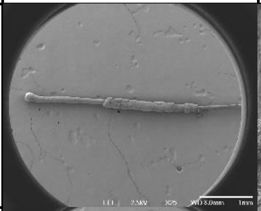 | 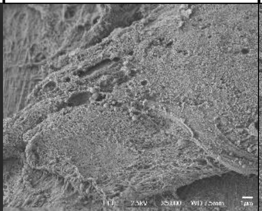 | 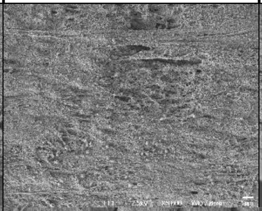 | 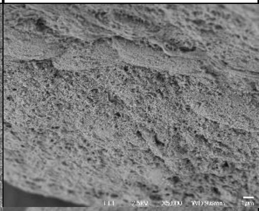 |
| | Sample 2 | 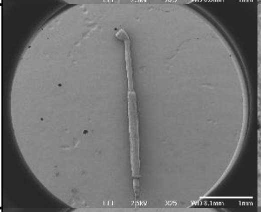 | 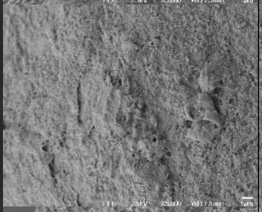 | 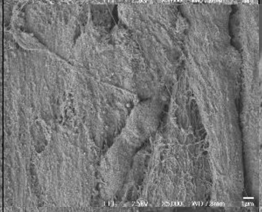 | 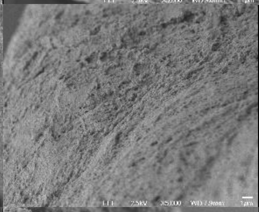 |
| | Sample 3 | 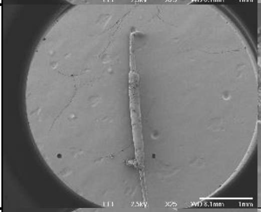 | 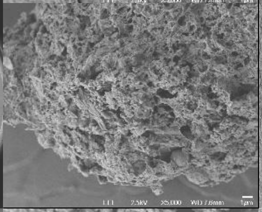 | 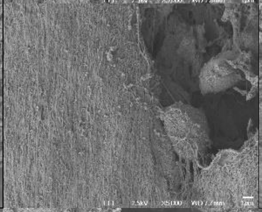 | 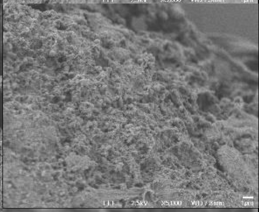 |
| P017<br>SD<br>Dandruff Score: 4.625 | Sample 1 | 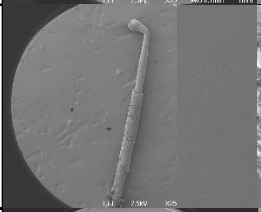 | 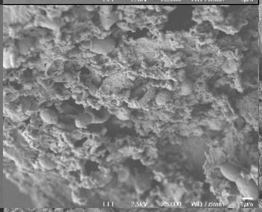 | 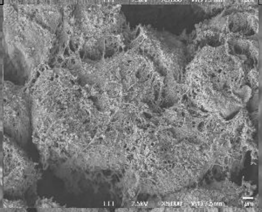 | 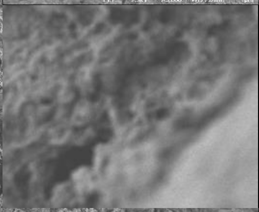 |
| | Sample 2 | 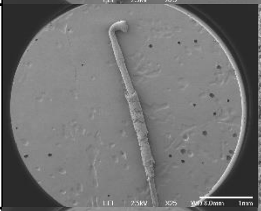 | 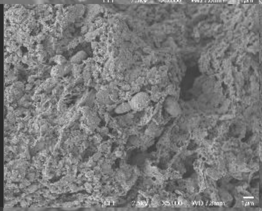 | 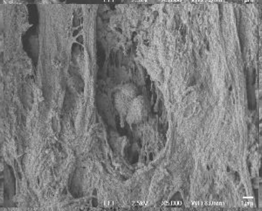 | 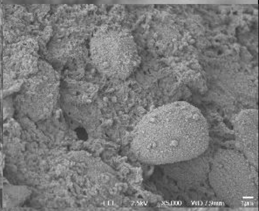 |
| | Sample 3 | 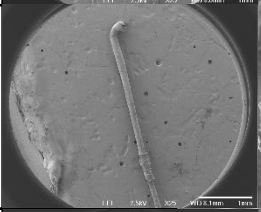 | 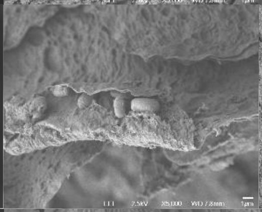 | 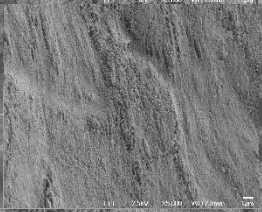 | 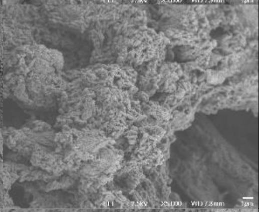 |
| P018<br>SD<br>Dandruff Score: 3.625 | Sample 1 | 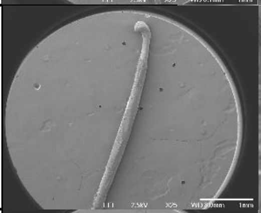 | 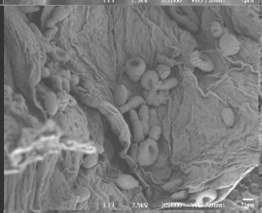 | 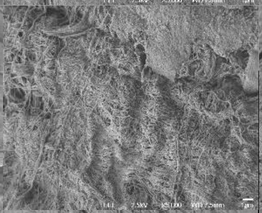 | 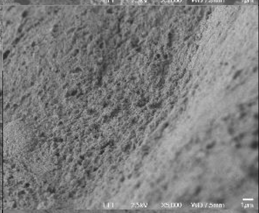 |
| | Sample 2 | 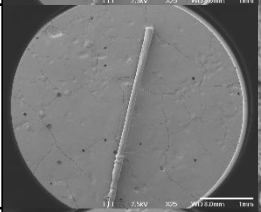 | 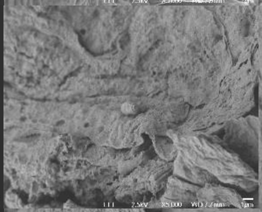 | 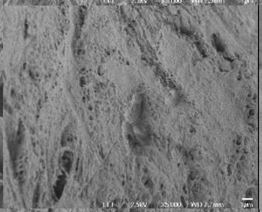 | 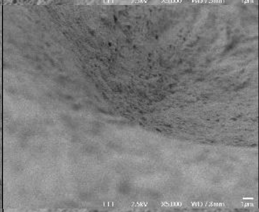 |
| | Sample 3 | 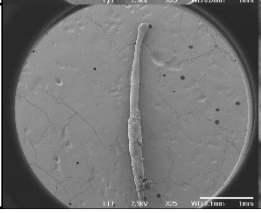 | 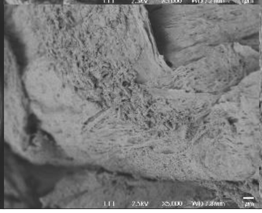 | 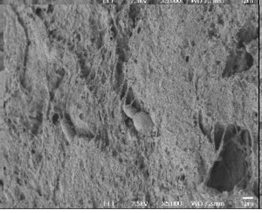 | 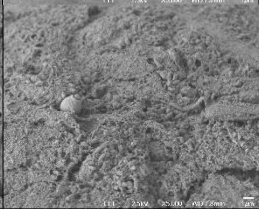 |

| Sample ID | | Full | Top | Middle | Root |
|---|---|---|---|---|---|
| P019<br>Healthy<br>Dandruff Score: 0 | Sample 1 | 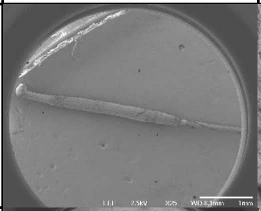 | 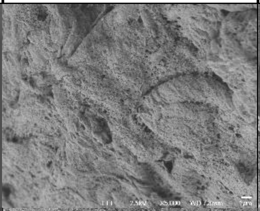 | 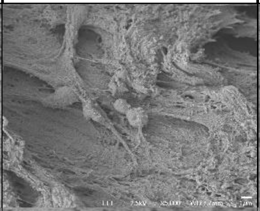 | 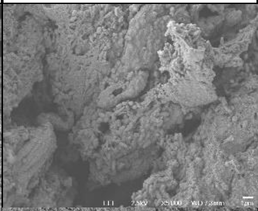 |
| | Sample 2 | 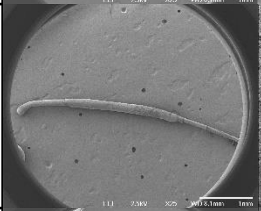 | 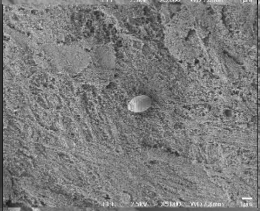 | 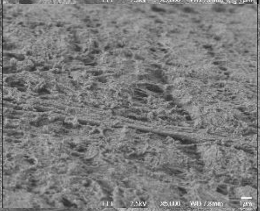 | 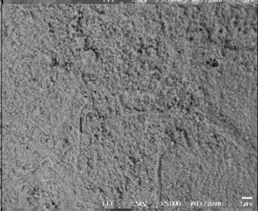 |
| | Sample 3 | 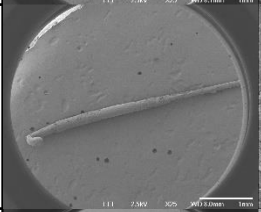 | 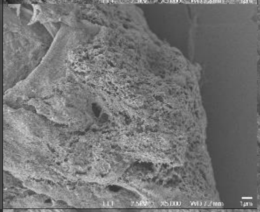 | 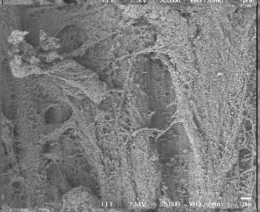 | 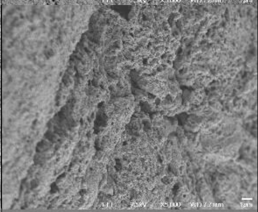 |
| P021<br>Healthy<br>Dandruff Score: 0 | Sample 1 | 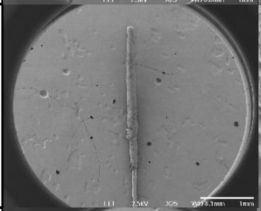 | 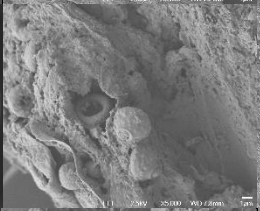 | 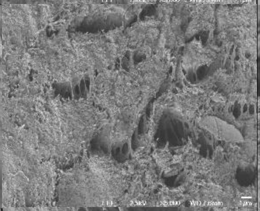 | 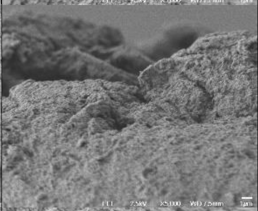 |
| | Sample 2 | 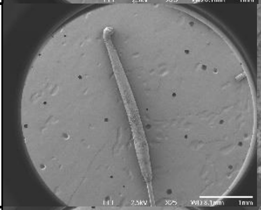 | 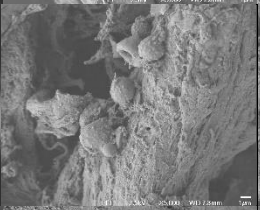 | 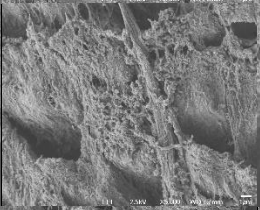 | 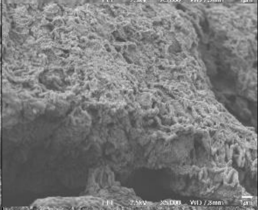 |
| | Sample 3 | 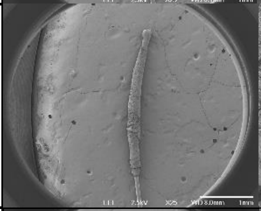 | 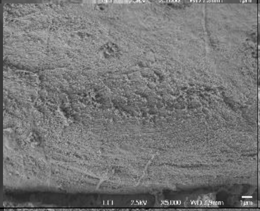 | 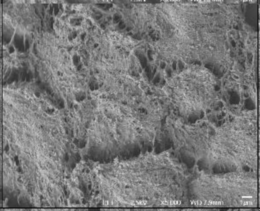 | 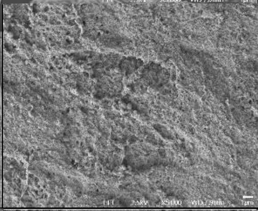 |
| P022<br>Healthy<br>Dandruff Score: 0 | Sample 1 | 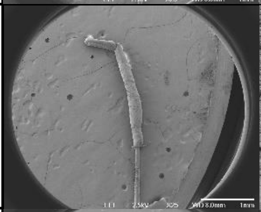 | 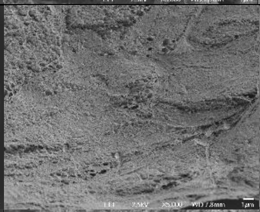 | 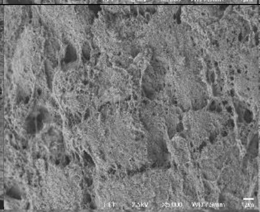 | 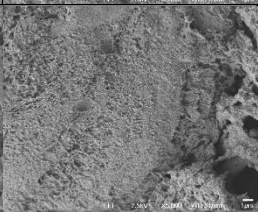 |
| | Sample 2 | 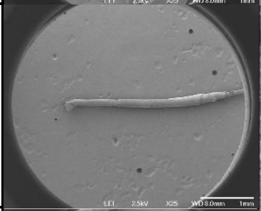 | 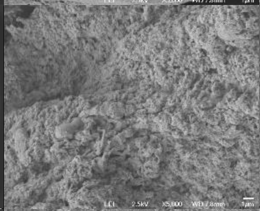 | 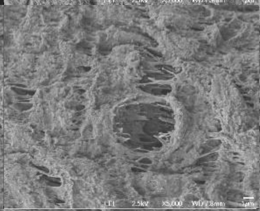 | 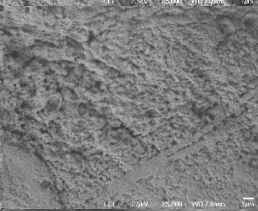 |
| | Sample 3 | 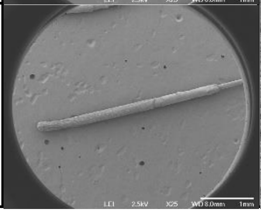 | 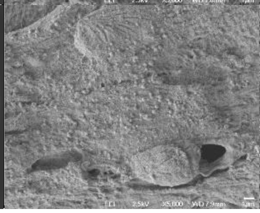 | 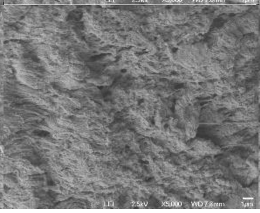 | 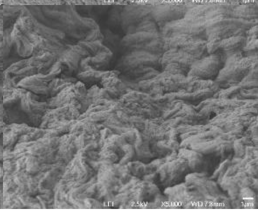 |

| Sample ID | | Full | Top | Middle | Root |
|---|---|---|---|---|---|
| P023<br>Healthy<br>Dandruff Score: 0 | Sample 1 | 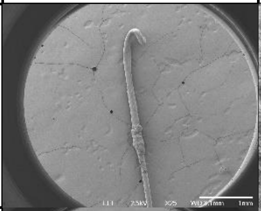 | 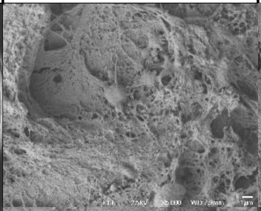 | 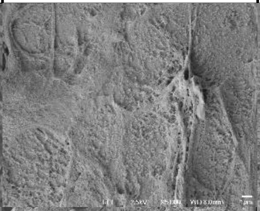 | 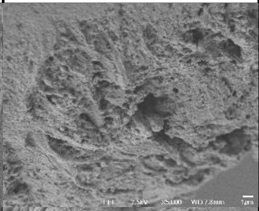 |
| | Sample 2 | 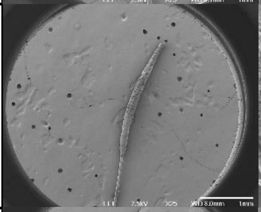 | 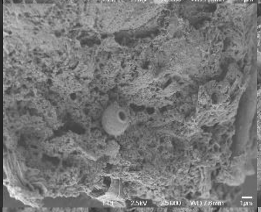 | 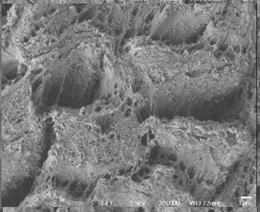 | 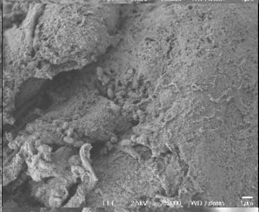 |
| | Sample 3 | 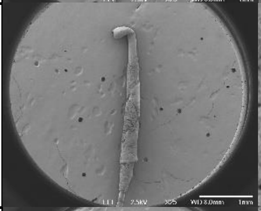 | 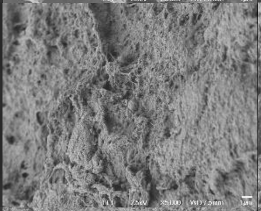 | 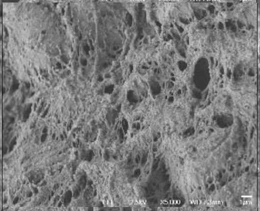 | 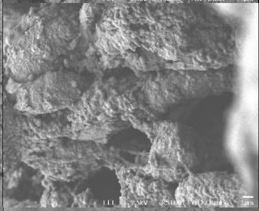 |
| P024<br>Healthy<br>Dandruff Score: 0 | Sample 1 | 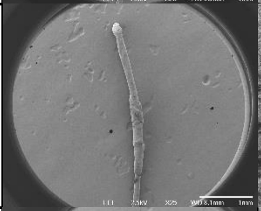 | 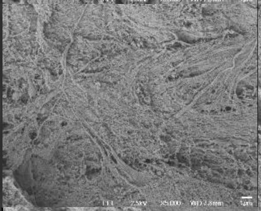 | 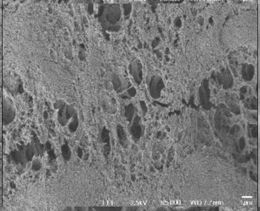 | 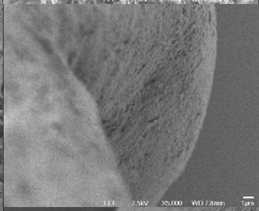 |
| | Sample 2 | 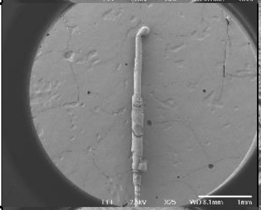 | 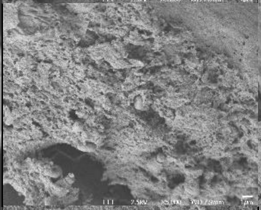 | 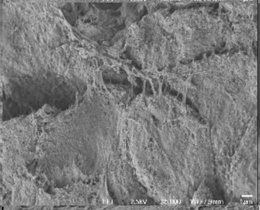 | 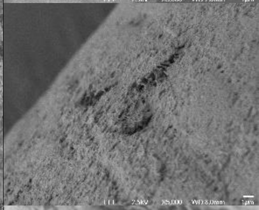 |
| | Sample 3 | 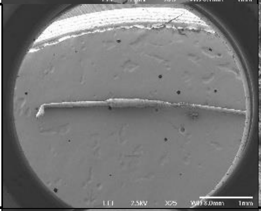 | 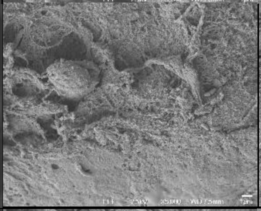 | 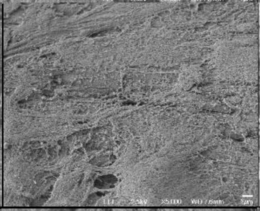 | 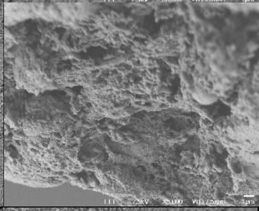 |
| P025<br>SD<br>Dandruff Score: 4.625 | Sample 1 | 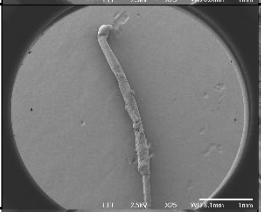 | 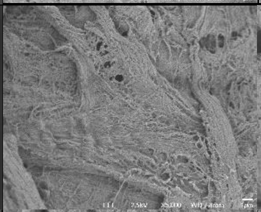 | 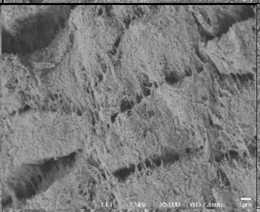 | 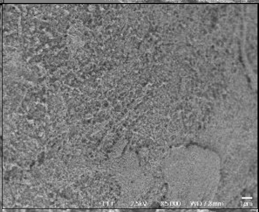 |
| | Sample 2 | 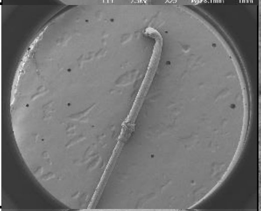 | 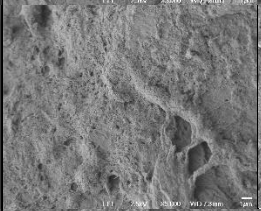 | 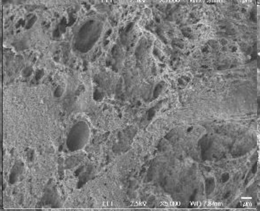 | 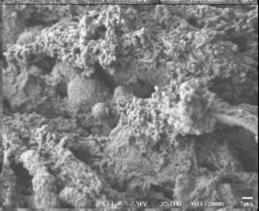 |
| | Sample 3 | 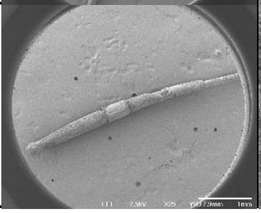 | 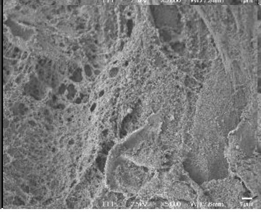 | 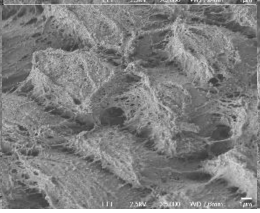 | 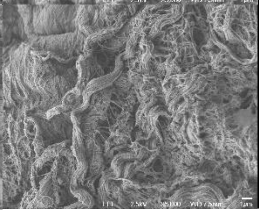 |

| Sample ID | | Full | Top | Middle | Root |
|---|---|---|---|---|---|
| P026<br>Healthy<br>Dandruff Score: 0 | Sample 1 | 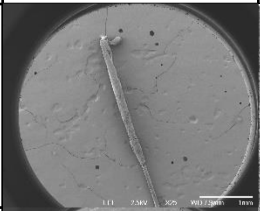 | 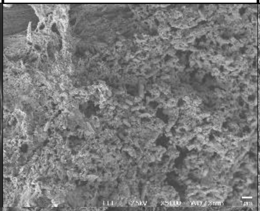 | 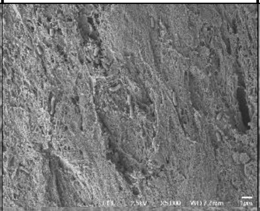 | 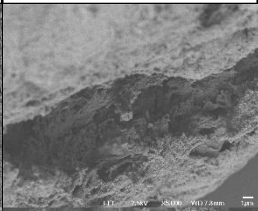 |
| | Sample 2 | 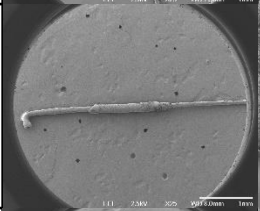 | 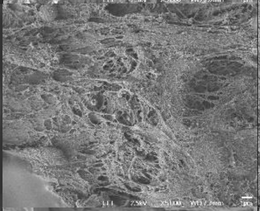 | 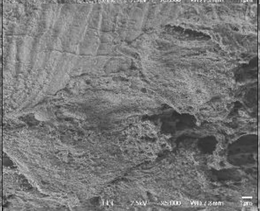 | 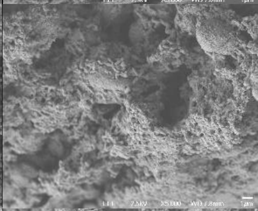 |
| | Sample 3 | 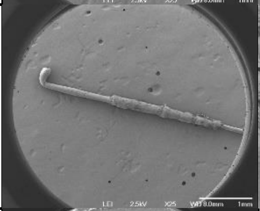 | 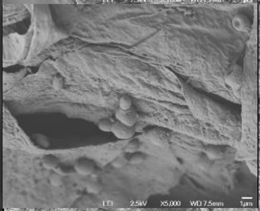 | 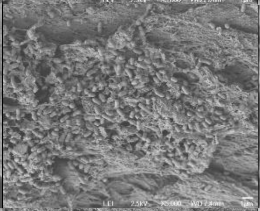 | 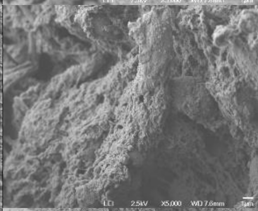 |
| P028<br>Dandruff<br>Dandruff Score: 2.25 | Sample 1 | 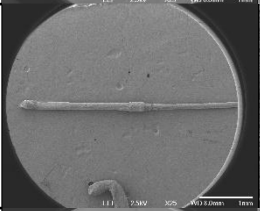 | 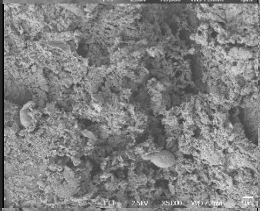 | 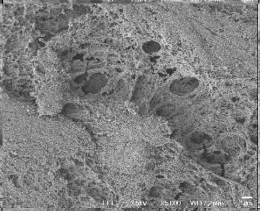 | 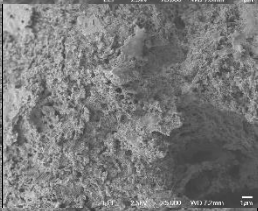 |
| | Sample 2 | 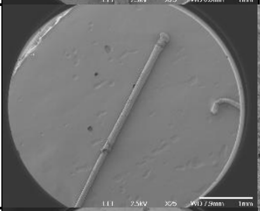 | 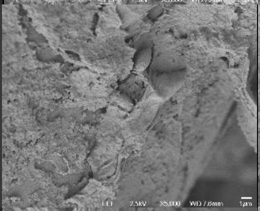 | 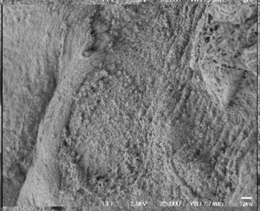 | 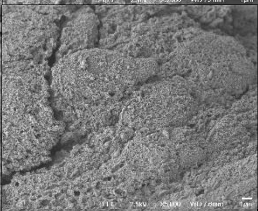 |
| | Sample 3 | 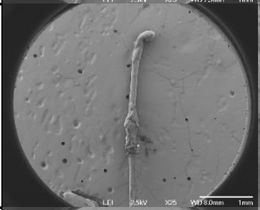 | 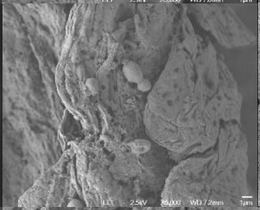 | 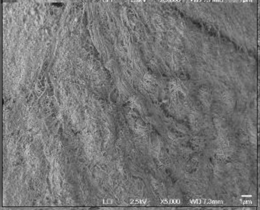 | 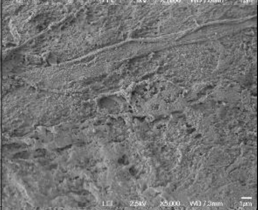 |
| P029<br>Healthy<br>Dandruff Score: 0 | Sample 1 | 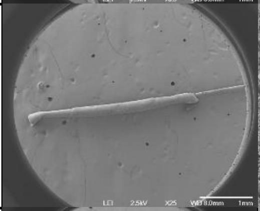 | 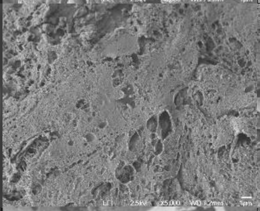 | 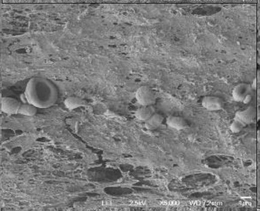 | 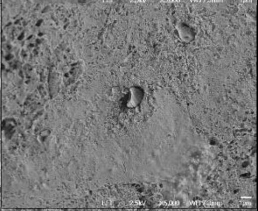 |
| | Sample 2 | 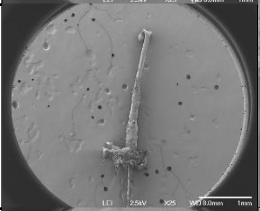 | 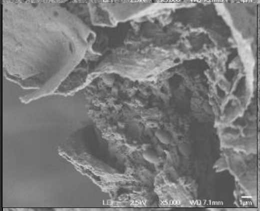 | 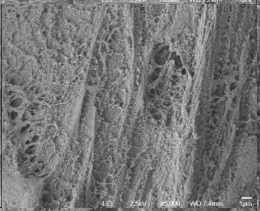 | 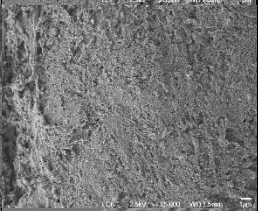 |
| | Sample 3 | 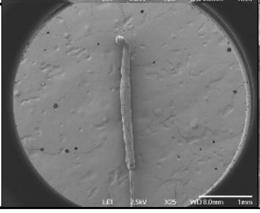 | 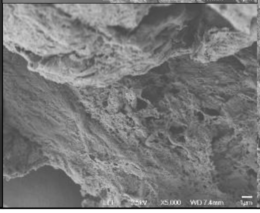 | 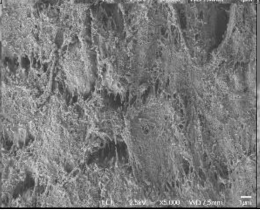 | 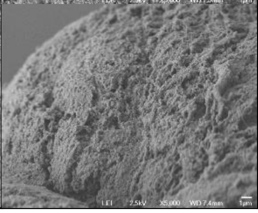 |

| Sample ID | | Full | Top | Middle | Root |
|---|---|---|---|---|---|
| P030<br>SD<br>Dandruff Score: 5 | Sample 1 | 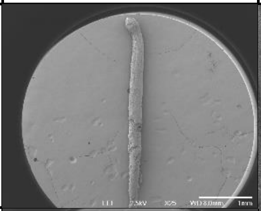 | 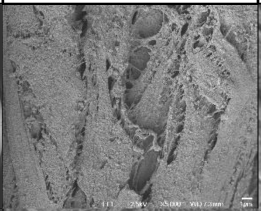 | 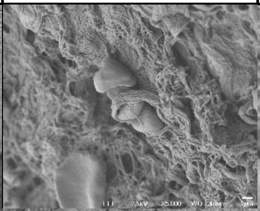 | 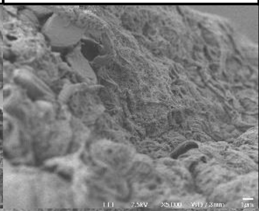 |
| | Sample 2 | 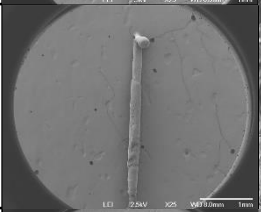 | 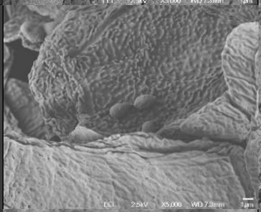 | 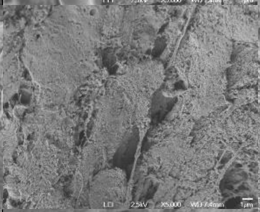 | 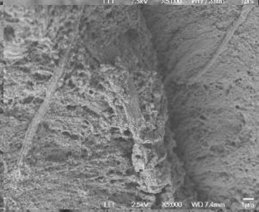 |
| | Sample 3 | 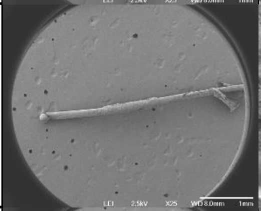 | 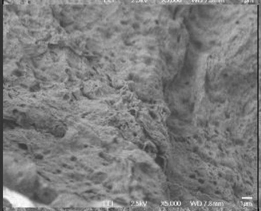 | 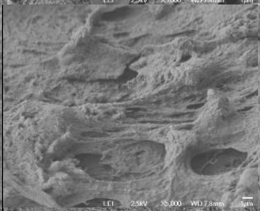 | 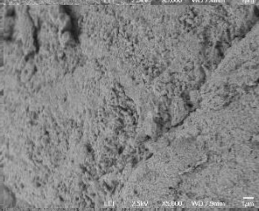 |
| P031<br>Healthy<br>Dandruff Score: 0 | Sample 1 | 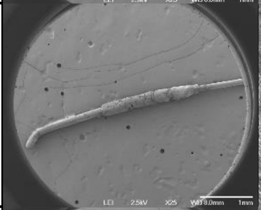 | 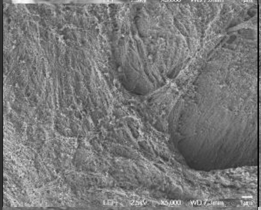 | 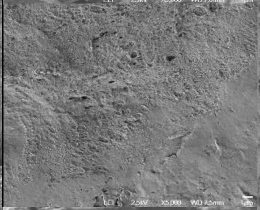 | 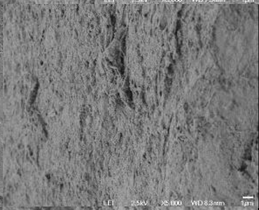 |
| | Sample 2 | 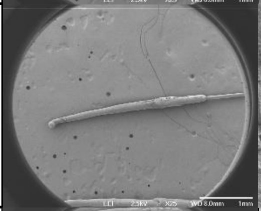 | 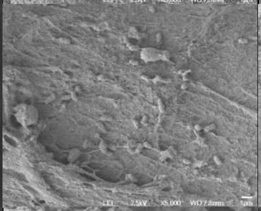 | 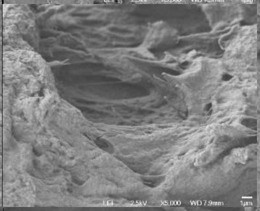 | 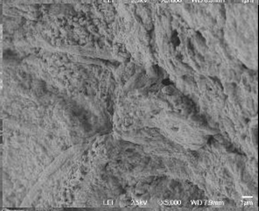 |
| | Sample 3 | 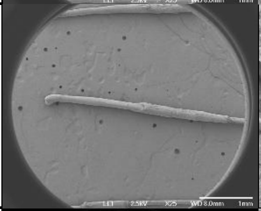 | 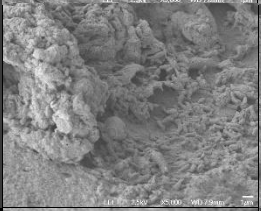 | 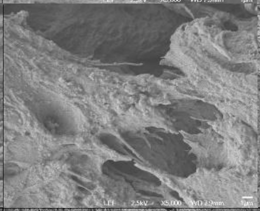 | 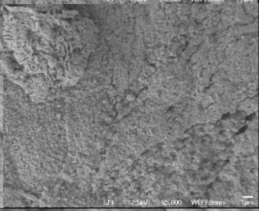 |
| P032<br>Dandruff<br>Dandruff Score: 2.825 | Sample 1 | 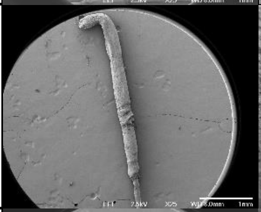 | 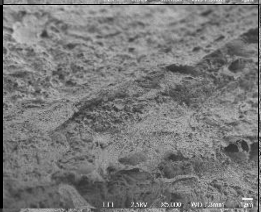 | 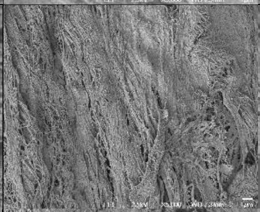 | 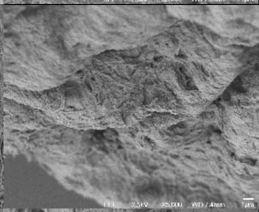 |
| | Sample 2 | 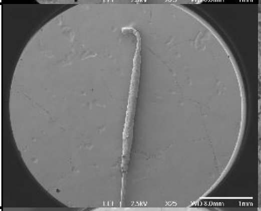 | 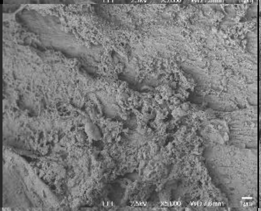 | 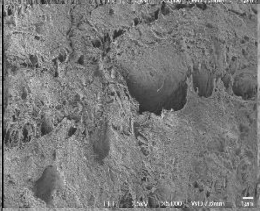 | 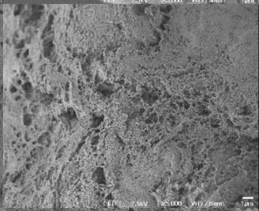 |
| | Sample 3 | 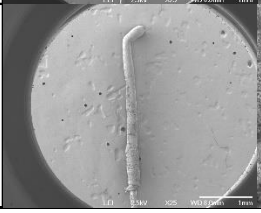 | 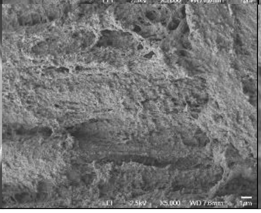 | 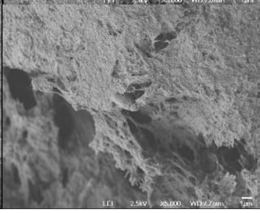 | 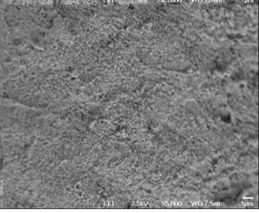 |

| Sample ID | | Full | Top | Middle | Root |
|---|---|---|---|---|---|
| P033<br>Dandruff<br>Dandruff Score: 2.125 | Sample 1 | 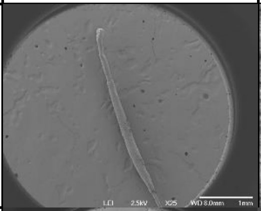 | 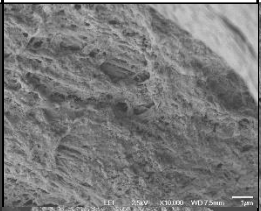 | 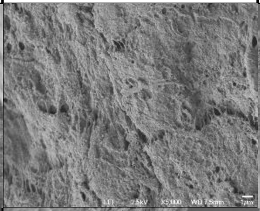 | 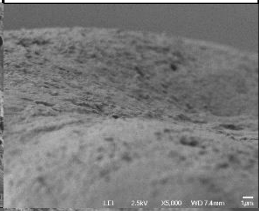 |
| | Sample 2 | 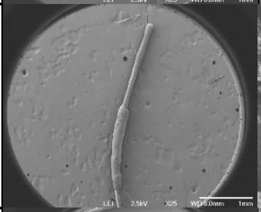 | 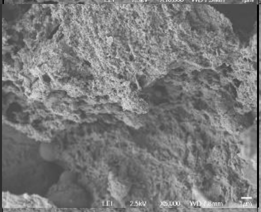 | 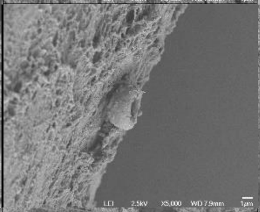 | 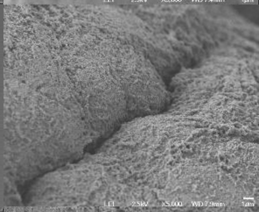 |
| | Sample 3 | 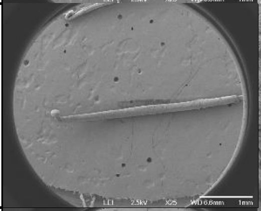 | 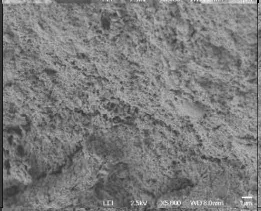 | 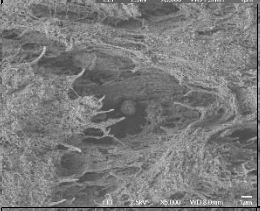 | 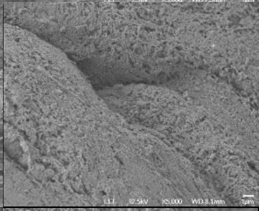 |
| P034<br>Healthy<br>Dandruff Score: 0 | Sample 1 | 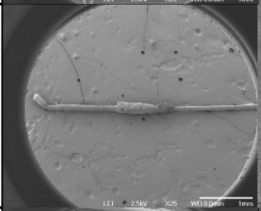 | 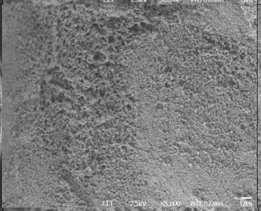 | 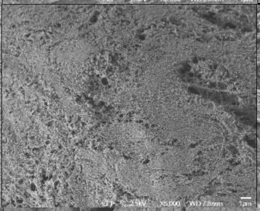 | 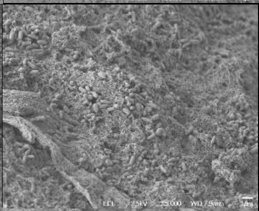 |
| | Sample 2 | 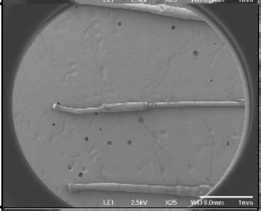 | 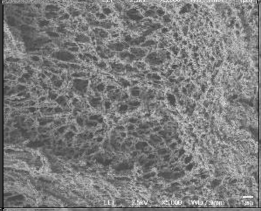 | 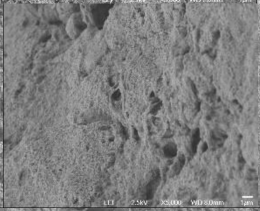 | 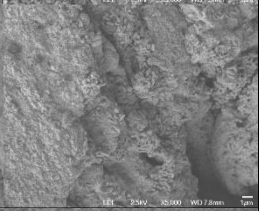 |
| | Sample 3 | 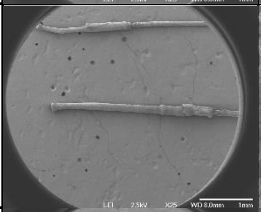 | 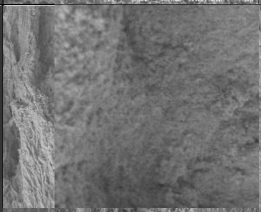 | 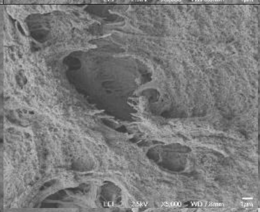 | 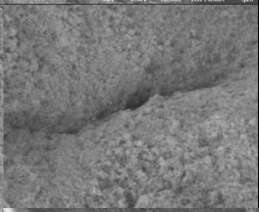 |
| P036<br>Dandruff<br>Dandruff Score: 3 | Sample 1 | 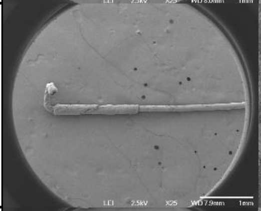 | 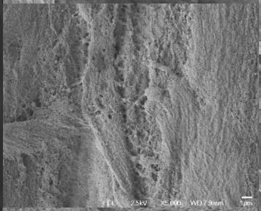 | 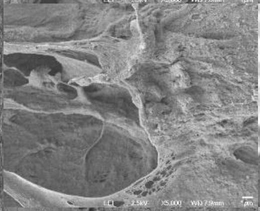 | 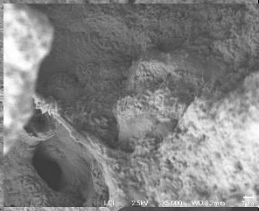 |
| | Sample 2 | 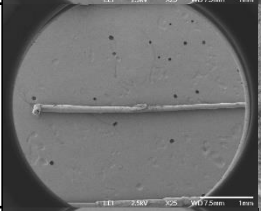 | 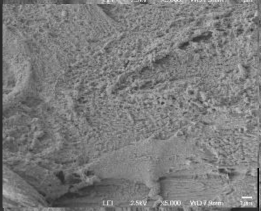 | 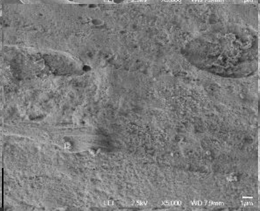 | 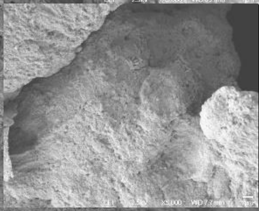 |
| | Sample 3 | 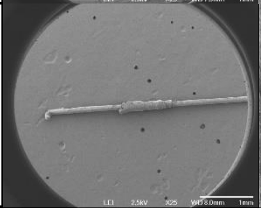 | 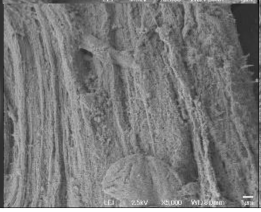 | 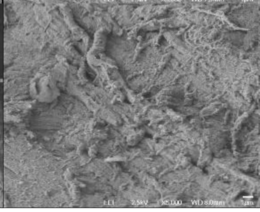 | 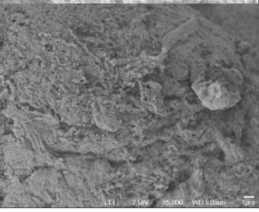 |

| Sample ID | | Full | Top | Middle | Root |
|---|---|---|---|---|---|
| P037<br>Dandruff<br>Dandruff Score: 2.875 | Sample 1 | 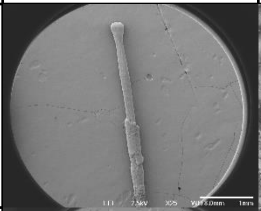 | 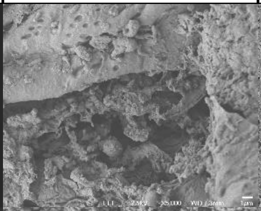 | 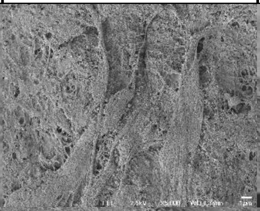 | 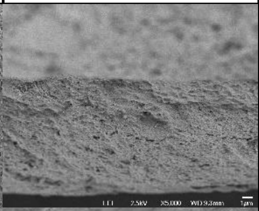 |
| | Sample 2 | 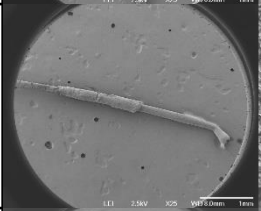 | 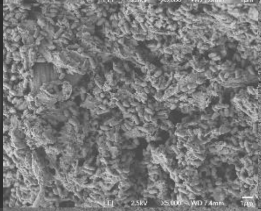 | 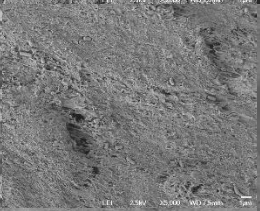 | 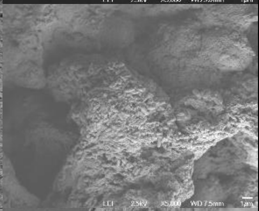 |
| | Sample 3 | 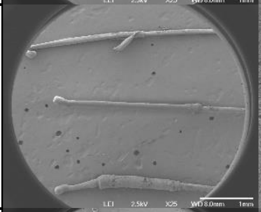 | 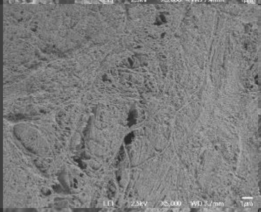 | 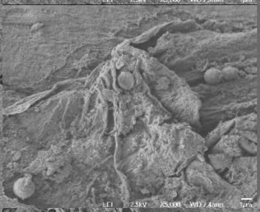 | 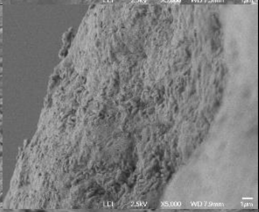 |
| P039<br>Dandruff<br>Dandruff Score: 2.75 | Sample 1 | 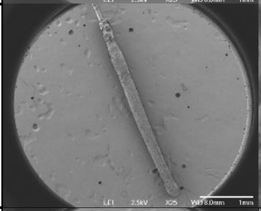 | 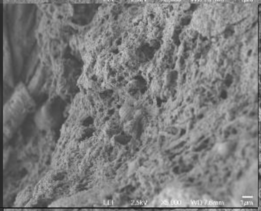 | 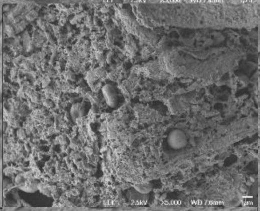 | 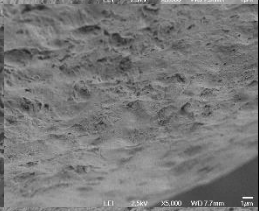 |
| | Sample 2 | 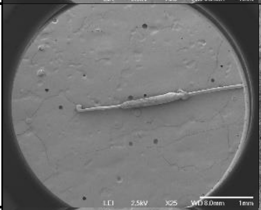 | 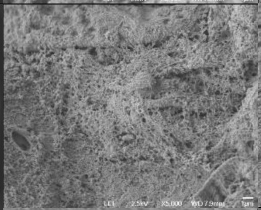 | 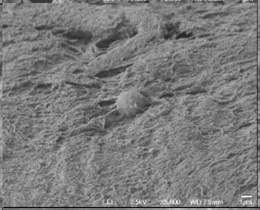 | 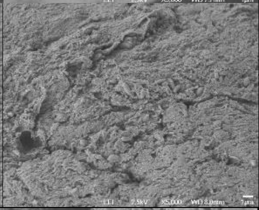 |
| | Sample 3 | 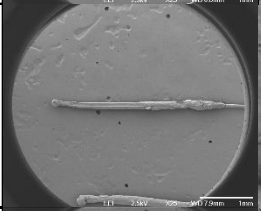 | 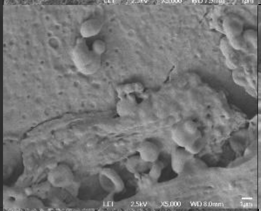 | 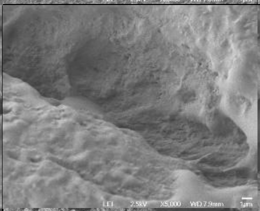 | 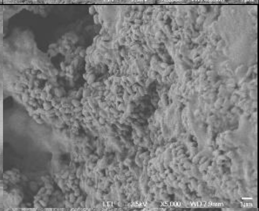 |
| P040<br>Dandruff<br>Dandruff Score: 2.75 | Sample 1 Dandruff | 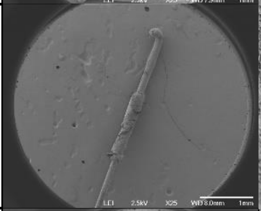 | 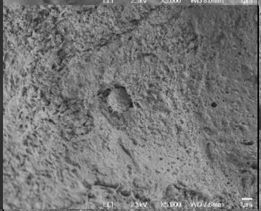 | 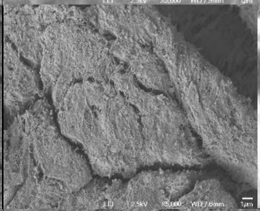 | 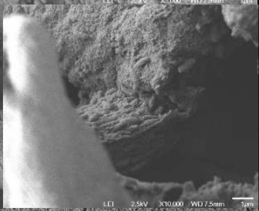 |
| | Sample 2 | 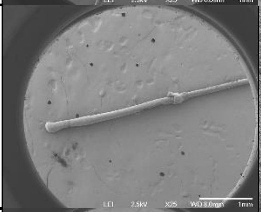 | 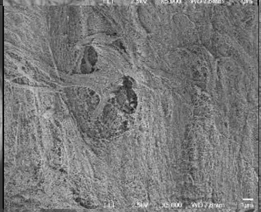 | 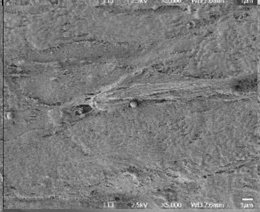 | 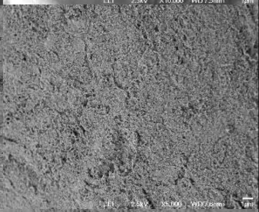 |
| | Sample 3 | 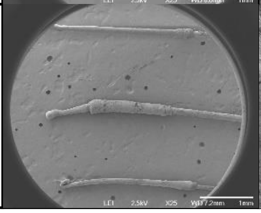 | 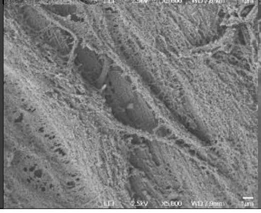 | 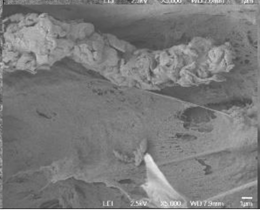 | 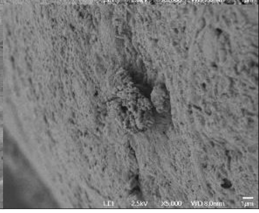 |

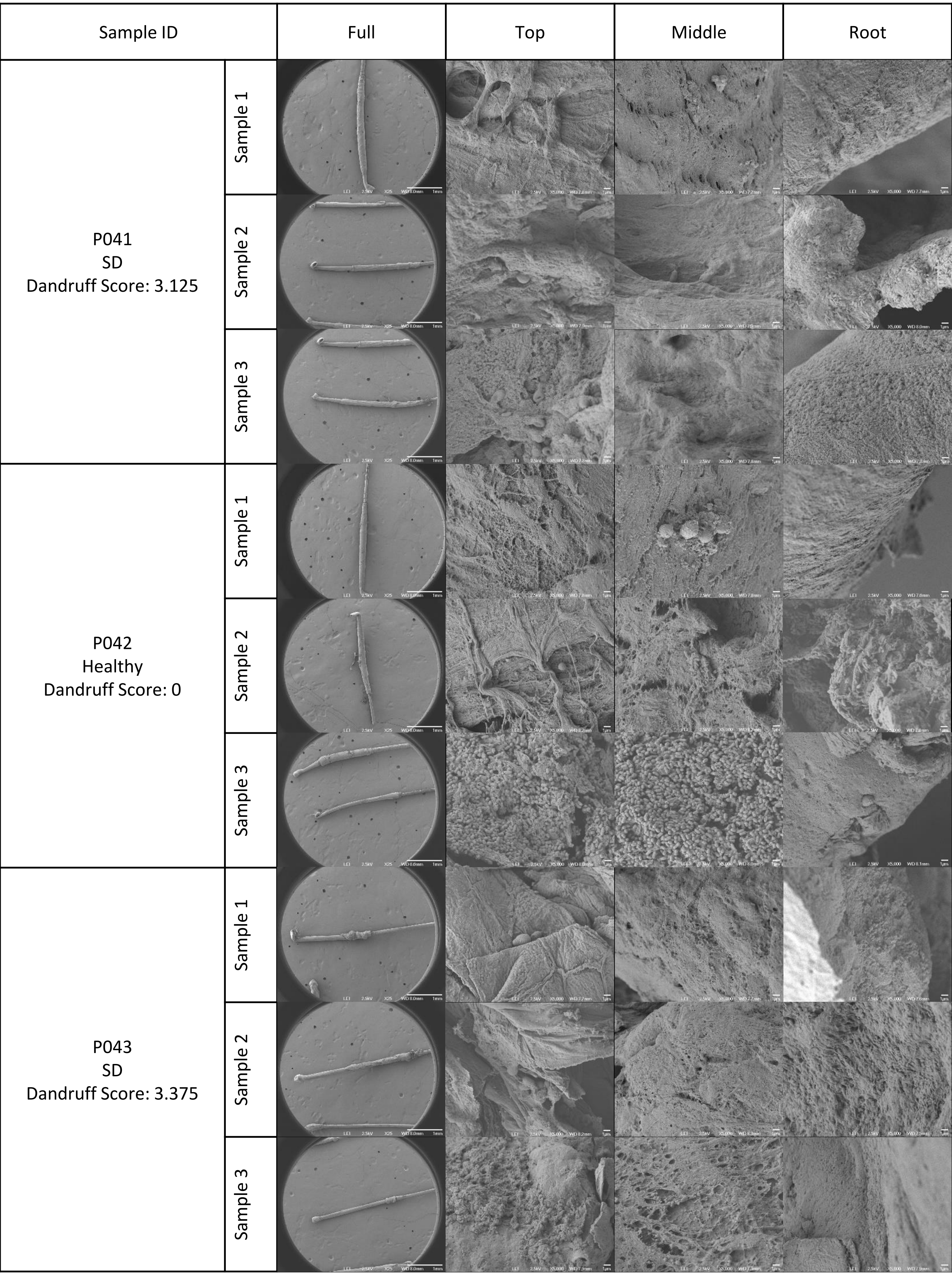

Sample ID
Full
Top
Middle
Root
P041
SD
Dandruff Score: 3.125
Sample 1
Sample 2
Sample 3
P042
Healthy
Dandruff Score: 0
Sample 1
Sample 2
Sample 3
P043
SD
Dandruff Score: 3.375
Sample 1
Sample 2
Sample 3

| Sample ID | | Full | Top | Middle | Root |
|---|---|---|---|---|---|
| P045<br>Dandruff<br>Dandruff Score: 2.75 | Sample 1 | 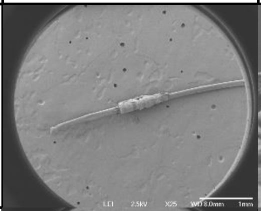 | 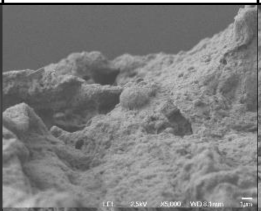 | 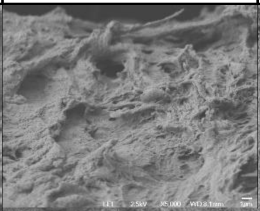 | 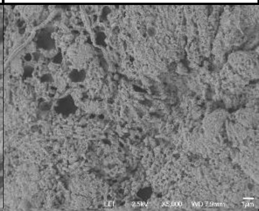 |
| | Sample 2 | 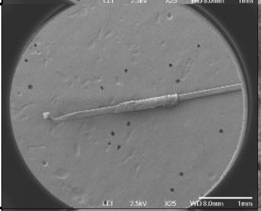 | 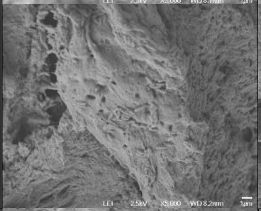 | 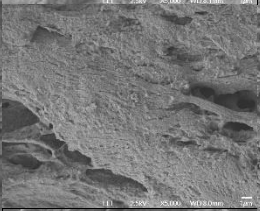 | 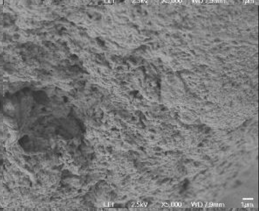 |
| | Sample 3 | 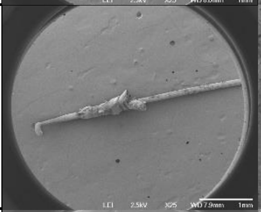 | 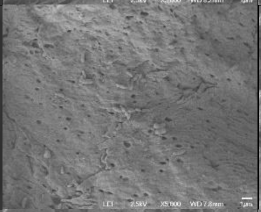 | 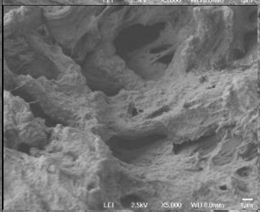 | 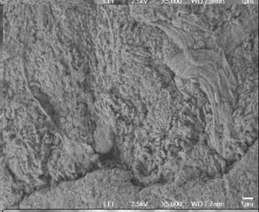 |
| P047<br>Healthy<br>Dandruff Score: 0 | Sample 1 | 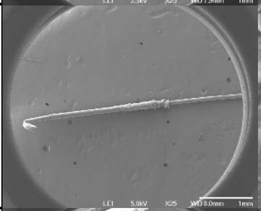 | 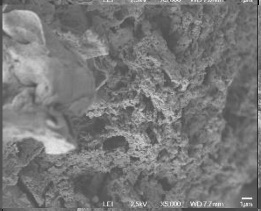 | 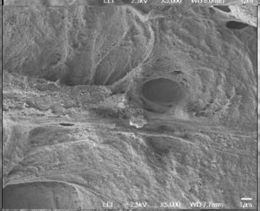 | 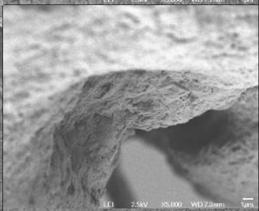 |
| | Sample 2 | 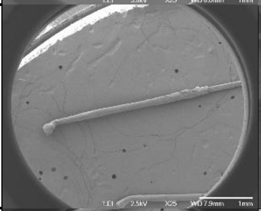 | 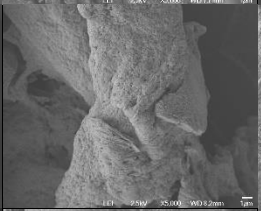 | 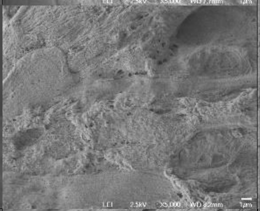 | 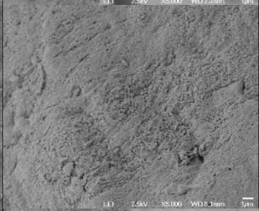 |
| | Sample 3 | 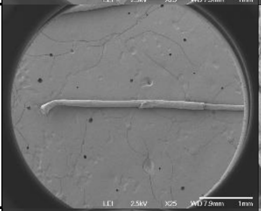 | 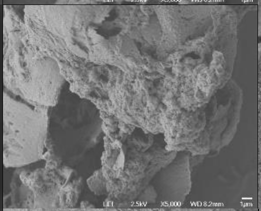 | 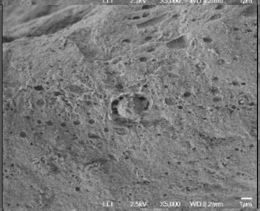 | 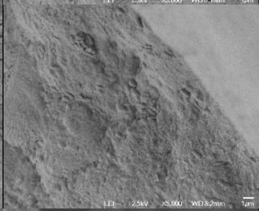 |
| P048<br>SD<br>Dandruff Score: 4 | Sample 1 | 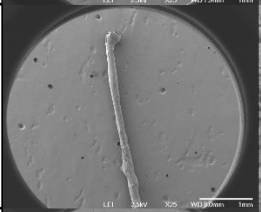 | 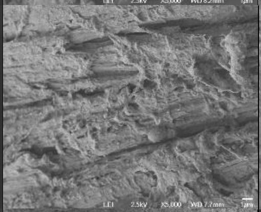 | 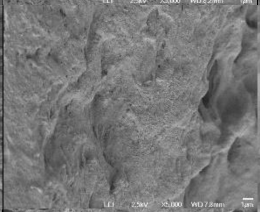 | 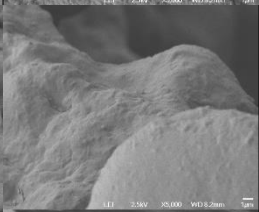 |
| | Sample 2 | 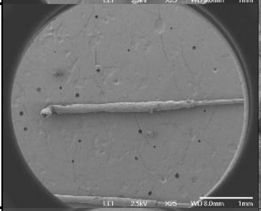 | 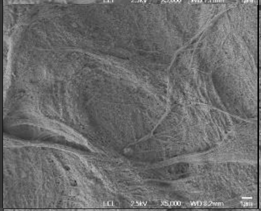 | 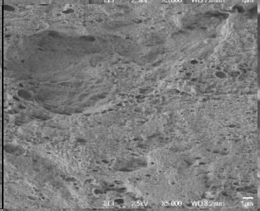 | 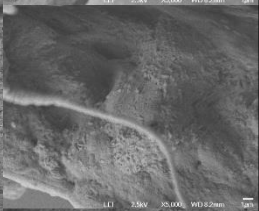 |
| | Sample 3 | 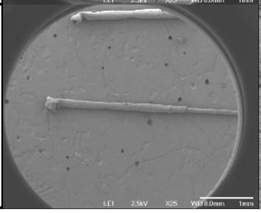 | 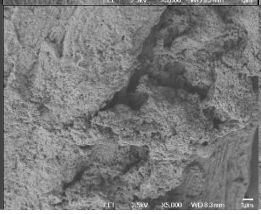 | 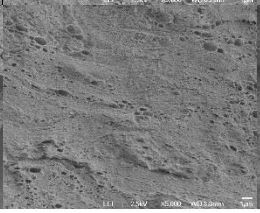 | 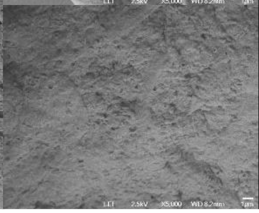 |

| Sample ID | | Full | Top | Middle | Root |
|---|---|---|---|---|---|
| P051<br>SD<br>Dandruff Score: 4.625 | Sample 1 | 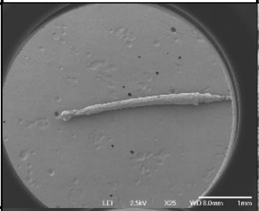 | 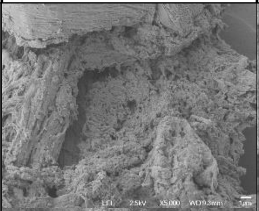 | 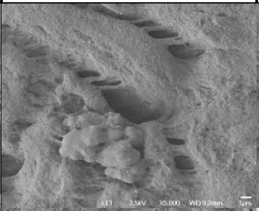 | 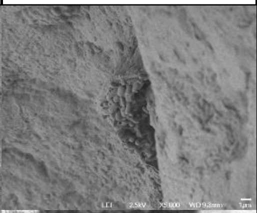 |
| | Sample 2 | 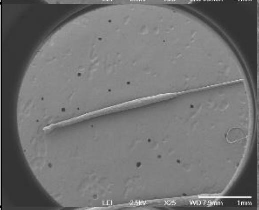 | 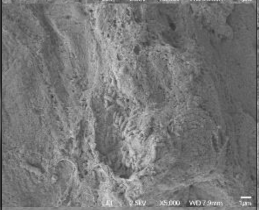 | 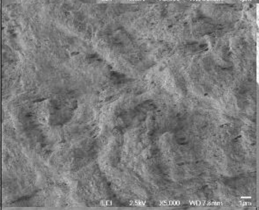 | 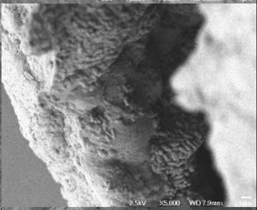 |
| | Sample 3 | 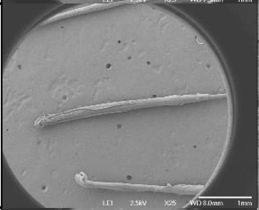 | 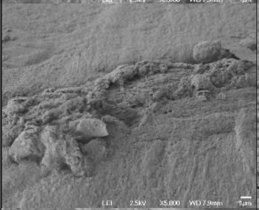 | 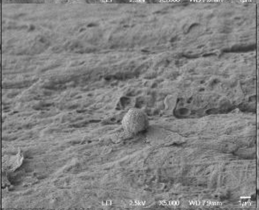 | 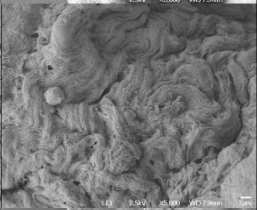 |
| P052<br>Dandruff<br>Dandruff Score: 3 | Sample 1 | 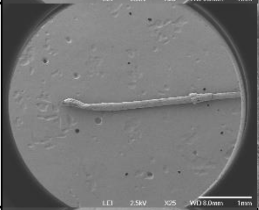 | 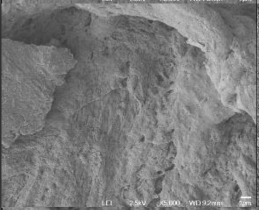 | 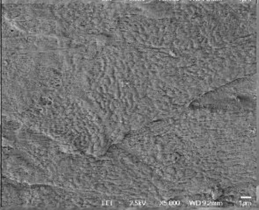 | 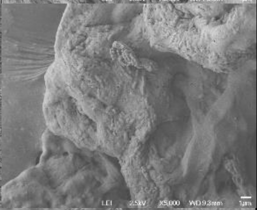 |
| | Sample 2 | 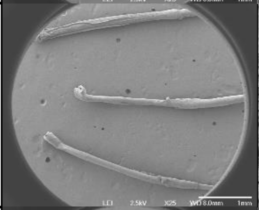 | 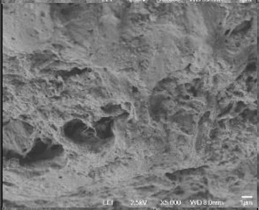 | 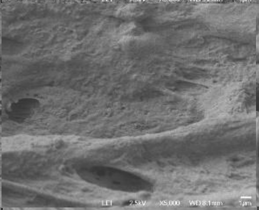 | 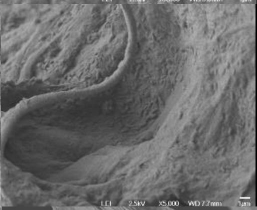 |
| | Sample 3 | 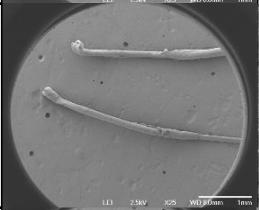 | 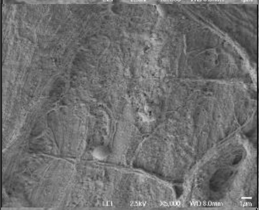 | 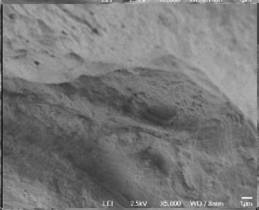 | 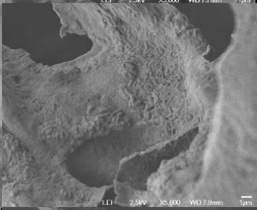 |
| P053<br>SD<br>Dandruff Score: 3.25 | Sample 1 | 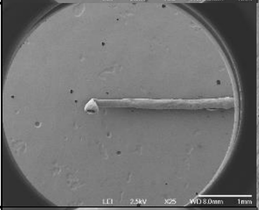 | 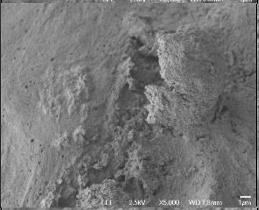 | 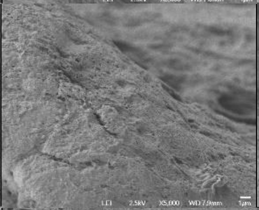 | 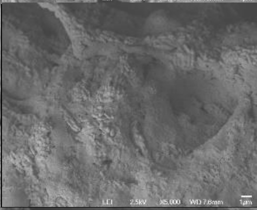 |
| | Sample 2 | 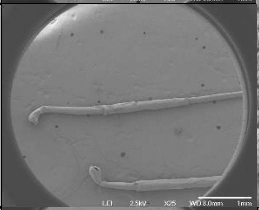 | 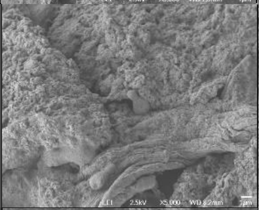 | 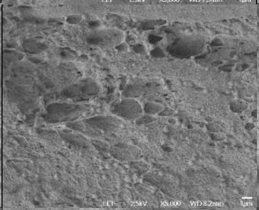 | 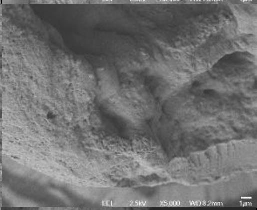 |
| | Sample 3 | 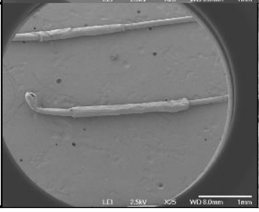 | 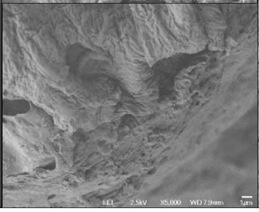 | 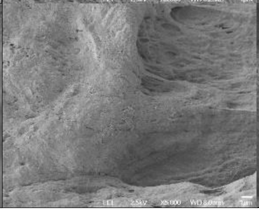 | 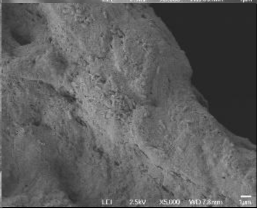 |

| Sample ID | | Full | Top | Middle | Root |
|---|---|---|---|---|---|
| P055<br>Dandruff<br>Dandruff Score: 2.625 | Sample 1 | 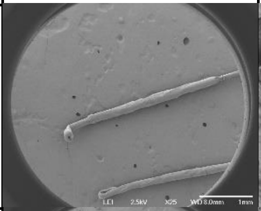 | 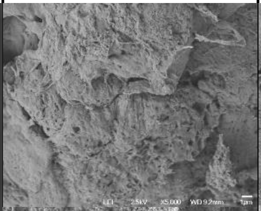 | 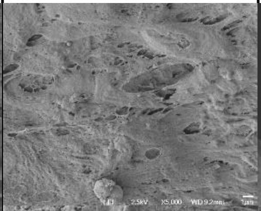 | 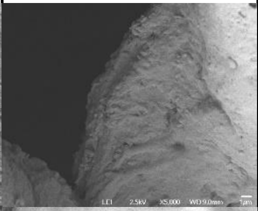 |
| | Sample 2 | 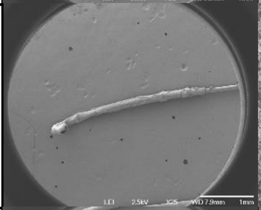 | 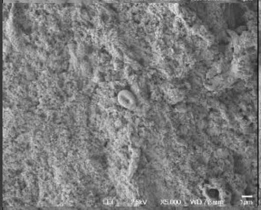 | 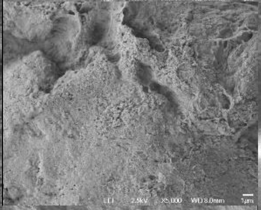 | 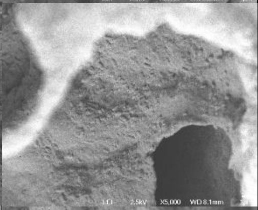 |
| | Sample 3 | 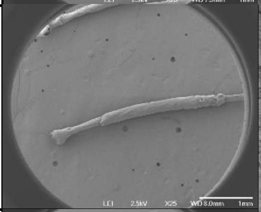 | 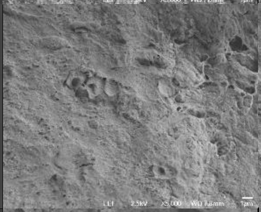 | 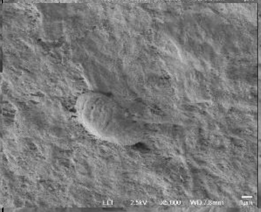 | 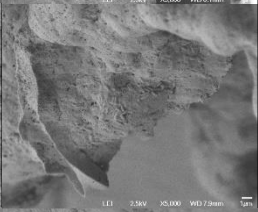 |
| P056<br>SD<br>Dandruff Score: 3.625 | Sample 1 | 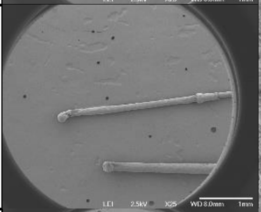 | 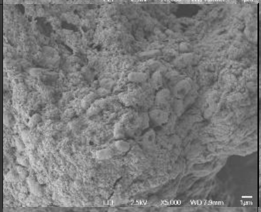 | 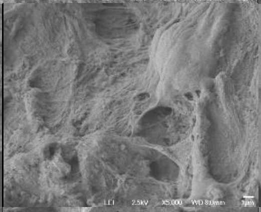 | 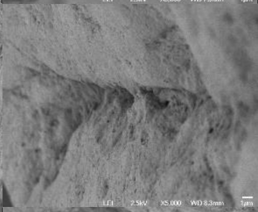 |
| | Sample 2 | 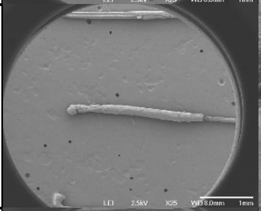 | 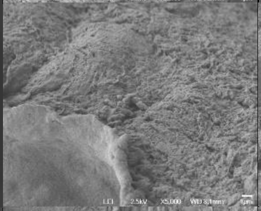 | 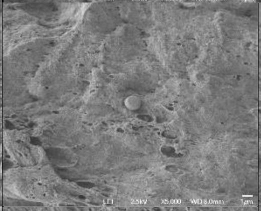 | 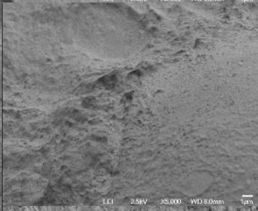 |
| | Sample 3 | 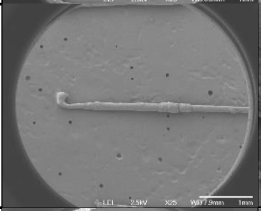 | 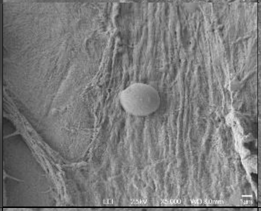 | 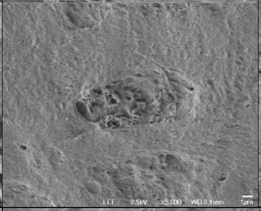 | 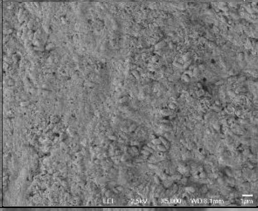 |
| P059<br>Dandruff<br>Dandruff Score: 2.5 | Sample 1 | 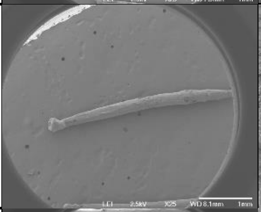 | 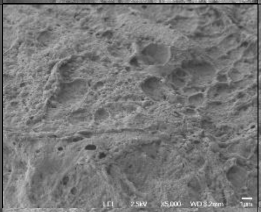 | 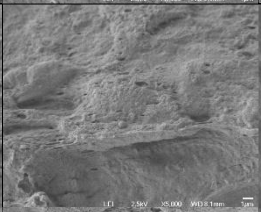 | 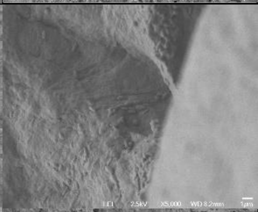 |
| | Sample 2 | 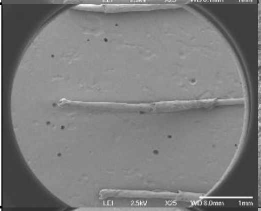 | 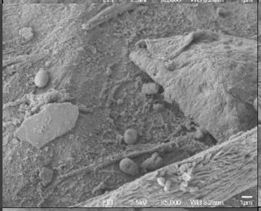 | 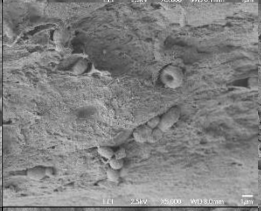 | 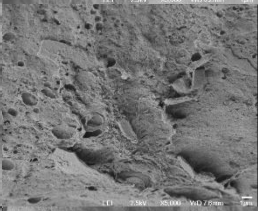 |
| | Sample 3 | 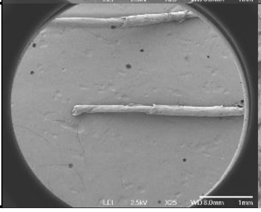 | 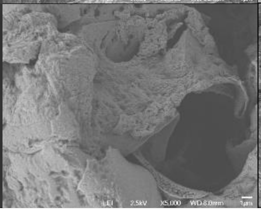 | 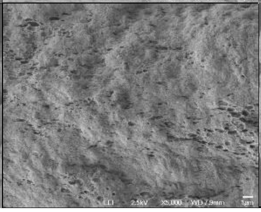 | 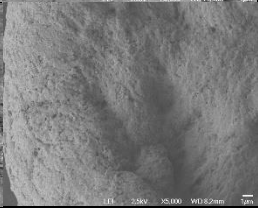 |

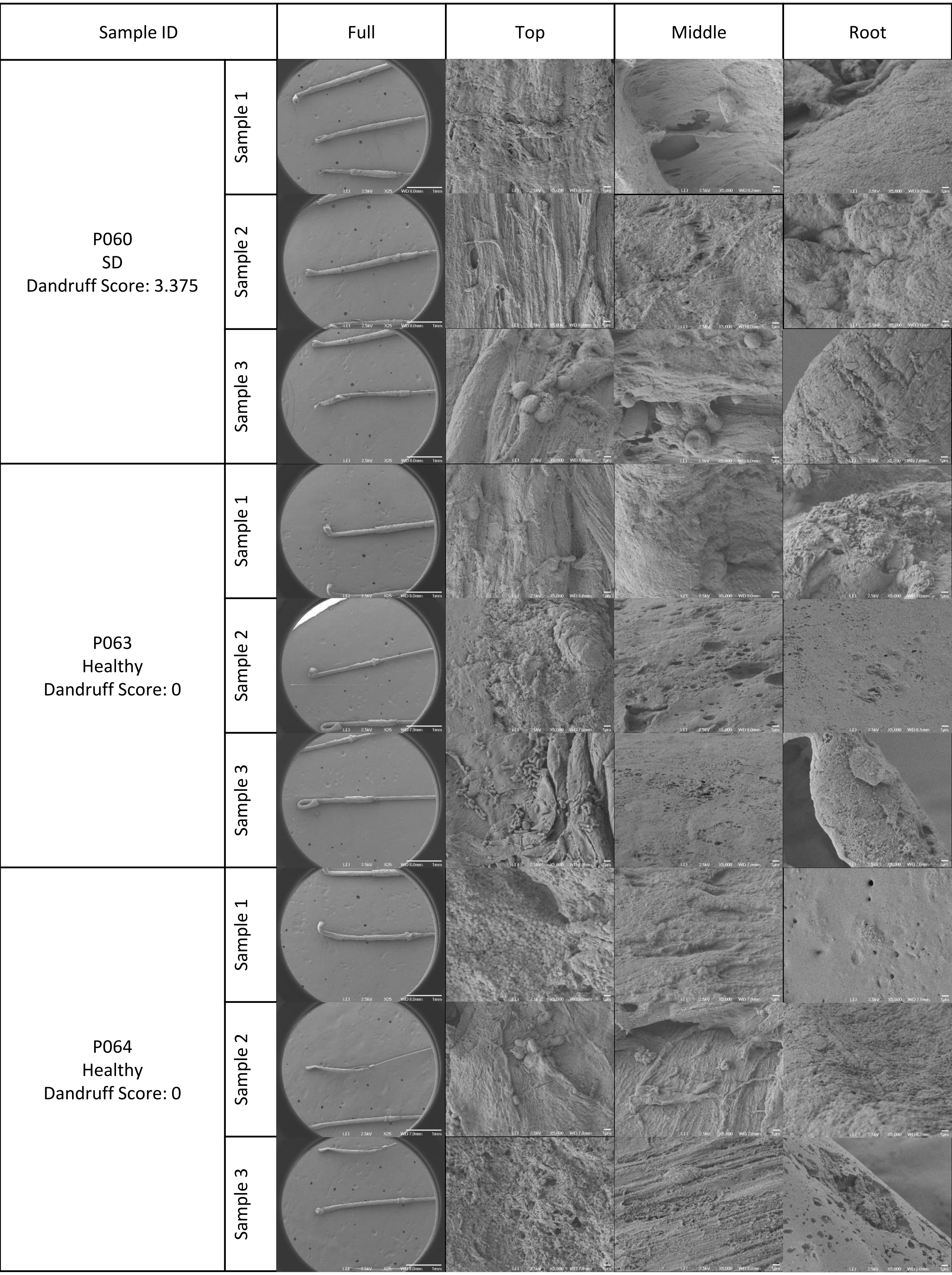
Sample ID
Full
Top
Middle
Root
P060
SD
Dandruff Score: 3.375
Sample 1
Sample 2
Sample 3
P063
Healthy
Dandruff Score: 0
Sample 1
Sample 2
Sample 3
P064
Healthy
Dandruff Score: 0
Sample 1
Sample 2
Sample 3

| Sample ID | | Full | Top | Middle | Root |
|---|---|---|---|---|---|
| P066<br>Healthy<br>Dandruff Score: 0 | Sample 1 | 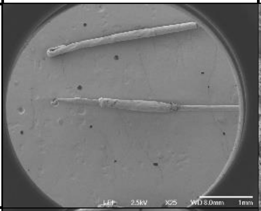 | 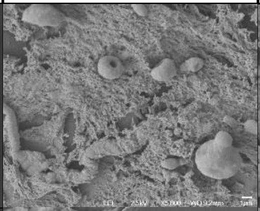 | 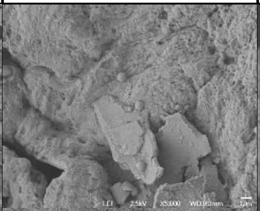 | 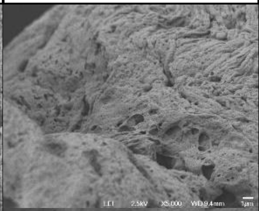 |
| | Sample 2 | 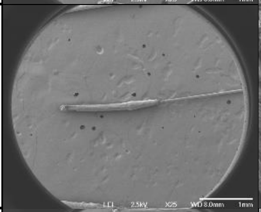 | 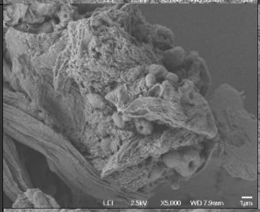 | 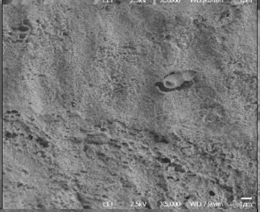 | 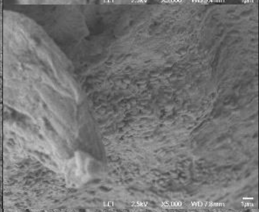 |
| | Sample 3 | 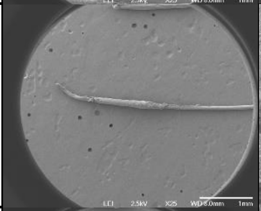 | 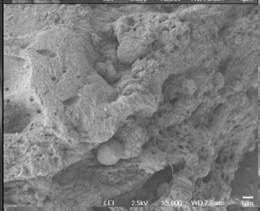 | 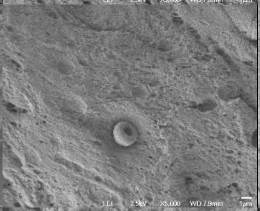 | 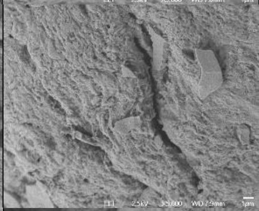 |
| P067<br>Healthy<br>Dandruff Score: 0 | Sample 1 | 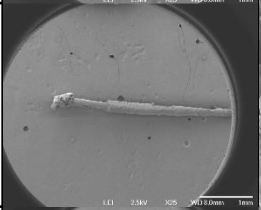 | 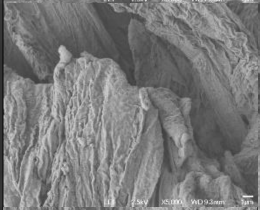 | 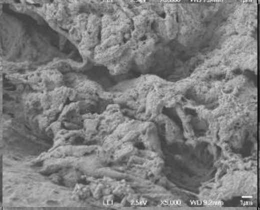 | 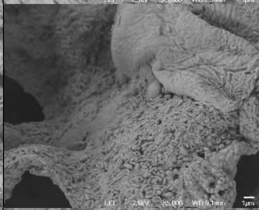 |
| | Sample 2 | 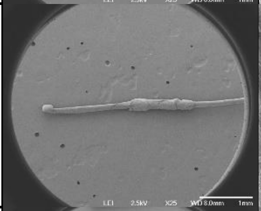 | 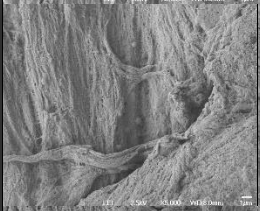 | 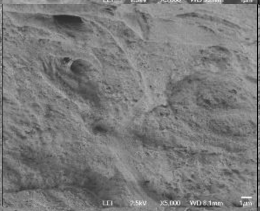 | 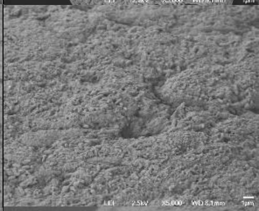 |
| | Sample 3 | 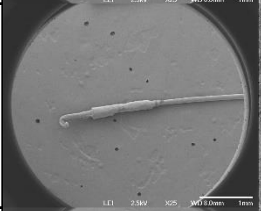 | 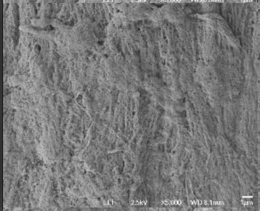 | 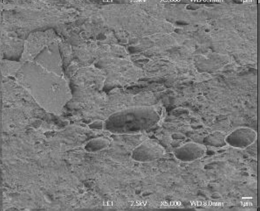 | 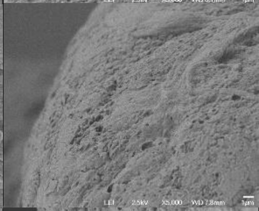 |
| P068<br>Healthy<br>Dandruff Score: 0 | Sample 1 | 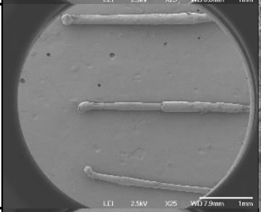 | 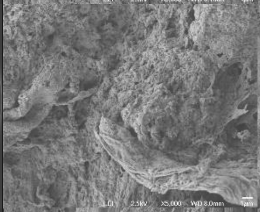 | 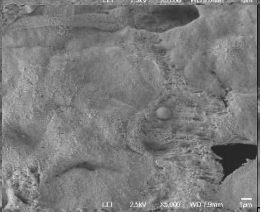 | 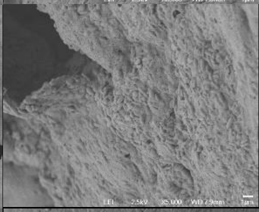 |
| | Sample 2 | 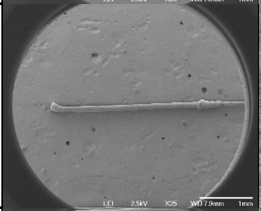 | 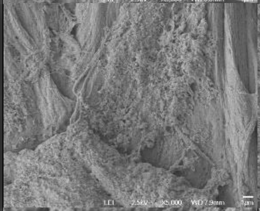 | 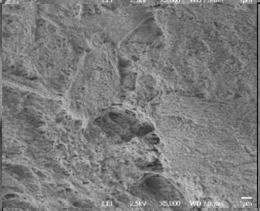 | 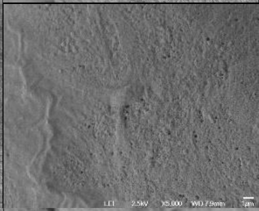 |
| | Sample 3 | 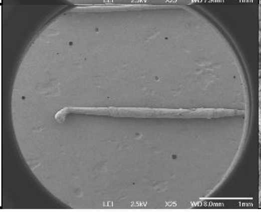 | 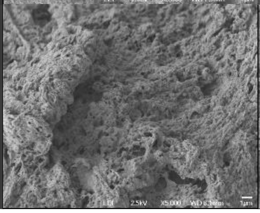 | 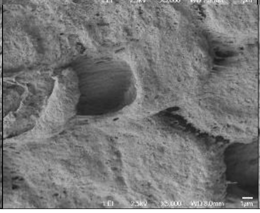 | 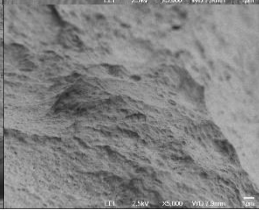 |

| Sample ID | | Full | Top | Middle | Root |
|---|---|---|---|---|---|
| P069 Healthy Dandruff Score: 0 | Sample 1 | 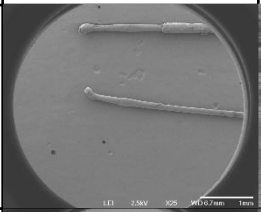 | 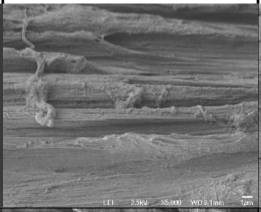 | 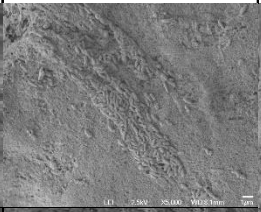 | 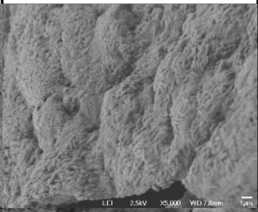 |
| | Sample 2 | 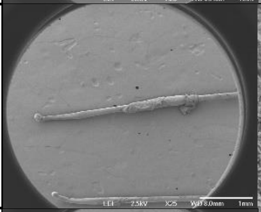 | 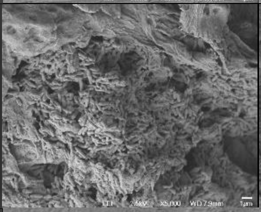 | 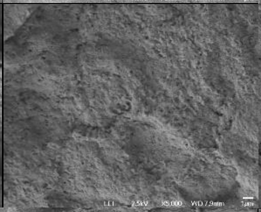 | 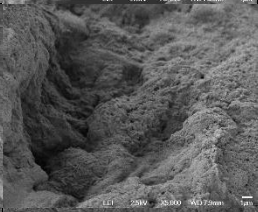 |
| | Sample 3 | 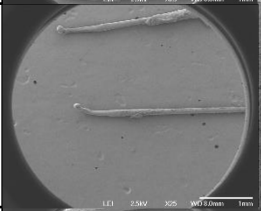 | 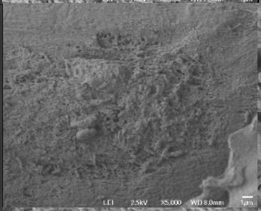 | 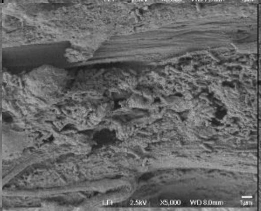 | 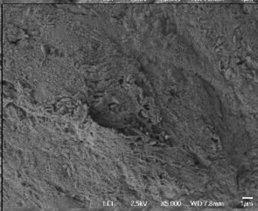 |
| P070 Healthy Dandruff Score: 0 | Sample 1 | 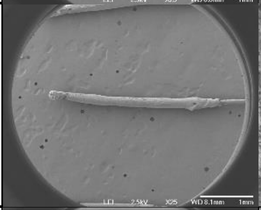 | 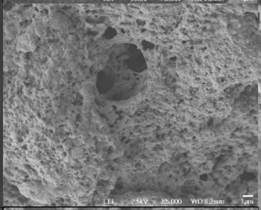 | 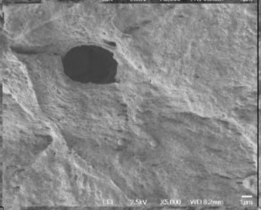 | 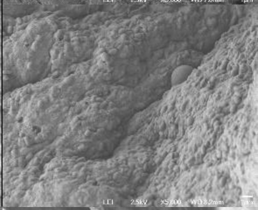 |
| | Sample 2 | 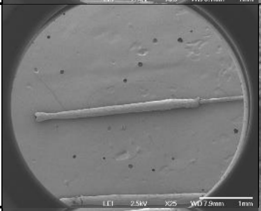 | 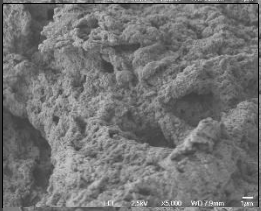 | 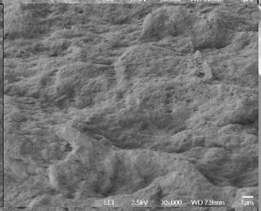 | 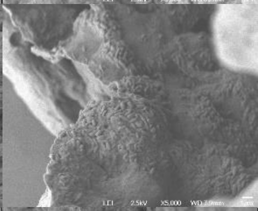 |
| | Sample 3 | 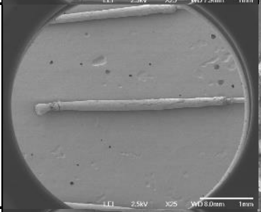 | 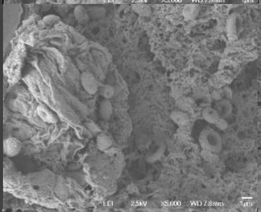 | 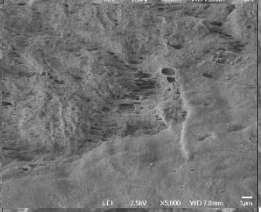 | 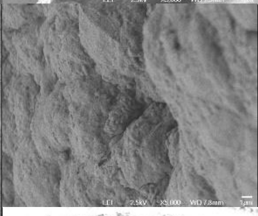 |
| P071 Healthy Dandruff Score: 0 | Sample 1 | 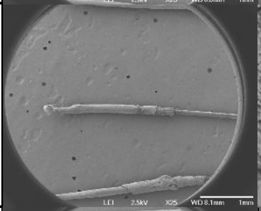 | 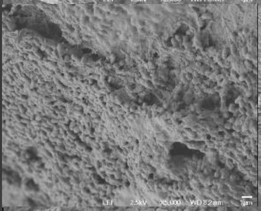 | 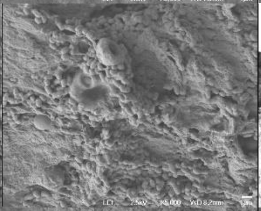 | 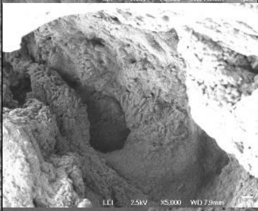 |
| | Sample 2 | 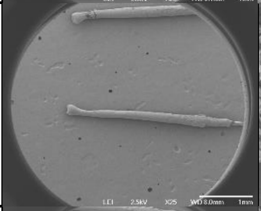 | 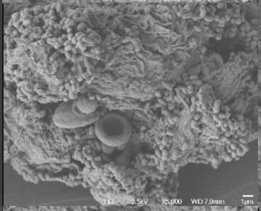 | 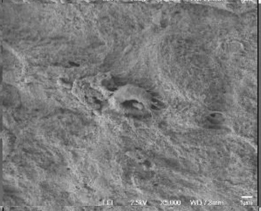 | 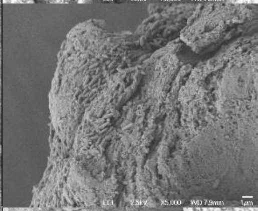 |
| | Sample 3 | 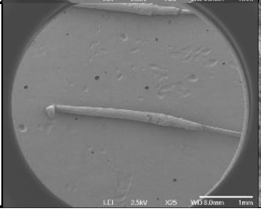 | 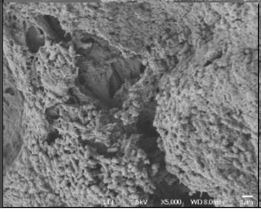 | 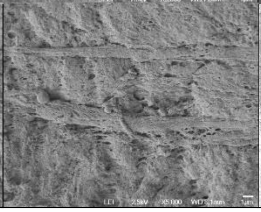 | 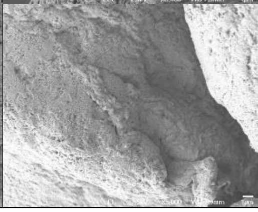 |

| Sample ID | | Full | Top | Middle | Root |
|---|---|---|---|---|---|
| P072<br>Healthy<br>Dandruff Score: 0 | Sample 1 | 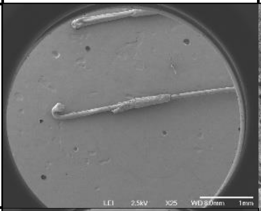 | 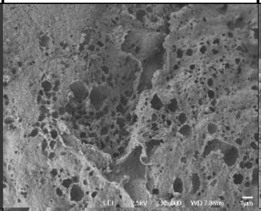 | 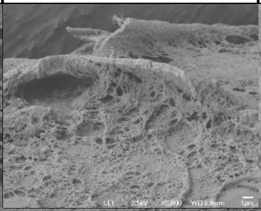 | 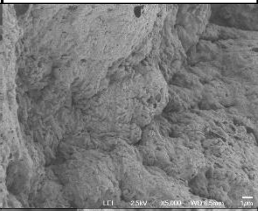 |
| | Sample 2 | 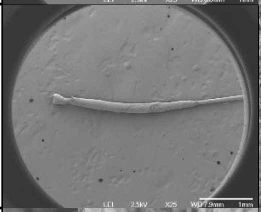 | 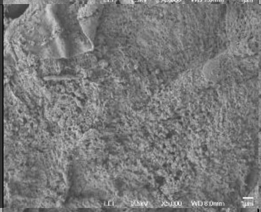 | 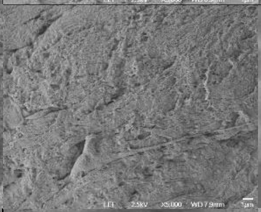 | 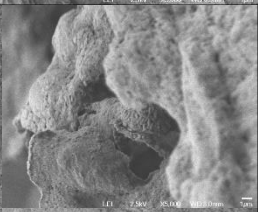 |
| | Sample 3 | 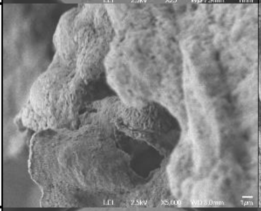 | 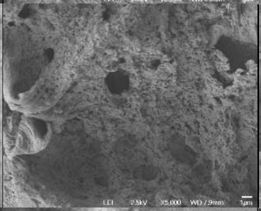 | 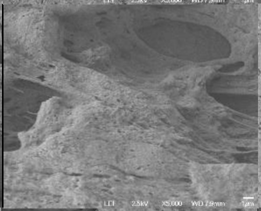 | 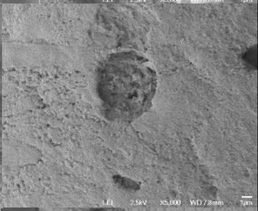 |
| P074<br>Healthy<br>Dandruff Score: 0 | Sample 1 | 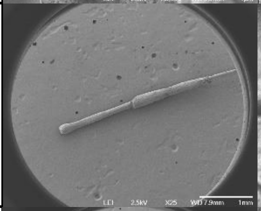 | 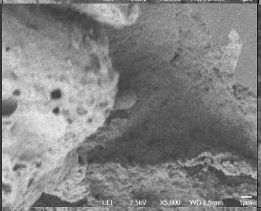 | 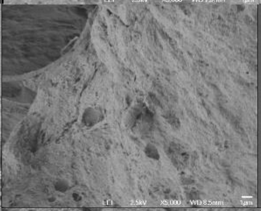 | 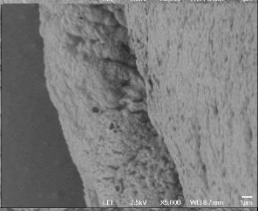 |
| | Sample 2 | 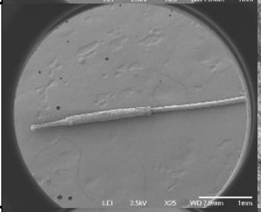 | 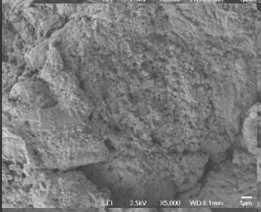 | 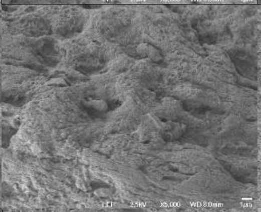 | 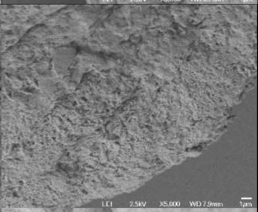 |
| | Sample 3 | 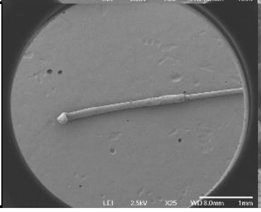 | 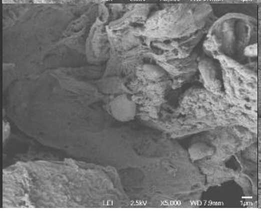 | 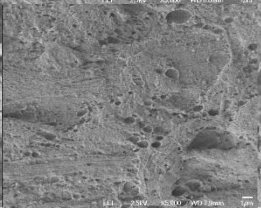 | 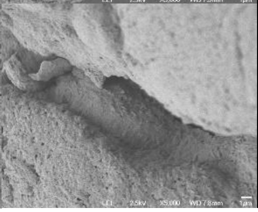 |
