## Supplementary table 1 for "Decoding scalp health and microbiome dysbiosis in dandruff"

**Table S1. Semi-quantitative estimation of microbial structures.** Semi quantitative estimation of the relative abundance of rod, cocci, and fungus structures from the images based on a 5 point (0 to5) scoring system visualized by SEM imaging of hair follicles (Fig. S4) collected from healthy, dandruff and seborrheic dermatitis individuals.

| Sample ID | Replicate | Condition | | | | | | | | |
|---|---|---|---|---|---|---|---|---|---|---|
| | | | Infundibulum | | | Isthmas | | | Bulb | | |
| | | | Bacteria | | Fungi | Bacteria | | Fungi | Bacteria | | Fungi |
| | | | Cocci | Rods | | Cocci | Rods | | Cocci | Rods | |
| P008 | 1 | Healthy | + | ++ | + | ++ | - | - | + | +++ | - |
| | 2 | Healthy | + | ++++ | +++ | + | +++ | - | ++ | ++ | - |
| | 3 | Healthy | + | ++ | ++ | + | + | - | - | - | - |
| P009 | 1 | Healthy | + | - | - | ++ | +++ | - | - | - | - |
| | 2 | Healthy | - | ++++ | - | - | ++++ | - | - | ++++ | - |
| | 3 | Healthy | - | ++++ | - | + | - | - | + | - | - |
| P010 | 1 | Healthy | - | ++++ | - | - | - | - | - | ++ | - |
| | 2 | Healthy | ++ | ++ | ++ | + | + | - | + | - | - |
| | 3 | Healthy | + | - | - | - | +++ | - | - | ++ | - |
| P012 | 1 | SD | ++ | - | - | - | - | - | - | +++ | - |
| | 2 | SD | - | - | - | - | - | - | - | ++++ | - |
| | 3 | SD | - | - | - | ++ | - | + | - | - | - |
| P013 | 1 | SD | ++ | - | +++ | +++ | - | - | ++ | ++ | - |
| | 2 | SD | + | - | + | + | - | - | - | - | - |
| | 3 | SD | - | - | ++ | - | - | - | - | - | - |
| P014 | 1 | SD | ++ | +++ | + | ++ | - | - | ++ | + | - |
| | 2 | SD | ++ | - | + | - | - | - | + | ++ | - |
| | 3 | SD | +++ | - | - | +++ | - | - | - | - | - |
| P016 | 1 | Healthy | +++ | ++ | + | +++ | - | - | +++ | ++++ | - |
| | 2 | Healthy | - | + | - | + | - | - | - | - | - |
| | 3 | Healthy | +++ | ++ | + | ++ | - | - | + | +++ | - |
| P017 | 1 | SD | - | ++++ | ++ | + | - | - | - | - | - |
| | 2 | SD | ++ | - | + | ++ | - | + | ++ | - | - |
| | 3 | SD | +++ | - | + | + | - | - | + | ++++ | - |
| P018 | 1 | SD | + | ++ | +++ | + | - | - | + | - | - |
| | 2 | SD | - | - | + | - | - | + | - | - | - |
| | 3 | SD | ++ | +++ | + | - | - | + | - | ++++ | + |
| P019 | 1 | Healthy | - | ++ | + | + | - | + | + | ++++ | - |
| | 2 | Healthy | - | ++ | + | + | - | + | + | - | - |
| | 3 | Healthy | + | + | - | - | - | - | + | +++ | - |
| P021 | 1 | Healthy | + | - | ++ | + | - | - | ++ | +++ | - |
| | 2 | Healthy | ++ | + | ++ | - | - | - | ++ | ++++ | - |
| | 3 | Healthy | +++ | - | - | ++ | + | - | ++ | + | - |

| | | | | | | | | | | | |
|---|---|---|---|---|---|---|---|---|---|---|---|
| P022 | 1 | Healthy | + | + | - | - | - | - | ++ | - | - |
| | 2 | Healthy | - | ++ | +++ | - | - | - | - | ++ | - |
| | 3 | Healthy | +++ | - | - | - | - | + | - | - | + |
| P023 | 1 | Healthy | ++ | - | + | - | - | - | +++ | ++ | + |
| | 2 | Healthy | ++ | - | + | - | - | - | - | ++ | - |
| | 3 | Healthy | - | - | - | - | - | - | + | +++ | - |
| P024 | 1 | Healthy | +++ | + | + | + | - | - | maybe+ | - | - |
| | 2 | Healthy | ++ | + | ++ | + | - | - | - | - | - |
| | 3 | Healthy | + | ++ | - | + | - | - | - | ++++ | - |
| P025 | 1 | SD | - | - | - | - | - | - | - | - | - |
| | 2 | SD | - | + | - | - | - | - | ++ | ++++ | + |
| | 3 | SD | ++ | - | - | - | - | - | + | - | - |
| P026 | 1 | Healthy | +++ | +++ | + | +++ | +++ | - | - | +++ | - |
| | 2 | Healthy | ++ | - | + | + | + | + | - | ++++ | - |
| | 3 | Healthy | - | + | +++ | ++ | ++++ | - | ++ | +++ | - |
| P027 | 1 | Healthy | ++ | - | - | - | + | - | - | - | - |
| | 2 | | | | | | | | | | |
| | 3 | | | | | | | | | | |
| P028 | 1 | Dandruff | - | ++ | + | - | - | - | ++ | ++ | - |
| | 2 | Dandruff | - | - | + | + | - | - | ++ | +++ | + |
| | 3 | Dandruff | + | - | + | - | - | - | ++ | - | - |
| P029 | 1 | Healthy | + | + | - | - | ++ | ++ | - | + | + |
| | 2 | Healthy | ++ | - | ++ | ++ | - | - | + | ++ | - |
| | 3 | Healthy | ++ | - | + | - | - | - | - | - | - |
| P030 | 1 | SD | +++ | - | - | ++ | - | - | - | - | - |
| | 2 | SD | - | - | ++ | ++ | - | - | - | ++ | - |
| | 3 | SD | - | - | - | - | - | - | ++ | ++++ | - |
| P031 | 1 | Healthy | + | - | - | ++ | ++ | - | - | - | - |
| | 2 | Healthy | - | +++ | + | - | - | - | - | ++++ | - |
| | 3 | Healthy | + | ++++ | + | - | - | - | - | - | - |
| P032 | 1 | Dandruff | +++ | + | + | + | - | - | + | - | - |
| | 2 | Dandruff | - | - | + | - | + | - | - | - | - |
| | 3 | Dandruff | ++ | + | + | + | - | + | + | - | - |
| P033 | 1 | Dandruff | - | - | - | - | - | - | - | - | - |
| | 2 | Dandruff | ++ | +++ | ++ | - | - | + | - | - | - |
| | 3 | Dandruff | - | ++ | - | - | - | - | + | +++ | - |
| P034 | 1 | Healthy | - | - | - | ++ | - | - | - | ++++ | - |

| | | | | | | | | | | |
|---|---|---|---|---|---|---|---|---|---|---|
| | 2 | Healthy | - | + | + | - | - | - | - | ++++ | - |
| | 3 | Healthy | ++ | ++ | ++ | - | - | - | - | - | - |
| P035 | 1 | Healthy | + | + | - | - | - | - | - | + | - |
| | 2 | Healthy | - | + | - | - | - | - | - | + | + |
| | 3 | | | | | | | | | | |
| P036 | 1 | Dandruff | ++ | - | - | - | - | - | - | ++++ | - |
| | 2 | Dandruff | + | - | + | - | - | - | ++ | ++++ | - |
| | 3 | Dandruff | ++ | - | - | - | - | - | - | +++ | - |
| P037 | 1 | Dandruff | - | +++ | ++ | +++ | - | - | - | + | - |
| | 2 | Dandruff | - | ++++ | + | + | +++ | - | + | ++++ | - |
| | 3 | Dandruff | - | - | - | - | + | + | - | ++++ | - |
| P039 | 1 | Dandruff | + | +++ | ++ | ++ | ++ | ++ | - | - | - |
| | 2 | Dandruff | + | + | - | + | - | + | + | ++ | - |
| | 3 | Dandruff | - | - | ++ | - | - | - | - | ++++ | - |
| P040 | 1 | Dandruff | - | - | + | - | - | - | - | +++ | - |
| | 2 | Dandruff | - | - | + | + | - | - | - | - | - |
| | 3 | Dandruff | - | - | - | - | - | - | - | +++ | - |
| P041 | 1 | SD | - | - | - | - | - | - | + | - | - |
| | 2 | SD | - | - | + | - | - | + | - | + | - |
| | 3 | SD | + | + | +++ | - | - | - | - | - | - |
| P042 | 1 | Healthy | ++ | ++ | - | - | - | + | - | +++ | - |
| | 2 | Healthy | + | - | + | + | ++ | - | - | ++++ | - |
| | 3 | Healthy | - | ++++ | - | - | ++++ | - | - | - | + |
| P043 | 1 | SD | - | + | + | + | - | - | - | +++ | - |
| | 2 | SD | ++ | - | + | ++ | + | - | ++ | ++++ | - |
| | 3 | SD | - | - | - | ++ | - | - | - | ++ | - |
| P045 | 1 | Dandruff | + | - | + | - | - | - | - | ++++ | - |
| | 2 | Dandruff | - | +++ | - | + | - | - | - | ++++ | - |
| | 3 | Dandruff | - | ++ | - | - | - | - | - | ++++ | - |
| P047 | 1 | Healthy | + | - | - | - | - | - | - | +++ | - |
| | 2 | Healthy | - | - | - | - | - | - | - | ++ | + |
| | 3 | Healthy | + | - | - | - | - | + | + | ++++ | - |
| P048 | 1 | SD | ++ | + | + | - | - | - | - | - | - |
| | 2 | SD | + | - | - | - | - | - | - | ++++ | - |
| | 3 | SD | - | - | - | + | - | - | - | ++++ | - |
| P049 | 1 | SD | - | + | +++ | - | - | ++ | - | ++++ | - |
| P050 | 1 | SD | + | - | - | - | - | - | - | ++++ | - |

| | | | | | | | | | | | |
|---|---|---|---|---|---|---|---|---|---|---|---|
| P051 | 1 | SD | + | - | - | - | - | - | - | +++ | - |
| | 2 | SD | ++ | ++ | - | - | - | - | - | ++++ | - |
| | 3 | SD | - | - | - | - | - | + | - | - | - |
| P052 | 1 | Dandruff | - | - | - | - | - | - | - | +++ | - |
| | 2 | Dandruff | + | - | + | - | - | - | - | ++++ | - |
| | 3 | Dandruff | ++ | - | + | - | - | - | - | ++++ | - |
| P053 | 1 | SD | - | - | - | - | - | - | - | ++++ | - |
| | 2 | SD | + | - | ++ | +++ | - | - | - | ++++ | - |
| | 3 | SD | - | - | - | - | - | - | - | +++ | - |
| P054 | 1 | Healthy | - | - | - | + | - | - | - | - | - |
| P055 | 1 | Dandruff | + | + | - | + | - | - | - | ++ | + |
| | 2 | Dandruff | - | - | + | - | - | - | - | - | ++++ |
| | 3 | Dandruff | - | + | ++ | ++ | - | + | - | +++ | - |
| P056 | 1 | SD | - | - | +++ | - | - | - | - | - | - |
| | 2 | SD | + | - | - | - | - | + | + | - | + |
| | 3 | SD | + | - | + | + | ++ | - | - | ++++ | + |
| P057 | 1 | Dandruff | - | - | - | - | - | - | - | +++ | - |
| P058 | 1 | Dandruff | + | - | + | - | - | - | - | - | - |
| P059 | 1 | Dandruff | +++ | - | - | + | - | + | ++ | +++ | + |
| | 2 | Dandruff | ++ | - | ++ | ++ | - | ++ | - | - | - |
| | 3 | Dandruff | +++ | - | - | - | - | - | ++ | - | - |
| P060 | 1 | SD | - | - | - | - | - | - | +++ | - | + |
| | 2 | SD | + | - | - | - | - | - | + | ++++ | - |
| | 3 | SD | - | - | ++ | - | - | ++ | - | ++++ | - |
| P061 | 1 | SD | ++ | - | - | + | - | - | - | ++++ | - |
| P062 | 1 | Healthy | - | + | + | + | - | - | ++ | ++++ | - |
| P063 | 1 | Healthy | + | + | - | ++ | ++ | + | - | ++++ | - |
| | 2 | Healthy | - | - | + | - | - | - | - | - | - |
| | 3 | Healthy | +++ | ++++ | +++ | - | - | - | + | - | - |
| P064 | 1 | Healthy | - | ++++ | +++ | + | - | - | - | ++ | - |
| | 2 | Healthy | + | - | ++ | - | - | - | ++ | +++ | - |
| | 3 | Healthy | - | +++ | ++ | - | ++ | + | - | + | - |
| P065 | 1 | Healthy | - | ++++ | ++ | - | ++ | ++ | - | - | - |
| | 2 | Healthy | - | ++ | - | - | +++ | - | - | - | - |

| | | | | | | | | | | | |
|---|---|---|---|---|---|---|---|---|---|---|---|
| P066 | 1 | Healthy | - | - | ++ | + | - | + | + | - | - |
| | 2 | Healthy | + | - | ++ | - | - | + | - | ++++ | - |
| | 3 | Healthy | ++ | - | ++ | - | - | + | - | + | - |
| P067 | 1 | Healthy | - | - | - | - | - | - | - | ++++ | + |
| | 2 | Healthy | + | - | - | - | - | - | - | ++++ | - |
| | 3 | Healthy | - | - | - | + | - | - | + | +++ | - |
| P068 | 1 | Healthy | + | - | - | + | - | + | - | +++ | - |
| | 2 | Healthy | - | - | - | - | - | - | - | - | - |
| | 3 | Healthy | + | - | + | - | - | - | - | +++ | - |
| P069 | 1 | Healthy | - | - | - | - | +++ | - | - | ++++ | - |
| | 2 | Healthy | - | ++++ | - | - | - | - | - | ++++ | - |
| | 3 | Healthy | - | ++ | + | - | ++++ | ++ | - | ++++ | - |
| P070 | 1 | Healthy | - | ++ | - | - | - | + | - | +++ | + |
| | 2 | Healthy | - | + | - | - | + | - | - | ++++ | - |
| | 3 | Healthy | - | +++ | +++ | + | - | - | ++ | +++ | - |
| P071 | 1 | Healthy | - | ++++ | ++ | - | ++++ | ++ | - | ++++ | - |
| | 2 | Healthy | - | ++++ | ++ | - | ++ | + | - | ++++ | - |
| | 3 | Healthy | - | ++++ | + | ++ | - | - | - | ++ | - |
| P072 | 1 | Healthy | ++ | - | - | + | + | - | - | ++++ | - |
| | 2 | Healthy | - | - | + | + | - | - | - | +++ | - |
| | 3 | Healthy | - | - | - | - | - | - | + | - | - |
| P073 | 1 | Healthy | - | - | - | - | - | - | - | + | - |
| P074 | 1 | Healthy | - | - | + | - | - | + | - | - | - |
| | 2 | Healthy | - | + | + | - | - | ++ | - | ++++ | + |
| | 3 | Healthy | - | - | ++ | ++ | - | - | + | +++ | - |
| P075 | 1 | Healthy | + | - | - | - | + | + | + | + | - |
| P076 | 1 | SD | - | - | + | + | - | - | - | +++ | - |
